# A general LC/MS-based RNA sequencing method for direct analysis of multiple-base modifications in RNA mixtures

**DOI:** 10.1101/643387

**Authors:** Ning Zhang, Shundi Shi, Tony Z. Jia, Ashley Ziegler, Barney Yoo, Xiaohong Yuan, Wenjia Li, Shenglong Zhang

## Abstract

A complete understanding of the structural and functional potential of RNA requires understanding of chemical modifications and noncanonical bases; this in turn requires advances in current sequencing methods to be able to sequence not only canonical ribonucleotides, but at the same time directly sequence these nonstandard moieties. Here, we present the first direct and modification type-independent RNA sequencing method *via* integration of a hydrophobic end-labeling strategy with of 2-D mass-retention time LC/MS analysis to allow *de novo* sequencing of RNA mixtures and enhance sample usage efficiency. Our method can directly read out the complete sequence, while identifying, locating, and quantifying base modifications accurately in both single and mixed RNA samples containing multiple different modifications at single-base resolution. Our method can also quantify stoichiometry/percentage of modified RNA *vs*. its canonical counterpart RNA, simulating a real biological sample where modifications exist but may not be 100% at a particular site of the RNA. This method is a critical step towards fully sequencing real complex cellular RNA samples of any type and containing any modification types and can also be used in the quality control of modified therapeutic RNAs.

## INTRODUCTION

RNAs deliver a diverse spectrum of biological functions in nature not only through sequences of the four canonical nucleosides, but also through hundreds of types of structural modifications, both known and unknown. Aberrant RNA modifications, such as methylations and pseudouridinylations, have been correlated with major human diseases such as cancers (1–3), type-2 diabetes (4,5), obesity (6,7), and neurological disorders (8,9). Despite their significance, there is no method available efficient or effective enough to determine sequences of highly modified RNA with different chemical modifications, and thus we only understand the function of a handful of the >160 identified RNA modifications. Knowing the correct sequences of therapeutic oligoribonucleotides containing modified bases is also a prerequisite for their own quality control, and without a widely-available accurate sequencing method for noncanonical oligoribonucleotides, most therapeutic oligoribonucleotides containing modifications have been used clinically without direct sequence determination (10).

The primary challenge in structural and functional elucidation of RNA modifications in biological samples is that these chemical modifications are typically of low abundance relative to unmodified nucleobases and, subsequently, are often undetectable using conventional methods including Next-Generation Sequencing (NGS). To overcome the low sample-amount problem, methods for studying the transcriptome often require complementary DNA (cDNA) synthesis followed by PCR (11–13). However, this results in analytes that contain only information of canonical nucleobases, and therefore, nucleobase modification information is permanently lost in these indirect sequencing methods. Other base-specific indirect NGS-based RNA sequencing techniques (14–17) are typically tailored to only one specific modification and cannot report information regarding any other modifications, known or unknown, that coexist in the same sample. Even with the recent development of novel specific detecting chemistries/antibodies, the list of NGS-detectable RNA modifications is still extremely limited—far behind what is needed to detect all >160 RNA modifications to properly elucidate their structures and functions.

As opposed to indirect NGS sequencing methods, direct sequencing of RNA molecules without the need for cDNA synthesis or PCR would theoretically allow direct analysis of RNA sequences including all associated modified nucleotides. However, some methods rely on reading DNA bases that are added to the RNA template (the cDNA), instead of the RNA template itself, and suffer from the same problems as sequencing-by-synthesis techniques (18). Nanopore RNA-Seq has detected modifications like m^6^A (19) and m^7^G (20) in RNA, but it relies on distinct electronic signatures to sequence each modification, and the system must be trained with sequences containing known modifications (19,21), severely limiting its discovery application potential. There are other methods for detecting RNA modifications that do not involve cDNA, but they usually employ complete enzymatic or chemical hydrolysis (22), which annihilates simultaneous location and sequence information.

In contrast with modification-specific methods, mass spectrometry (MS)-based approaches are theoretically applicable to all modifications in general, as they take advantage of the fact that most nucleobase modifications either inherently have different masses themselves or can be easily converted into different masses, which result in their use as unique natural/artificial mass tags for sequencing (23). These methods, especially liquid chromatography (LC)/MS, have long been used for identifying known and unknown modifications (24–27) as well as RNA modification mapping (28–31). However, without mass laddering, accurate and *de novo* RNA sequencing has not been possible because the location and sequence information of RNA modifications in the sample strands is lost. These methods typically rely on complete and uncontrolled degradation of RNA into single nucleotides or smaller fragments (22,32); even when the degradation is controlled, for example *via* enzymatic cleavage (24,33), a ladder suitable for sequencing is not generated. Direct top-down MS and tandem MS have been used for characterizing some RNA modifications(30,31,34–38) and for sequence determination (39–41). Other traditional MS-based RNA sequencing methods have also been used, and these have been discussed and reviewed at length (32,39,42–45). However, these traditional MS-based methods are still not widely applicable for direct and *de novo* sequencing of RNA due in part to significant methodological inadequacies in the preparation of high-quality sequence ladders, which must be unaffected by sequence content/context and must also produce the complete suite of mass-ladders required for accurate sequencing basecalling (32,46). Even if complete and sequence-independent ladders can be produced, it still is difficult to differentiate one mass ladder from other overlapping ladders, *e.g*., due to multiple possible fragmentation sites (26,32,47), and it remains challenging in identifying these mass ladder components for sequencing due to signal/noise issues; MS peaks arising from noise often overlap with the desired ladder components and complicate data analysis. As such, to this date, MS-sequencing techniques could mainly be applied towards quality control methods serving only as a tool for sequence confirmation of RNA with known sequences (10,32,42,48), with very limited applications as a *de novo* RNA sequencing method.

One way to circumvent the above issues is to perform enzymatic or chemical degradation of RNA so that well-defined sequencing ladders can be formed before introduction into the mass spectrometer (49–54). Ideally, ladder cleavage must be highly uniform with exactly one random and unbiased cut on each RNA strand (46). However, the uniformity of ladder sequences generated by current techniques is still unsatisfactory, often mixed with undesired fragments from multiple cuts on each RNA strand and subsequent metal adduct formation, complicating downstream data analysis. Therefore, even with generation of all necessary ladder components prior to injection into the MS, based on conventional one-dimensional (1-D) MS data, it is still not trivial to sequence even a purified single-stranded RNA due to difficulty in identification of all ladder components required for sequencing (50). Samples containing multiple sequences would be even more challenging, if not impossible, due to generation of a large number of overlapping mass ladders, significantly complexicating downstream data analysis.

Therefore, we previously established a two-dimensional (2-D) LC/MS-based RNA sequencing method to assist in ladder identification by taking advantage of predictable regularities in high performance liquid chromatography (HPLC) separation of formic acid-degraded RNA digests to produce optimized mass-retention time (t_R_) ladders (46). LC/MS in particular is the optimal tool for direct RNA sequencing, as the two primary variables that factor into RNA sequencing are the mass and t_R_ of the various ladder compounds, both of which are endogenous values unique to each ladder component and which complement each other nicely. Typically, smaller compounds are eluted from the LC column first with a smaller t_R_, followed by compounds with increasing mass and longer t_R_. An ideal set of RNA ladders would be visualized in a mass *vs*. t_R_ graph as a sigmoidal curve, with 2-D mass-t_R_ ladders now becoming more easily visually identifiable than a 1-D mass ladder due to the increase in the dimensionality of the data. By calculating the mass differences between the desired constituent sequence fragments of each ladder in the curve, the sequence of the RNA, and the identity, quantity, and location of mass-altering modifications, can be unambiguously determined.

However, the previously developed 2-D LC/MS-based RNA sequencing method requires paired-end reads for reading terminal nucleobases, and therefore cannot read a complete sequence from one single ladder. Also, the 2-D method still experiences difficulties in MS sequencing of multiple RNA strands mainly due to the complexity of LC/MS data analysis, which includes the adjacent existence of two ladders (5’ and 3’), which may cause confusion as to which fragment belongs to which ladder, becoming much more complicated with multiple RNA strands. To unleash the MS-based sequencing from its current restrains for its much broader applications, two essential issues have to be addressed: 1) how to generate a complete suite of well-defined ladder fragments that allows reading from the first ribonucleotide to the final one and 2) how to find easily each of the complete set of ladders in complicated MS data. Here we present an innovative solution to address these two critical issues using a hydrophobic end-labeling strategy *via* introducing 2-D mass-t_R_ shifts for ladder identification. Specifically, we added mass-t_R_ labels on the 5’ and/or 3’ end, and at least one of these moieties results in a retention time shift to longer times, causing all of the 5’ and/or 3’ ladder fragments to have a markedly delayed t_R_, which clearly distinguished the 5’ ladder from the 3’ ladder. The hydrophobic label tags not only result in mass-t_R_ shifts of labeled ladders, making it much easier to identify each of the 2-D mass ladders needed for MS sequencing of RNA and thus simplifying base-calling procedures, but labeled tags also inherently increase the masses of the RNA ladder fragments so that the terminal bases can even be identified, thus allowing the complete reading of a sequence from one single ladder, rather than requiring paired-end reads. We test the efficacy of the new strategy on a series of synthetic RNA oligonucleotides of varying lengths containing both canonical and modified bases as a proof-of-concept study. We were able to sequence RNA containing both pseudouridine (ψ) and 5-methylcytosine (m^5^C) simultaneously at single-base resolution and quantify the stoichiometry/percentage of the RNA containing the m^5^C modified base *vs*. its canonical counterpart RNA accordingly as an analog to a real system in which the modification efficiency of a given base is not 100%. Together with our end-labeling strategy, we were able to identify, locate, and quantify these multiple base modifications while accurately sequencing the complete RNA not only in a single purified RNA strand, but also in sample mixtures containing 12 distinct sequences of RNAs.

## MATERIAL AND METHODS

### Chemical materials

The following RNA oligonucleotides were obtained from Integrated DNA Technologies (Coralville, IA, USA) and used without further purification.

RNA #1: 5’-HO-CGCAUCUGACUGACCAAAA-OH-3’

RNA #2: 5’-HO-AUAGCCCAGUCAGUCUACGC-OH-3’

RNA #3: 5’-HO-AAACCGUUACCAUUACUGAG-OH-3’

RNA #4: 5’-HO-UGUAAACAUCCUACACUCUC-OH-3’

RNA #5: 5’-HO-UAUUCAAGUUACACUCAAGA-OH-3’

RNA #6: 5’-HO-GCGUACAUCUUCCCCUUUAU-OH-3’

RNA #7: 5’-HO-CGCCAUGUGAUCCCGGACCG-OH-3’

RNA #8: 5’-HO-ACACUGACAUGGACUGAAUA-OH-3’

RNA #9: 5’-HO-GCGGAUUUAGCUCAGUUGGG-OH-3’

RNA #10: 5’-HO-CACAAAUUCGGUUCUACAAG-OH-3’

RNA #11: 5’-HO-GCGGAUUUAGCUCAGUUGGGA-OH-3’

RNA #12: 5’-HO-AAACCGUψACCAUUAm^5^CUGAG-OH-3’

RNA #13: 5’-HO-AAACCGUψACCAUUACψGAG-OH-3’

RNA #14: 5’-HO-AAACCGUUACCAUUAm^5^CUGAG-OH-3’

Formic acid (98-100%) was purchased from Merck (Darmstadt, Germany). Biotinylated cytidine bisphosphate (pCp-biotin), {Phos (H)}C{BioBB}, was obtained from TriLink BioTechnologies (San Diego, CA, USA). Adenosine-5’-5’-diphosphate-{5’-(cytidine-2’-*O*-methyl-3’-phosphate-TEG}-biotin, A(5’)pp(5’)Cp-TEG-biotin-3’, was synthesized by ChemGenes (Wilmington, MA, USA). T4 DNA ligase 1, T4 DNA ligase buffer (10 ×), the adenylation kit including reaction buffer (10 ×), 1 mM ATP, and *Mth* RNA ligase were obtained from New England Biolabs (Ipswich, MA, USA). ATPγS and T4 polynucleotide kinase (3’-phosphatase free) were obtained from Sigma-Aldrich (St. Louis, Missouri, USA). Biotin maleimide was purchased from Vector Laboratories (Burlingame, CA, USA). Cyanine3 maleimide (Cy3) and sulfonated Cyanine3 maleimide (sulfo-Cy3) were obtained from Lumiprobe (Hunt Valley, Maryland, USA). The streptavidin magnetic beads were obtained from Thermo Fisher Scientific (Waltham, MA, USA). Chemicals needed for conversion of pseudouridine including CMC (*N*-cyclohexyl-*N*’-(2-morpholinoethyl)-carbodiimide metho-*p*-toluenesulfonate), bicine, urea, EDTA and Na_2_CO_3_ buffer, were obtained from Sigma-Aldrich (St. Louis, MO, USA).

### Workflow

(1) Chemical conversion of pseudouridine was applied for distinguishing pseudouridine from uridine. (2) Labels were added on one or both ends of RNA strands with optimized experimental procedures. (3) The single RNA strand or mixtures of RNA strands was/were degraded into a series of short, well-defined fragments (sequence ladder), ideally by random, sequence context-independent, and single-cut cleavage of phosphodiester bonds on each RNA strand over its entire length, through a 2’-OH-assisted acidic hydrolysis mechanism. (4) If needed, physical separation of biotinylated RNA from unlabeled RNA was performed using streptavidin-coated magnetic beads. (5) The digested fragments were then subjected to LC/MS analysis and the deconvoluted masses and t_R_ were analyzed to identify each ladder fragment. (6) Algorithms were applied to automate the data processing and sequence generation process.

### 3’-End labeling method

Two-step protocol: (1) Adenylation: We set up the following reaction with a total reaction volume of 10 μL in an RNAse-free, thin walled 0.5 mL PCR tube: 1 × adenylation reaction buffer (5’-adenylation kit), 100 μM of ATP, 5.0 μM of *Mth* RNA ligase, 10.0 μM pCp-biotin, and nuclease-free, deionized water (Thermo Fisher Scientific, USA). The reaction was incubated in a GeneAmp™ PCR System 9700 (Thermo Fisher Scientific, USA) at 65°C for 1 hour followed by the inactivation of the enzyme *Mth* RNA ligase at 85°C for 5 min. (2) Ligation: A 30 μL reaction solution contained 10 μL of reaction solution from the adenylation step, 1 × reaction buffer, 5 μM target RNA sample, 10% (v/v) DMSO (anhydrous dimethyl sulfoxide, 99.9%, Sigma-Aldrich, USA), T4 RNA ligase (10 units), and nuclease-free, deionized water. The reaction was incubated overnight at 16°C, followed by column purification.

One-step protocol: A(5’)pp(5’)Cp-TEG-biotin-3’ was applied to improve the labeling efficiency by eliminating the adenylation step, while simplify the labeling method. The ligation step was achieved by a 30 μL reaction solution containing 1× reaction buffer, 5 μM target RNA sample, 10 μM A(5’)pp(5’)Cp-TEG-biotin-3’, 10% (v/v) DMSO, T4 RNA ligase (10 units), and nuclease-free, deionized water. The reaction was incubated overnight at 16°C, followed by column purification. Oligo Clean & Concentrator (Zymo Research, Irvine, CA, U.S.A.) was used to remove enzymes, free biotin, and short oligonucleotides.

### 5’-End labeling method

Biotin labeling at the 5’-end required two steps. In an RNase-free, thin walled PCR tube (0.5 mL) containing 10 × reaction buffer, 90 μM of RNA, 1 mM of ATPγS, and 10 units of T4 polynucleotide kinase, the total reaction volume was diluted to 10 μL with nuclease-free, deionized water; incubation was then carried out for 30 min at 37 °C. We then added 5 μL of biotin maleimide that was dissolved in 312 μL anhydrous DMF (anhydrous dimethyl sulfoxide, 99.9%, Sigma-Aldrich, USA), mixed the sample by vortexing, and incubated the sample for 30 min at 65 °C. Column purification using Oligo Clean & Concentrator was performed as described above.

Different tags, such as a hydrophobic Cy3 (cyanine 3) or Cy5 (cyanine 5), were introduced to the 5’-end by the same method as above (except through Cy3-maleimide or sulfo-Cy3 maleimide replacement of the biotin maleimide), to distinguish its ladder from the 3’-biotinylated ladder. The optimization of the reaction conditions, compared to the above described 2-step protocol, was performed to obtain high labeling efficiency in the following manner: 1) sulfo-Cy3 was used for obtaining high water solubility of the dye with a molar ratio of reactants at 50:1 (sulfo-Cy3 to RNA); 2) the pH of the reaction solution was adjusted to 7.5 by Tris-HCl buffer (1 M) with a final concentration of 50 mM; and 3) the reaction time was lengthened to overnight (16 hrs) with constant stirring.

### Acid hydrolysis degradation

Unless otherwise indicated, formic acid was applied to degrade full length RNA samples for producing mass ladders (46,50). We divided each RNA sample solution into three equal aliquots for formic acid degradation using 50% (v/v) formic acid at 40 °C, with one reaction running for 2 min, one for 5 min, and one for 15 min. For the experiments regarding generation of internal fragments (Figure S4), a 60 min formic acid treatment was performed on RNA #3. The reaction mixture was immediately frozen on dry ice followed by lyophilization to dryness, which was typically completed within 30 min. The dried samples were combined and suspended in 20 μL nuclease-free, deionized water for LC/MS measurement. In Figure 6 we started with two separate samples of the same 11 sequences (RNA #1 – RNA #11), one with a 3’-biotin-label and one with a 5’-sulfo-Cy3 label, and mixed these samples along with a sample containing 3’-biotin-labeled RNA #12 before injection into the LC/MS.

### Biotin/streptavidin capture/release step

For the sample in Figure 1b (no other samples required this step), 200 μL of Dynabeads™ MyOne™ Streptavidin C1 beads were activated by first adding an equal volume of 1 × B&W buffer. This solution was vortexed and placed on the magnet-stand for 2 min, followed by the discarding of the supernatant. We washed the beads twice with 200 μL of Solution A (DEPC-treated 0.1 M NaOH and DEPC-treated 0.05 M NaCl) and once in 200 μL of Solution B (DEPC-treated 0.1 M NaCl). A final addition of 100 μL of 2 × B&W buffer brought the concentration of the beads to 20 mg/mL. We then added an equal volume of biotinylated RNA in 1 × B&W buffer, incubated the sample for 15 min at room temperature using gentle rotation, placed the tube on the magnet for 2 min, and discarded the supernatant. The coated beads were washed 3 times in 1 × B&W buffer and we measured the final concentration of each wash step supernatant by Nanodrop for recovery analysis, to confirm that the target RNA molecules remained on the beads. For releasing the immobilized biotinylated RNAs, we incubated the beads in 10 mM EDTA (Thermo Fisher Scientific, USA), pH 8.2 with 95% formamide (Thermo Fisher Scientific, Waltham, MA, USA) at 65°C for 5 min. Finally, the sample tube was placed on the magnet-stand for 2 min and the supernatant (containing the target RNA molecules) was collected by pipet.

**Figure 1.**
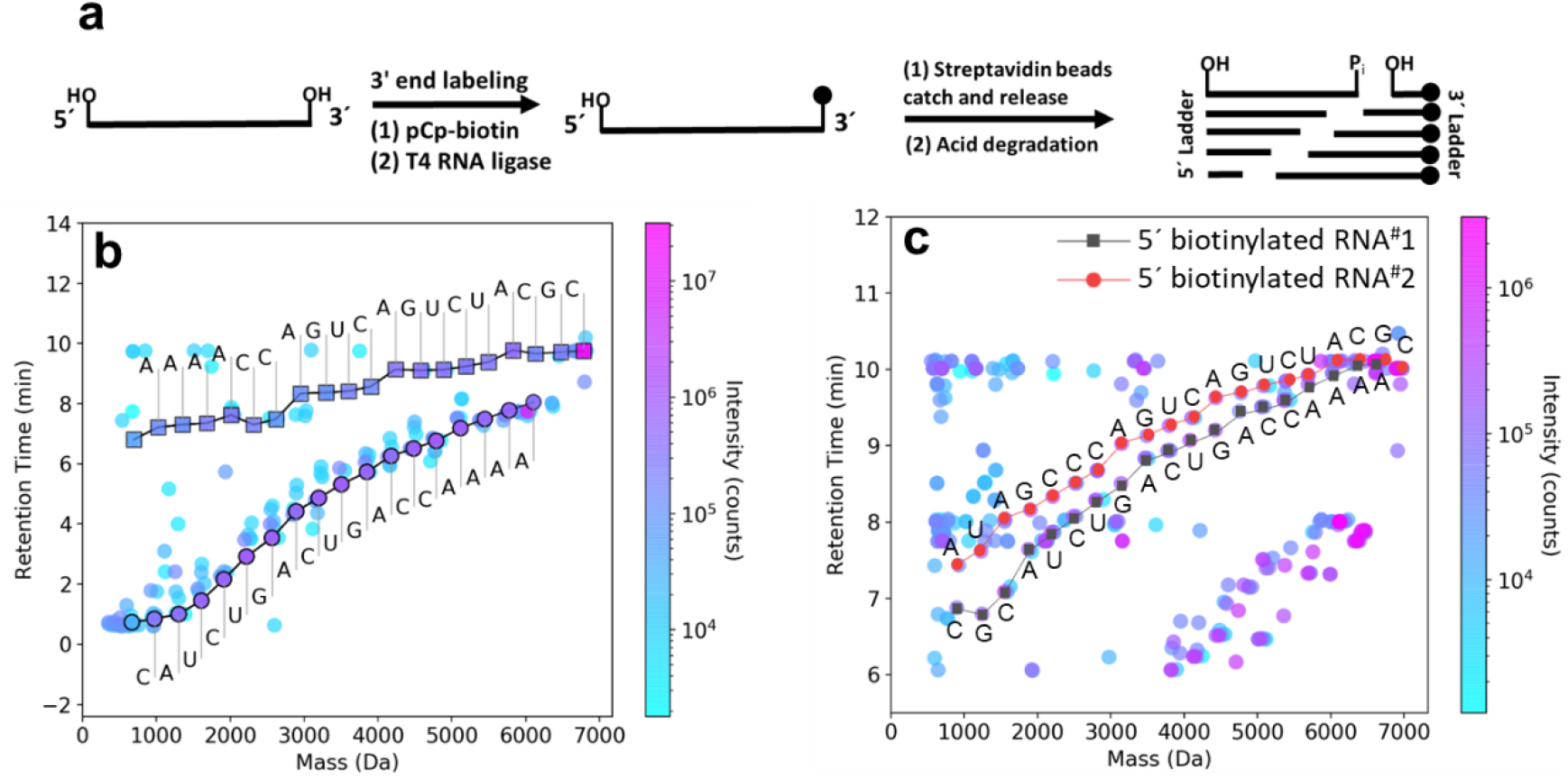
**a)** Method for introducing a biotin label to the 3’-end of RNA. **b)** Separation of the 3’-ladder from the 5’-ladder and other undesired fragments on a mass-retention time (tR)-plot based on systematic changes in t_R_ of 3’-biotin-labeled mass-t_R_ ladders of RNA #1 (19 nt). The sequences were *de novo* generated automatically by an algorithm described in the SI. **c)** Simultaneous sequencing of two RNAs of different lengths (RNA #1 and RNA #2; 19 nt and 20 nt, respectively) after 5’-biotin labeling. The sequences presented were manually acquired based on the mass-t_R_ ladders identified from the algorithm-processed data.

### Chemistry for differentiating pseudouridine from uridine

The experiments to convert pseudouridine into CMC-ψ adducts were performed according to a reported method (55). Each RNA sample (1 nmol) was treated with 0.17 M CMC in 50 mM Bicine, pH 8.3, 4 mM EDTA, and 7 M urea at 37 °C for 20 min in a total reaction volume of 90 μL. The reaction was stopped with 60 μL of 1.5 M sodium acetate (NaOAc) and 0.5 mM EDTA, pH 5.6 (buffer A). After purification using an Oligo Clean & Concentrator, 60 μL of 0.1 M Na_2_CO_3_ buffer, pH 10.4 was added into the solution, the solution was brought to a reaction volume of 120 μL by addition of nuclease-free, deionized water, and incubated at 37 °C for 2 h. Reaction was stopped with buffer A and purified by Oligo Clean & Concentrator.

### Experimental protocol for quantifying percentage of RNA modifications

RNA #14 (m^5^C modified RNA) and RNA #3 (non-modified RNA) were mixed with percentages of m^5^C modified RNA in the mixed samples were 10%, 20%, 30%, 40%, 50% and 100%, respectively. One-step protocol was applied for labeling their 3’ ends with biotin. The ligation step was achieved by a 30 μL reaction solution containing 1× reaction buffer, 5 μM mixed RNA sample, 10 μM A(5’)pp(5’)Cp-TEG-biotin-3’, 10% (v/v) DMSO, T4 RNA ligase (10 units), and nuclease-free, deionized water. The reaction was incubated overnight at 16°C, followed by column purification. Oligo Clean & Concentrator (Zymo Research, Irvine, CA, U.S.A.) was used to remove enzymes, free biotin, and short oligonucleotides. Formic acid was applied to degrade full length RNA samples for producing mass ladders. The reaction mixture was immediately frozen on dry ice followed by lyophilization to dryness within 30 min. The relative quantities of different product species were quantified by integrating the extracted ion current (EIC) peaks of 3’-biotin labeled methylated RNA at m/z 722.7118 and non-modified RNA at m/z 721.3105, respectively, at a charge state of −10 (56).

### LC-MS Analysis

Samples were separated and analyzed on a 6550 Q-TOF mass spectrometer coupled to a 1290 Infinity LC system equipped with a MicroAS autosampler and Surveyor MS Pump Plus HPLC system (Agilent Technologies, Santa Clara, CA, USA) (Hunter College Mass Spectrometry, NY, USA). All separations were performed reversed-phase HPLC using an aqueous mobile phase (A), 25 mM hexafluoro-2-propanol (HFIP) (Thermo Fisher Scientific, USA) with 10 mM diisopropylamine (DIPA) (Thermo Fisher Scientific, USA) at pH 9.0, and an organic mobile phase (B), methanol, across a 50 mm × 2.1 mm Xbridge C18 column with a particle size of 1.7 μm (Waters, Milford, MA, USA). The flow rate was 0.3 mL/min, and all separations were performed with the column temperature maintained at 35 °C. Injection volumes were 20 μL, and sample amounts were 15-400 pmol of RNA. Data were recorded in negative ion mode. The sample data were acquired using the MassHunter Acquisition software (Agilent Technologies, USA). To extract relevant spectral and chromatographic information from the LC-MS experiments, we used the Molecular Feature Extraction (MFE) workflow in MassHunter Qualitative Analysis (Agilent Technologies, USA). This proprietary molecular feature extractor algorithm performs untargeted feature finding in the mass and retention time dimensions. In principal, any software capable of compound identification could be used. The software settings were varied depending on the amount of RNA used in the experiment. In general, we wanted to include as many identified compounds as possible, up to a maximum of 1000. Details about data extraction can be found in *Supplementary Materials*.

In addition to automating the sequence generation, we also manually searched for the mass ladders using the MFE workflow in MassHunter Qualitative Analysis in order to confirm the accuracy of automating sequencing. In Tables S1–S40, we provide the theoretical mass of each fragment (obtained by ChemDraw), base mass, base name, observed mass, t_R_, volume (peak intensity), quality score, and ppm mass difference.

All figures presented are representative data of multiple experimental trials (n≥3). For ease of visualization, the 5’-sulfo-Cy3 labeled mass ladders and the 3’-biotinylated mass ladders were plotted separately (*i.e*., 3’-biotinylated mass ladders were all plotted in Figure 6a and the 5’-sulfo-Cy3 labeled mass ladders were all plotted in Figure 6b). Then, for each sequence curve (up to 12 on a given plot), the starting t_R_ values were normalized to start at 4 min intervals (except in the case of RNA #12 in Figure 6a, where we used an 8-min interval gap). The absolute differences between the starting t_R_ value and subsequent t_R_ values of any single given curve remain unchanged; only the visual “height” at which each curve is plotted was changed. Plots for Figure 6 were produced with OriginLab. In all figures except Figures 6a and 6b, the mass-t_R_ plot was generated without normalization of any of the t_R_ values. Because of a missing base assignment in the original sample, we combined two samples and analyzed and visualized the combined data in Figure 2b. One sample contained RNA #1 with both 5’-Cy3 and 3’-biotin labels, while the second combined sample contained RNA #1 with only a 5’-Cy3 label (Table S6).

**Figure 2.**
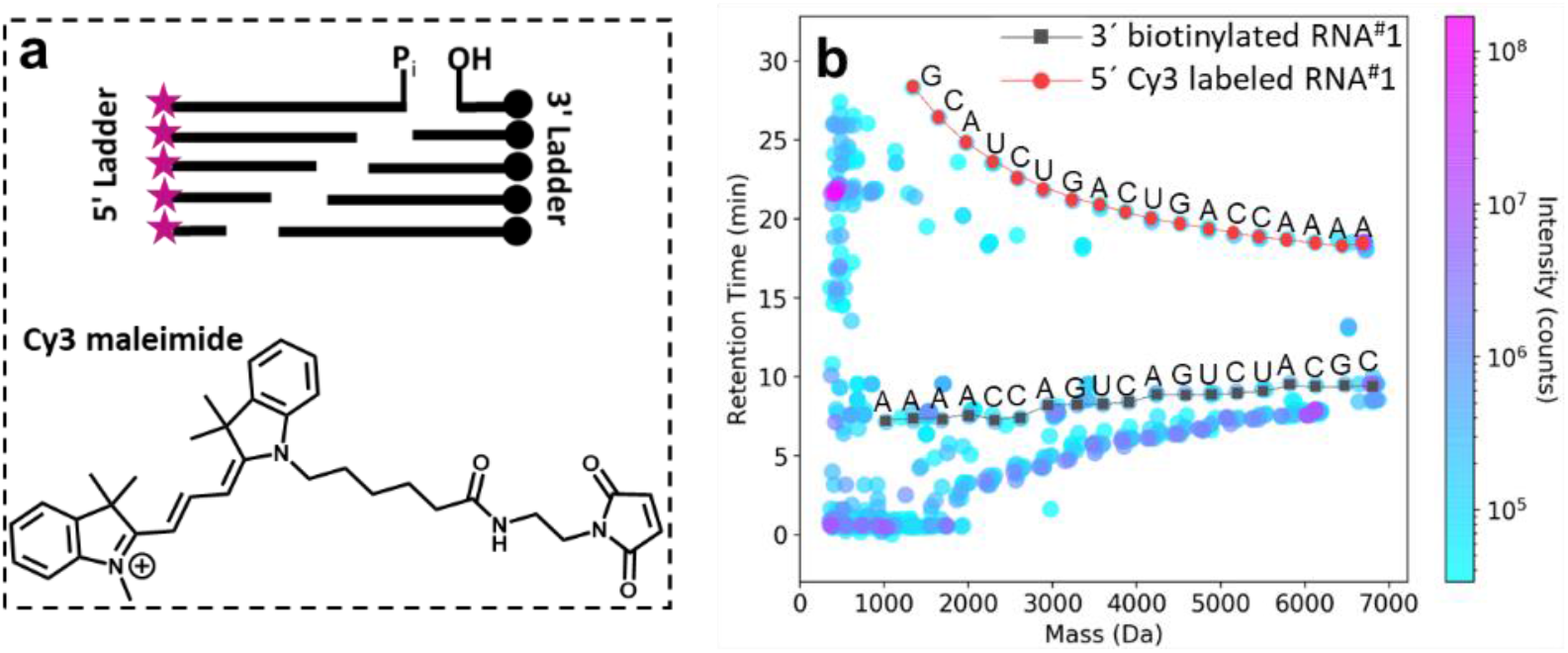
**a)** General strategy to differentiate two series of ladder fragments (5’ *vs*. 3’) from each other by introducing a hydrophobic cyanine 3 (Cy3) to the 5’-end and biotin to the 3’-end, respectively, of any RNA. **b)** Mass-t_R_ plot of a sample containing all the ladder fragments needed for sequencing from 5’-Cy3-labeled and 3’-biotin-labeled RNA #1. Differentiation of the ladders can occur due to significant changes in the t_R_ s afforded by the two tags. The sequence was manually read from both mass-t_R_ ladders identified from the algorithm-processed data.

### Automated RNA sequencing and visualization algorithm

The first step of the LC/MS data analysis is to perform data pre-processing and reduction so that the LC/MS data will become less noisy, and consequently easier to read out the RNA sequence(s) from the data in the next step. From the multi-dimensional LC/MS data, there are several dimensions that can be used to pre-process the data and reduce its volume, such as Retention Time (tR), Intensity (Volume), and Quality Score (QS). Please see *Supplementary Materials* for details on data processing and modifications to the sequencing algorithm. The source codes of the revised algorithms are available upon request.

## RESULTS AND DISCUSSION

### Generation of labeled RNA degraded ladder fragments for mass analysis

In our new experimental approaches, we either label one RNA end and leave the other end unlabeled, or label the two ends of the RNA with different tags to better distinguish them in our 2-D LC/MS method. In one labeling strategy, we introduced a biotin tag to either the 3’-end or the 5’-end of the RNA prior to LC/MS analysis in order to introduce an t_R_ and mass shift to exactly one mass ladder (46). This method can help simplify LC/MS data analysis and prevent confusion as to which fragment belongs to which ladder when sequencing mixed RNA samples. It increases the masses of RNA ladders so that the terminal bases can be identified, avoiding messy low mass regions where it is difficult to differentiate mononucleotides and dinucleotides from multi-cut internal fragments; improves sequencing accuracy by reading a complete sequence from one single ladder, rather than requiring paired-end reads; simplifies base-calling procedures, making it easier for the ladder components to be identified due to selective t_R_ shifts; and improves sample efficiency by allowing for longer degradation time points (15 min up to 60 min) than reported before (5 min) (46). These improvements can help reduce the minimum RNA sample loading requirement as compared to the first-generation method, increasing the potential to sequence endogenous RNA samples with rare RNA modifications.

For labeling RNAs at their 3’-ends (Figure 1a), we first activated biotinylated cytidine bisphosphate (pCp-biotin) by adenylation using ATP and *Mth* RNA ligase to produce AppCp-biotin. Then, the RNAs with a free 3’-terminal hydroxyl were ligated to the activated AppCp-biotin *via* T4 RNA ligase. Streptavidin-coupled beads were used to isolate the 3’-biotin-labeled RNAs, which were released for acid degradation and subsequent LC/MS analysis after breaking the biotin-streptavidin interaction. Biotin can also be labeled at the 5’-end (*MATERIAL AND METHODS*).

As an example to test this strategy, short RNA oligonucleotides (19 nt and 20 nt RNA: RNA #1 and RNA #2, respectively) were designed and synthesized as model RNA oligonucleotides for individual and group tests, but other RNA with different lengths could be tested in the same way. We first subjected 3’-biotin-labeled RNA #1 to physical separation by streptavidin bead capture and release. In Figure 1b, subsequent separation using t_R_ shifts of a 3’-biotin-labeled mass ladder from an unlabeled 5’-ladder of RNA #1 avoids confusion as to which fragment belongs to which ladder, and the isolated curve in the output is much simpler to analyze than the two adjacent curves of the first-generation method. The *de novo* sequencing process was performed by a modified version of a published algorithm (*Supplementary Materials*) (46). This algorithm uses hierarchical clustering of mass adducts to augment compound intensity. Co-eluting neutral and charge-carrying adducts were recursively clustered, such that their integrated intensities were combined with that of the main peak. This increased the intensity of ladder fragment compounds and reduced the data complexity in the regions critical for generating sequencing reads.

In Figure 1b, the 3’-ladder curve is shifted up (with respect to the y-axis) because the biotin label causes an increase in t_R_, and the complete sequence of RNA #1 can be read from the top curve alone. Similarly, the complete RNA #1 reverse sequence can be read from the unlabeled 5’-ladder curve (which does not have a shift in t_R_) directly, with the exception of the first nucleotide as it was below the detection threshold applied in the LC/MS. Without this strategy, end pairing would be required to read out the complete sequence, as reported before (46). With this advance, each RNA can be read out completely from one curve, and we were able to sequence mixed samples containing multiple RNAs each labeled with a 5’-biotin label (Figure 1c). The separation of the 3’- and 5’-ladders for each sample significantly reduces the complexity of the resultant LC/MS data so that it is much easier than the previous method (46) to find complete sets of ladder components needed for sequencing, and thus reduce the complexity of the base-calling procedures.

Thanks to this end labeling, we can read out both complete sequences in a mixture of two RNAs, one 19 nt (RNA #1) and one 20 nt (RNA #2), from exactly one curve per RNA strand. In the case of this sample, we used the algorithm to perform crucial mass adduct clustering in order to further simplify the data for finding the complete sets of mass ladder components needed for sequencing. Form the sigmoidal curves consisting of all mass ladder components in the simplified 2-D mass-t_R_ plot, we then were able to manually determine the sequences of the sample RNA strands simply by calculating the mass differences of two adjacent ladder components (Figure 1c). Although our samples were all synthetic samples and we did not necessarily need to use biotin-streptavidin binding-cleavage to physically separate our sample of interest from other RNA strands (we only actually required the t_R_ shift associated with biotin-labeling), incorporation of the biotin label also provides the possibility of physical separation of specific samples that could be useful for sequencing real biological samples.

In order to further increase the observed t_R_ shift afforded by end-labeling, an RNA sample may be labeled with other bulky moieties such as a hydrophobic cyanine 3 (Cy3) or cyanine 5 (Cy5), to magnify their t_R_ difference. We introduced different tags, such as Cy3, which is bulky and can cause a greater t_R_ shift than biotin (46) at the 5’-end of the original RNA strand to be sequenced; a biotin moiety was introduced to the 3’-end of the RNA as described before. These end labels should systematically affect the t_R_ s of all 5’- and 3’-ladder fragments so as to differentiate the two ladder curves for sequencing, which was confirmed by *in silico* studies (Figures S1a and S1b). As shown in Figure 2a, a Cy3 tag was added *via* a two-step reaction at the 5’-end of the RNA sample. Similar to the 5’-biotinylation methodology, after thiolphosphorylation at the first step, Cy3 maleimide was conjugated to RNA. After acid degradation of the double end-labeled RNAs, the resulting fragments were directly subjected to LC/MS without any affinity-based physical separation. Our preliminary data showed that in the 2-D mass-t_R_ plot, the 5’-Cy3-labeled ladder fragments form a curve further away from the 5’-biotin-labeled ladder (Figure 2b), as more hydrophobic tags elicit larger t_R_ shifts. In fact, the t_R_ trend for the Cy3-labeled 5’-ladder changes direction, as in the mass-t_R_ plot, the sequence curve goes down in t_R_ with increasing mass due to the hydrophobic nature of the Cy3 moiety, as compared to the biotin-labeled 3’-ladder, which increases in t_R_ with increasing mass (as also observed in all previous biotin-labeled and unmodified mass ladder samples). This results in two curves that are more separable/distinguishable during the 2-D analysis, making it easier to base call the sequences of the ladders even without physical separation. With bidirectional sequencing, the method’s read length can be doubled, and its accuracy can be improved significantly by reading a complete sequence from both the 3’- and 5’-ladders.

### RNA labeling efficiency

Despite various reported RNA labeling methods, including those reported above, it remains a challenge to introduce tags such as biotin or fluorescent dyes onto RNA with high yield. However, labeling both ends of RNA with these selected tags is an essential step of our direct RNA sequencing method. The labeling efficiency directly results in how much RNA sample can be used to generate MS signals, with a higher labeling efficiency leading to a reduced sample requirement. To increase the labeling efficiency, we explored new labeling strategies and were able to achieve high labeling efficiency at both the 5’- and 3’-end (Figure 3). For the 5’-end label, the labeling efficiency of full-length RNA was improved from ~60% (a yield obtained when labeling RNA #1 in Figure 2b) to ~90% (Figure 3a) by using a modified reaction protocol, including 1) using sulfo-Cy3 (Figure 3c) instead of Cy3 to increase aqueous solubility of the tag, 2) adjusting the pH of the solution to 7.5, and 3) lengthening the reaction time while maintaining constant stirring (*MATERIAL AND METHODS*). We can see even after acid degradation of a 5’-sulfo-Cy3-labeled RNA #1 that the labeled ladder components far outnumber the unlabeled ladder components with respect to absolute intensity, as the unlabeled fragments do not appear on the plot after mild filtering (Figure S2), resulting in better sample usage and reduction of MS data complexity. For better labeling efficiency at the 3’-end, we synthesized A(5’)pp(5’)Cp-TEG-biotin-3’ (Figure 3c), an active form of biotinylated pCp, which eliminates the adenylation step previously required for biotinylation (57). A highly quantitative yield (~95%) for 3’-end labeling was observed (Figure 3b) when labeling a 21 nt RNA (RNA #11) using this method. By incorporating both optimized end-labeling strategies into the sample preparation protocol, not only is the minimum sample loading amount requirement is now less of a hindrance to the overall sequencing workflow, but the complexity of the data required for downstream sequencing may simultaneously be reduced.

**Figure 3.**
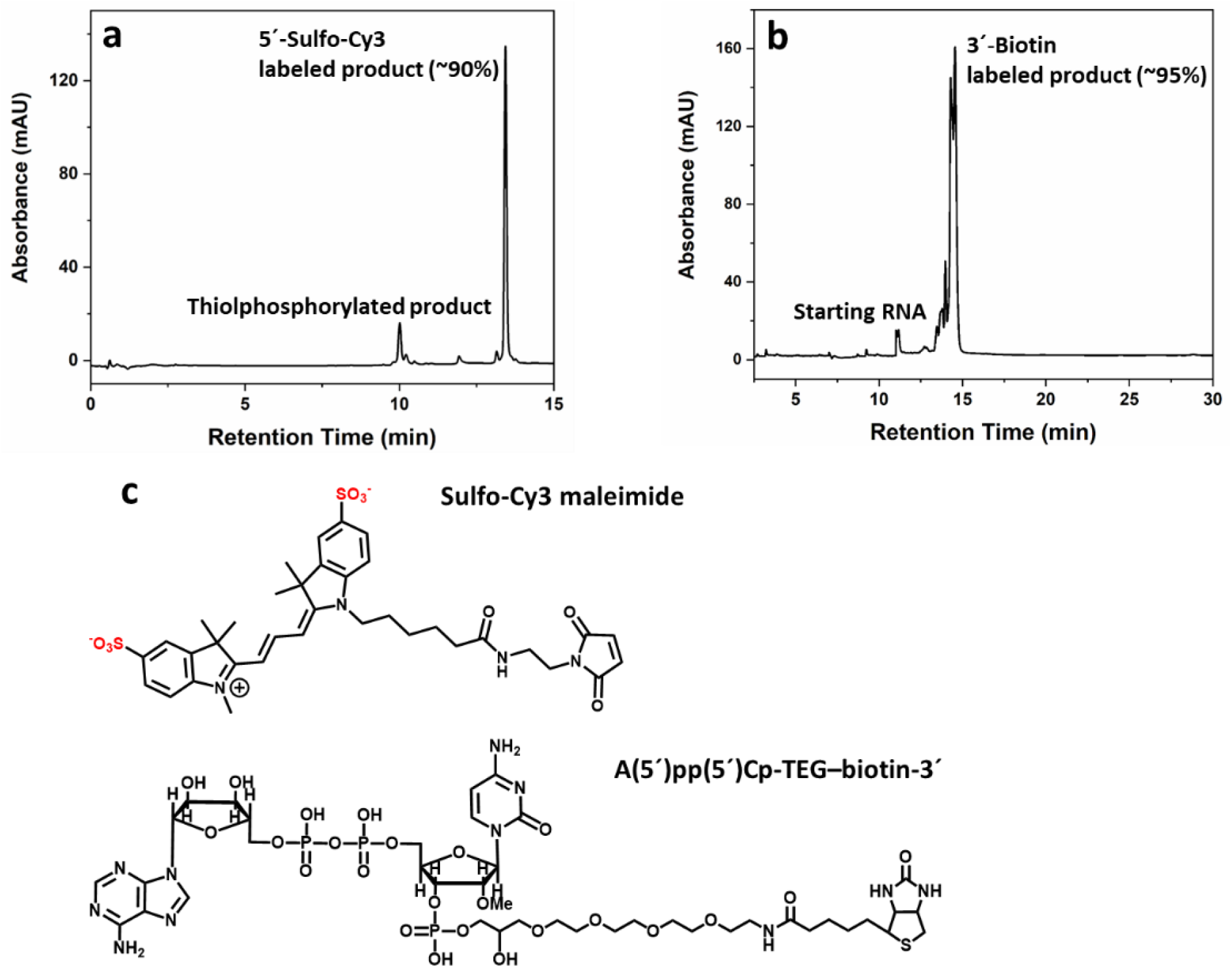
**a)** HPLC profile for the high yield of labeling of RNA #11 with sulfo-Cy3 at the 5’-end. **b)** HPLC profile for the high yield of labeling of RNA #11 with biotin at the 3’-end using A(5’)pp(5’)Cp-TEG-biotin-3’. **c)** Structure of sulfo-Cy3 maleimide and A(5’)pp(5’)Cp-TEG-biotin-3’, applied to achieve a higher labeling efficiency at the 5’- and 3’-ends, respectively.

### LC/MS sequencing of pseudouridine (ψ)

We next decided to apply our new end labeling-LC/MS sequencing strategy to a synthetic sample containing modified nucleobases. Pseudouridine (ψ) is the most abundant and widespread of all modified nucleotides found in RNA. It is present in all species and in many different types of RNAs, including both coding RNAs (mRNAs) and non-coding RNAs (58). However, it is impossible to distinguish ψ from U directly by MS because they have identical masses. An established chemical labeling approach was previously developed to distinguish ψ from U, relying on a nucleophilic addition with *N*-cyclohexyl-*N*’-(2-morpholinoethyl)-carbodiimide metho-p-toluenesulfonate (CMC) to form a CMC-ψ adduct (55). The CMC-ψ adduct stalls reverse transcription and terminates the cDNA one nucleotide towards the 3’ end downstream to it and is currently used to detect ψ sites in various RNAs at single-base resolution (55). Here, we adopt the same chemistry to form the same CMC-ψ adduct in our system (Figure 4a). The adduct will not only have a unique mass 252.2076 Dalton larger than U’s mass, but it is also more hydrophobic than the U, also resulting in an t_R_ shift. The CMC-ψ adduct will thus significantly shift both the masses and t_R_ s of all the ladder fragments containing the CMC-ψ adduct in the mass-t_R_ plot, which will help downstream analysis in identifying and locating the ψ in any of the RNA strands.

**Figure 4.**
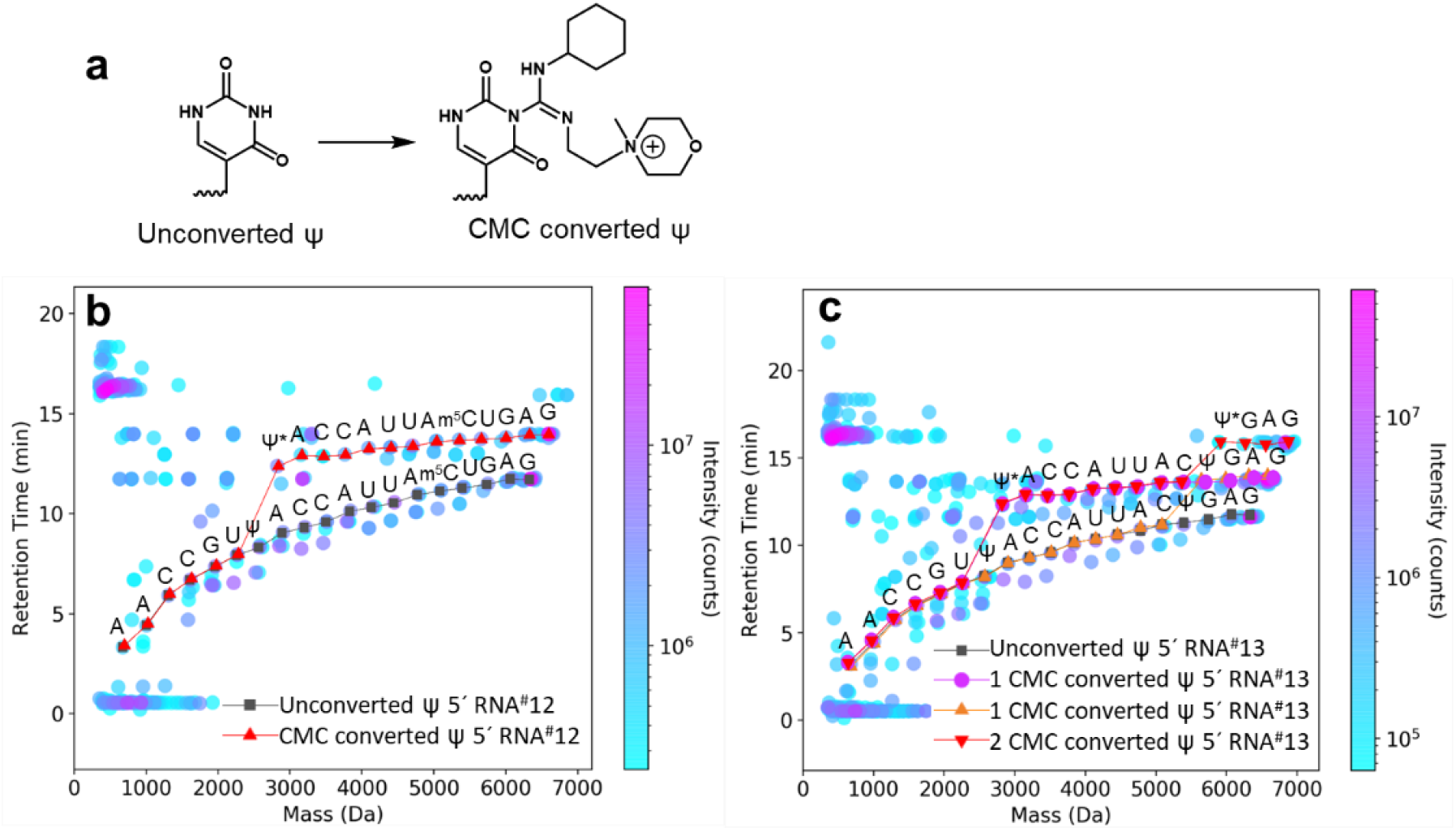
**a)** chemical conversion of pseudouridine (ψ) by reaction with *N*-cyclohexyl-*N*’-(2-morpholinoethyl)-carbodiimide metho-*p*-toluenesulfonate (CMC) to form CMC-ψ, shifting CMC-ψ-containing mass-t_R_ ladders in both mass and t_R_ compared to mass-t_R_ ladders containing unconverted ψ. **b)** sequencing of RNA #12, which contains 1 ψ. The CMC-converted ψ (depicted as ψ*) results in a shift in both t_R_ and mass, allowing facile identification and location of ψ at this position due to a single drastic jump in the mass-t_R_ ladder. c) sequencing of RNA #13, which contains 2 ψ. Each of the CMC-converted ψ (depicted as ψ*) results in a drastic jump in the mass-t_R_ ladder, corresponding to the locations of the ψ in the RNA sequence. For ease of visualization, only the sequences of the 5’-mass-t_R_ ladders are presented. Sequences were read manually from mass-t_R_ ladders identified from the algorithm-processed data.

Figures S3a and S3b show the HPLC profiles of the crude products of converting ψ-containing to their respective CMC-adducts in two RNAs using the reported conditions (55). These two RNAs contain 1 ψ and 2 ψ moieties, respectively (RNA #12 and #13). The conversion percentage of ψ-containing RNA calculated by integrating peaks from UV chromatogram was ~42% and ~64%, respectively. For the RNA strand containing 2 ψ, CMC conversion could be complete (both ψ were converted to CMC-ψ adducts) or partial (only one of the 2 ψ was converted), or none (none of the 2 ψ were converted to CMC-ψ adducts). Therefore, in Figure S3b, the peak around 16 min refers to the RNA strand with complete conversion (~24%), and the two adjacent peaks around 14 min reflect the partial conversion of either ψ (total ~40%).

Automated sequencing was applied to RNA #12 and #13 after acid degradation by formic acid. In the 2-D mass-t_R_ plot representing sequencing of a single ψ-containing RNA (RNA #12, which also contained a single m^5^C (5-methylcytidine)) (Figure 4b), a new curve (red) branched up off of the original sigmoidal curve (grey) at the ψ, corresponding to the part of the sequence with all CMC-ψ adduct-containing ladder fragments, which shift up and to the right in the 2-D mass t_R_ plot because the fragments with CMC-ψ adduct have 252.2076 Dalton larger masses and larger t_R_ s than their corresponding unreacted ones. Figure 4c depicts the 2-D mass-t_R_ plot representing sequencing of a 2 ψ-containing RNA (RNA #13). Similarly, one new curve (red) branched off at the second ψ, corresponding to the part of the sequence with conversion of both ψ to their CMC-ψ adducts. Two additional curves (purple and orange) branched up off of the original unconverted 5’-ladder (grey curve) separately in each of two positions of the ψ, indicating that the only one of two ψ was converted. As such, we can not only identify, locate, and quantify the base modification ψ in the ψ - containing RNA while reading out its complete sequence, but with further calculations incorporating the LC/MS profiles of the undegraded RNA sample (before the chemical degradation for ladder generation), we can also directly quantify the percentage of the CMC-containing RNA *vs* non-CMC-containing RNA in a given sample.

### Quantifying stoichiometry/percentage of modified RNA in a partially modified RNA sample

Understanding the dynamics of cellular RNA modifications (59,60) requires a method to quantify the stoichiometry/percentage of RNA with site-specific modifications *vs*. its canonical counterpart RNA, as base modifications may not occur on 100% of all identical RNA sequences in a cell or sample. Applying the above quantification strategy to other sequences, we expect that this method can allow us to accurately determine the percentage of RNA with any mass-altered modification *vs*. its corresponding non-modified counterpart. As shown in Figure5, not only can the complete sequence including the m^5^C be read out accurately from the mixture containing both modified and non-modified RNA (Figure 5a), but the relative percentage of m^5^C modified RNA (20%) *vs*. its non-modified counterpart (80%) can also be quantified based upon information from the extracted ion chromatograph (Figure 5b) (56). The relative quantities of different product species were quantified by integrating the extracted ion current (EIC) peaks of 3’-biotin labeled methylated RNA and nonmodified RNA before their formic acid degradation (*Supplementary Materials, Table S41*). In addition to sequencing, RNA mixtures with other different ratios have also been quantified similarly (Figure 5b). These relative percentages match well with the ratios of the absolute amounts of RNA initially used for RNA labeling with a difference less than 5%, indicating that EIC-based integration is an accurate method for relative quantification of modified RNA when not every RNA with the same sequence was modified.

**Figure 5.**
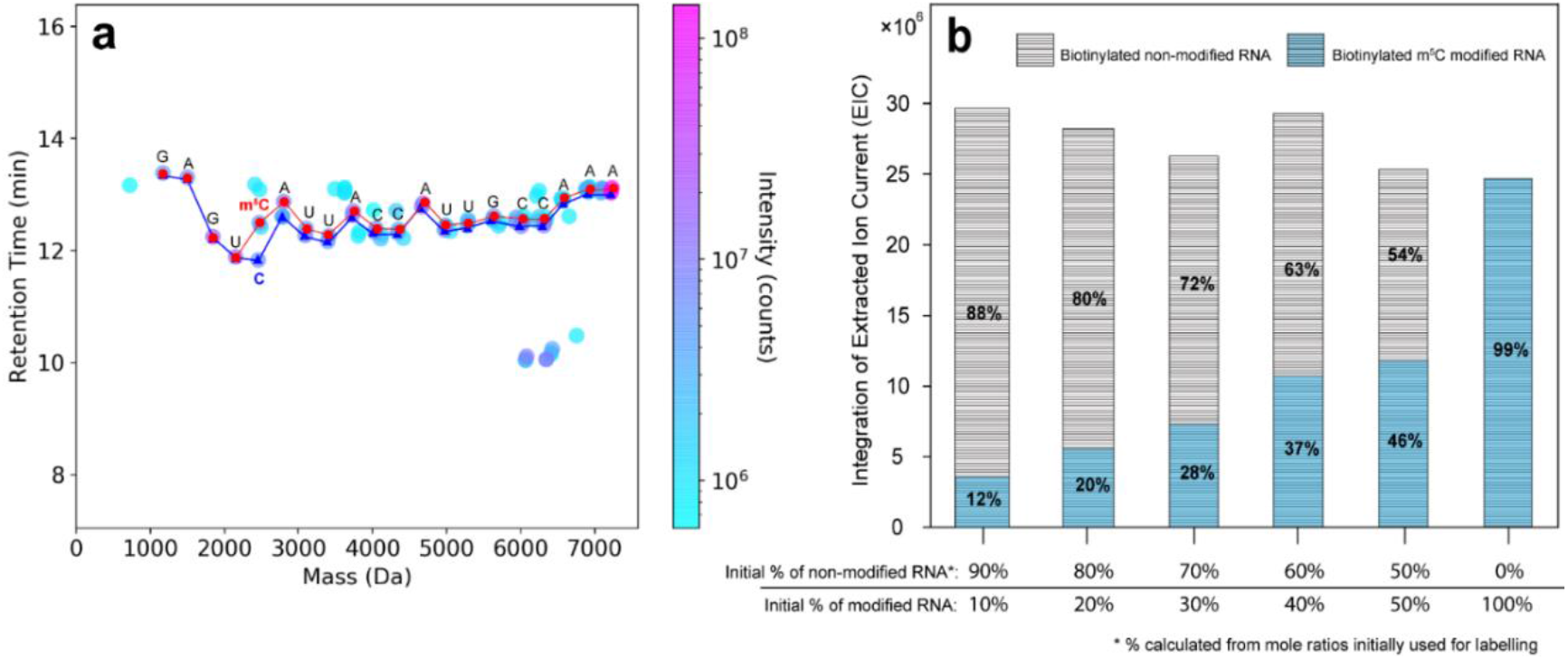
LC/MS sequencing and quantification. **a)** Sequencing of a mixture containing 20% m^5^C modified RNA (RNA #14) and 80% of non-modified RNA (RNA #3). Both curves share the identical sequence until the first C is reached; the t_R_ of the m^5^C-terminated ladder fragment was shifted up (due to the hydrophobicity increase from the methyl group) and the mass slightly increased (due to the 14 Da mass increase from the additional methyl group) compared to its non-modified counterpart. Both sequences were read manually from mass-t_R_ ladders identified from the algorithm-processed data. **b)** Quantifying the stoichiometry/percentage of RNA with modifications *vs*. its canonical counterpart RNA. The relative percentages are quantified by integrating the extracted ion current (EIC) of different labeled product species, and they match well with ratios of the absolute amounts initially used for labeling these RNA samples, *i.e*., percentages of m^5^C modified RNA in the mixed samples were 10%, 20%, 30%, 40%, 50% and 100%, respectively, which was calculated from their mole ratios initially used for labeling.

Extending this idea to ψ, this method can allow us to estimate the percentage of ψ-containing RNA *vs*. non-ψ-containing RNA if we can factor in the yield of CMC chemistry with ψ. Further optimization of CMC-labeling chemistry yield to quantitative would allow the accurate determination of the percentage of ψ-containing RNA *vs*. its unmodified U counterparts.

### Sequencing an RNA mixture with multiple modifications

Finally, with the end-labeling and ψ base-modification strategies in hand, we next sought to increase the throughput of the method in order to sequence a multiplex RNA sample (simultaneous sequencing of a mixed sample containing multiple distinct RNA sequences) containing RNA strands with multiple modifications. We subjected a sample mixture containing 12 RNAs with distinct sequences, containing 11 unmodified RNAs and one multiply-modified RNA (containing 1 ψ and 1 m^5^C), to our protocol. We first chemically labeled the 3’-ends of all RNA samples with biotin, while sulfo-Cy3 was added to the 5’-ends (except for the RNA strand containing the base modifications). After measurement by LC/MS, the data was analyzed using Agilent MassHunter Qualitative Analysis software with optimized MFE (molecular feature extraction) settings to extract data for sequence generation. With the improvements in labeling efficiency described above, we were able to detect all ladder fragments needed to accurately read out the complete sequences of all RNAs in the mixture. In the analysis of the multiplexed samples, the typical processing and basecalling algorithm (as was used in all previous figures) was not used mainly due to the largely increased data complexity resulting from the mixture. These sequences were base-called manually, and all sequences could be accurately sequenced (Figures 6a and 6b).

**Figure 6.**
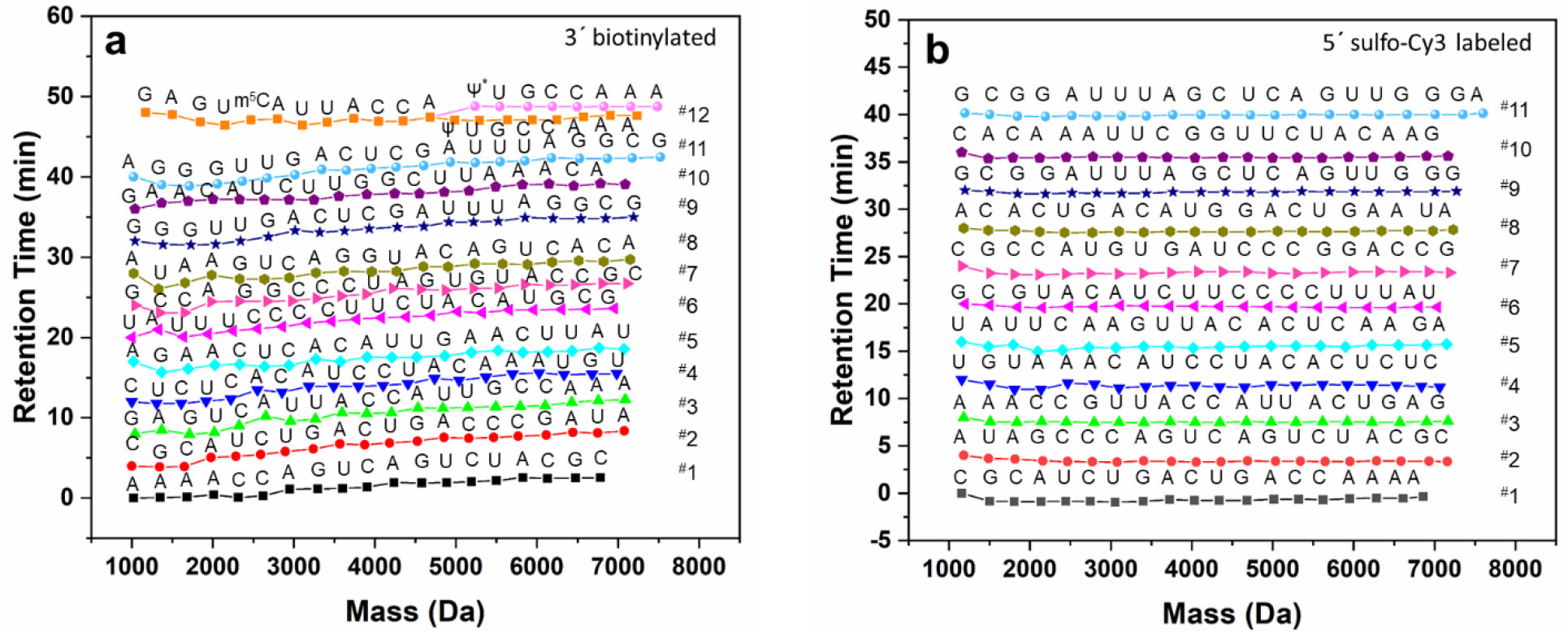
Simultaneous sequencing of a mixed sample containing 12 RNAs with either **a)** a single biotin label at the 3’-end or **b)** a single sulfo-Cy3 label at the 5’-end of each RNA (RNA #12 was only in the 3’-biotin-labeled sample mixture, and thus a) contains one additional sequence compared to b). t_R_ was normalized for ease of visualization (*MATERIAL AND METHODS*).

The results showed that we not only could sequence the four canonical nucleosides (A, C, G and U), but also identify, locate, and quantify multiple modified bases, such as ψ and m^5^C but not limited to just these modified nucleotides, at single-base resolution by mapping their masses in both single and mixed RNA samples. Similarly, for sequencing ψ-containing RNA, we treated RNA with CMC as described before, thus a new curve branched off of its corresponding non-CMC-containing ladder curve at the ψ (pink trace). Although in these studies we manually read out the sequences, as opposed to using an automated processing and basecalling algorithm, these studies show that there are no experimental or physical limitations in the sample preparation and mass spectrometry aspects of our system; the mass ladders of each component of the mixture can be properly generated and can be accurately sequenced and basecalled by the mass-t_R_ plot generated by the MFE file extracted from the LC/MS as a proof-of-principle. As the current algorithm is not yet optimized for automated basecalling of multiple sequences and multiple modified nucleotides simultaneously, further development of the basecalling algorithm can lead to increased throughput through automated basecalling and sequencing of multiplexed samples. These results show that our direct RNA method can sequence more complex RNA samples with multiple RNAs containing modified bases, not just limited to purified single RNA containing one noncanonical base as previously published.(46) It is a significant step forward for MS sequencing of various complex biological RNA samples. Once the automated sequencing algorithm is optimized, the exact capacity of this method’s throughput, *i.e*., how many RNA strands can be sequenced at one time, should be much larger than the sample containing 12 presented here and remains to be explored. Further studies are in progress, with an immediate next aim to increase the throughput to be able to sequence a mixture containing at least 30 distinct RNAs with the maximum length the instrument can handle, and accommodate multiple RNAs with multiple modified bases so that we can sequence real biological RNA samples with chemical modifications.

### Increase sample usage *via* utilization of internal fragments

The previous mass-t_R_ ladder-based RNA sequencing methods controlled degradation conditions to generate well-defined mass ladders with single cuts for sequencing, as opposed to the unwanted appearance of multiple-cut fragments (46). As such, a 5 min formic acid treatment was performed to digest ~10% of a 20 nt (RNA #3) sample into its corresponding 5’- and 3’-sequencing ladders to minimize formation of internal RNA fragments with more than one cut.(46) Thus, ~90% of the starting material remained intact, and could not yield any sequence information. For real biological samples with low abundance, the fact that ~90% of the sample would be unusable for sequencing results in the previous method’s inability to generate enough sample signal to accurately sequence these low-abundance samples. In order to increase the percentage of usable sample, a longer degradation step is required. However, the process of generating more of the desired ladder fragments in a longer chemical/enzymatic degradation step will lead to the production of large amounts of internal fragments that do not possess a 5’- or 3’-end from the original RNA sequence by virtue of more than one cut-site on a given sequence (this is a stochastically-controlled process). The previous method (46) disregarded internal fragments simply as “noise” as they were not a part of the RNA ladders that were actually used in determining the sequence of bases and modification analysis. Although there is still inherent information in these internal fragments, utilizing information from internal fragments effectively is difficult because these sequences are mixed with the desired ladder compounds, especially for fragments in the lower mass regions with mass less than 2000 Daltons. In this low mass region, monomer, dimer, and trimer nucleotides from any part of a given RNA strand cannot be easily separated in the LC phase of the LC/MS, leading to difficulty in accurate sequence identification and analysis. However, separation of desired ladder fragments from internal fragments by double-end labeling of the original sample makes it possible to actually take advantage of the previously unused internal fragments. We propose to gather and apply information from the internal fragments with more than one cut towards sequence generation where there are gaps (ironically generated from the same long acidic degradation step that generated the internal fragments) in the reported sequence greater than or equal to one missing base as observed in the sequence ladder of the 2-D mass-t_R_ plot of an RNA sample (RNA #3) which has been subjected to a 60 min degradation step. As shown in Figure S4, by combining three pieces of information: (a) the 5’-ladder, (b) the 3’-ladder, and (c) internal fragments without both ends, RNA sequencing accuracy can be significantly increased as gaps (unassignable bases) in the mass-t_R_ ladder caused by long degradation times can potentially be completely removed.

## CONCLUSIONS

Development of 2-D-mass-t_R_ direct RNA sequencing methodology brings the power of MS-based laddering technology to RNA, addressing a long-standing unmet need in the broad field of RNA modification studies. Not only does it provide a direct method for RNA sequencing without the need of a cDNA intermediate or PCR, it also provides a general method for sequencing multiple base modifications on multiple RNA strands in one single experiment. The method we have developed has been proven successful to sequence short single strands of synthetic RNA (~20 nucleotides) (Figures 1 and 2), and to quantify stoichiometry/percentage of RNA containing modifications *vs*. its canonical counterpart RNA. With end-labeling, we no longer require paired end sequencing for the complete sequence coverage as before; we can read out the complete sequence of a given RNA strand from either the 3’- or the 5’-end, thus increasing the throughput and ease of data analysis. By using end-labeling, we have also been able to extend the method to directly sequence multiplexed RNA mixtures (Figure 6), which is a crucial step forward in MS-based sequencing of cellular RNA samples, typically consisting of mixed RNAs of unknown sequence. Additionally, we demonstrated the power of the method in sequencing multiple modified bases in this work, including pseudouridine and m^5^C (although not limited to just these two modifications), allowing us to identify, locate, and quantify each of these RNA modifications at single base resolution in mixed samples containing 12 RNA strands.

The full potential of the LC/MS sequencing throughput remains to be explored, and it may be instrument-dependent, *i.e*., mass spectrometers with higher resolving powers may lead to increased throughput and/or lower sample requirements. With further improvements in instrument sensitivity, resolution, and commercially available software and further development of automated sequencing algorithms, this MS-based RNA sequencing method has the potential to become a highly robust, easy-to-use, and broadly applicable *de novo* sequencing approach. Such a platform can complement existing next-generation RNA sequencing protocols for in-depth functional studies of chemical modifications carried by endogenous RNAs.

Accordingly, our methods can facilitate the efficient detection of modified nucleotides, resulting in accurate sequencing of modified RNA molecules, including, for example, tRNAs, siRNAs, and synthetic therapeutic oligoribonucleotides having pharmacological properties, as well as mixtures of RNA molecules. Studies are in progress to expand this approach to sequence cellular RNAs with known chemical modifications, such as endogenous tRNA, to benchmark the method’s efficacy in read length and identification of extensive modifications. With continued improvements in read length, we will also expand this direct sequencing strategy to sequence longer RNAs, such as mRNA, and pinpoint the chemical identity and position of nucleotide modifications on these biological RNAs. Once fully optimized, we expect this direct MS-based RNA sequencing method to facilitate the discovery of more unknown modifications along with their location and abundance information, which no other established sequencing methods are currently capable of.

## ACKNOWLEDGEMENT

The authors acknowledge the R21 grant from NIH (1R21HG009576) to S. Z. and W. L. and New York Institute of Technology (NYIT) Institutional Support for Research and Creativity grants to S. Z., which supported this work. T. Z. J. is a research fellow from the Earth-Life Science Institute (ELSI) at Tokyo Institute of Technology. ELSI is a member of the World Premier International Research Center Initiative and is supported in part by the Japanese Ministry of Education, Culture, Sports, Science and Technology. The authors would also like to thank PhD student Xuanting Wang (Columbia University) for assisting in figure-making, and Prof. Jingyue Ju (Columbia University), Drs. James Russo, Shiv Kumar, Xiaoxu Li, Steffen Jockusch, and other members of the Ju lab (Columbia University), Dr. Yongdong Wang (Cerno Bioscience), Meina Aziz (NYIT), and Wenhao Ni (Fordham University) for helpful discussions and suggestions for our manuscript.

## FUNDING

This work was supported by the R21 grant from National Institutes of Health (1R21HG009576) and New York Institute of Technology (NYIT) Institutional Support for Research and Creativity grants.

## CONFLICT OF INTEREST

The authors have filed a provisional patent related to the technology discussed in this manuscript.

