## Supplementary Materials for "A general LC/MS-based RNA sequencing method for direct analysis of multiple-base modifications in RNA mixtures"

As a proof-of-principle of adding a 5´-tag to spatially separate ladders on a retention time ($t_R$) *vs.* mass plot, a simulated mass spectrum peak set for both 5´- and 3´-ladders of a synthetic, unmodified A10 (10-mer of polyadenine) sequence was first generated *in silico*. Each row represents a given mass ladder peak, and each peak was assigned a unitless retention time ($t_R$) and an arbitrarily constant unitless peak volume of 1000. The $t_R$ assigned for each ladder increased systematically with increasing mass, starting with 0 and increasing in 0.1 unit increments. The peak list for the simulated A10 mass spectrum was as follows:

A10 – unmodified MS peak list

| **Mass** | **$t_R$** | **Vol** |
|---|---|---|
| **347.063065** | 0 | 1000 |
| **676.115565** | 0.1 | 1000 |
| **1005.168065** | 0.2 | 1000 |
| **1334.220565** | 0.3 | 1000 |
| **1663.273065** | 0.4 | 1000 |
| **1992.325565** | 0.5 | 1000 |
| **2321.378065** | 0.6 | 1000 |
| **2650.430565** | 0.7 | 1000 |
| **2979.483065** | 0.8 | 1000 |
| **3228.569232** | 0.9 | 1000 |
| **Mass** | **$t_R$** | **Vol** |
| **267.096732** | 0 | 1000 |
| **596.149232** | 0.1 | 1000 |
| **925.201732** | 0.2 | 1000 |
| **1254.254232** | 0.3 | 1000 |
| **1583.306732** | 0.4 | 1000 |
| **1912.359232** | 0.5 | 1000 |
| **2241.411732** | 0.6 | 1000 |
| **2570.464232** | 0.7 | 1000 |
| **2899.516732** | 0.8 | 1000 |
| **3228.569232** | 0.9 | 1000 |

The mass ladder starting from 347.063065 represents the 5´-mass ladder, while the mass ladder starting from the 267.096732 represents the 3´-mass ladder.

Next a simulated mass spectrum peak set for both 5´- and 3´-ladders of a synthetic, 5´-cyanine 3 (Cy3)-labeled A10 (10-mer of polyadenine) sequence was generated *in silico*. This was done by taking the data set above, and adding the additional mass afforded by a 5´-Cy3 label (614.3061) to each member of the 5´-ladder in the data set. The peak volumes did not change. The associated $t_R$ for this new Cy3-labeled 5´-ladder was generated by now starting from an $t_R$ of 10, and decreased by an increment of 0.2 with increasing mass. This was done to simulate the potential change to an $t_R$ *vs*. mass spectrum of any end-labeled ladder (in this case, 5´-Cy3-labeled) in both absolute $t_R$ values, $t_R$ trends (monotonically increasing curve to a monotonically decreasing curve, for example), and absolute mass values. Of course, real changes in all of these values in a real system could not be absolutely predicted *in silico*, and thus this should be only taken as a proof-of-principle example. The peak list for the simulated 5´-Cy3-labeled A10 mass spectrum was as follows:

A10 – 5´-Cy3-labeled MS peak list

| **Mass** | **$t_R$** | **Vol** |
|---|---|---|
| **961.369165** | 3 | 1000 |
| **1290.421665** | 2.8 | 1000 |
| **1619.474165** | 2.6 | 1000 |
| **1948.526665** | 2.4 | 1000 |
| **2277.579165** | 2.2 | 1000 |
| **2606.631665** | 2 | 1000 |
| **2935.684165** | 1.8 | 1000 |
| **3264.736665** | 1.6 | 1000 |
| **3593.789165** | 1.4 | 1000 |
| **3842.875332** | 1.2 | 1000 |
| **Mass** | **$t_R$** | **Vol** |
| **267.096732** | 0 | 1000 |
| **596.149232** | 0.1 | 1000 |
| **925.201732** | 0.2 | 1000 |
| **1254.254232** | 0.3 | 1000 |
| **1583.306732** | 0.4 | 1000 |
| **1912.359232** | 0.5 | 1000 |
| **2241.411732** | 0.6 | 1000 |
| **2570.464232** | 0.7 | 1000 |
| **2899.516732** | 0.8 | 1000 |
| **3228.569232** | 0.9 | 1000 |

The mass ladder starting from 961.369165 represents the 5´-Cy3-labeled mass ladder, while the mass ladder starting from the 267.096732 represents the 3´-mass ladder.

Comparing these two $t_R$ *vs.* mass plots, we see that the two mass ladder curves are almost superimposed when there is no end-labeling (**Fig. S1a**), resulting in potential mis-sequencing in downstream basecalling and sequence identification, while the 5´-Cy3-labeled sample has two distinct and separate mass ladder curves (**Fig. S1b**), which allow for greater ease of visualization of all the ladder components needed for sequencing and higher accuracy in downstream basecalling and sequence identification.

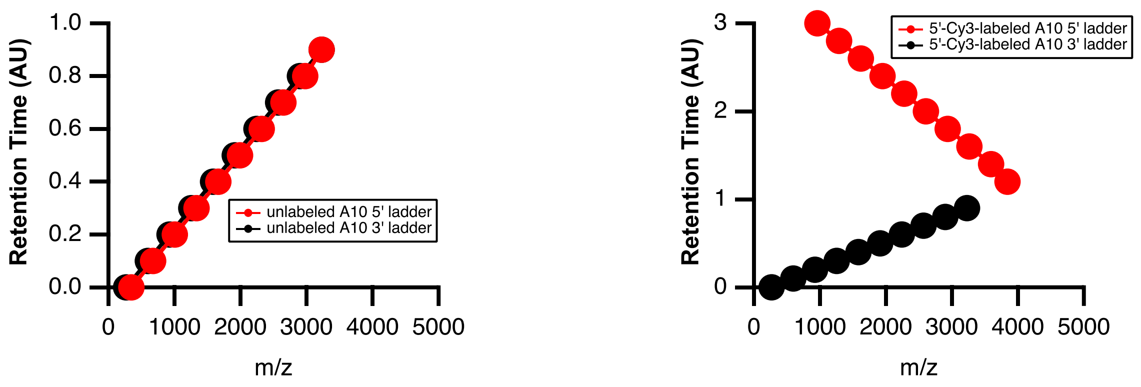


**Fig. S1. a)** Unlabeled 3´- and 5´-mass ladders of a synthetic, unmodified A10 (10-mer of polyadenine) sequence generated *in silico*. **b)** 5´- and 3´-mass ladders of a synthetic, 5´-Cy3-labeled A10 (10-mer of polyadenine) sequence generated *in silico.*

In addition to automating the sequence generation, we also manually searched for the mass ladders by the Molecular Feature Extraction (MFE) workflow in MassHunter Qualitative Analysis (Agilent Technologies), for confirming the accuracy of automating sequencing. In **Tables S1–S41**, we provided the theoretical mass of each fragment (obtained by ChemDraw), base mass, base name, observed mass, $t_R$, volume (peak intensity), quality score, and ppm difference (calculated by the equation as follows). The MFE settings were optimized to extract as many identified compounds as possible but with reasonable quality score. The MFE settings we applied were as follows: "centroid data format, small molecules (chromatographic), peak with height ≥ 500, up to a maximum of 1000, quality score ≥ 50". However, data reduction was performed to simplify algorithm sequencing if needed. For instance, retention time could be selected from 6 to 10 min for biotin labeled samples for a 20 nt RNA (*e.g.*, RNA #2). Also, the numbers of input compounds used for algorithm analysis are generally an order-of-magnitude higher than the number of ladder fragments needed for generating complete sequences, unless indicated otherwise; these input compounds are sorted out of all MFE extracted compounds typically with higher volumes and/or better quality scores.

The formula used to calculate the PPM in the manuscript:

$$\mathbf{ppm} = \mathbf{10^{-6}} \times (\mathbf{Mass_{theoretical}} - \mathbf{Mass_{observed}})/\mathbf{Mass_{theoretical}}$$

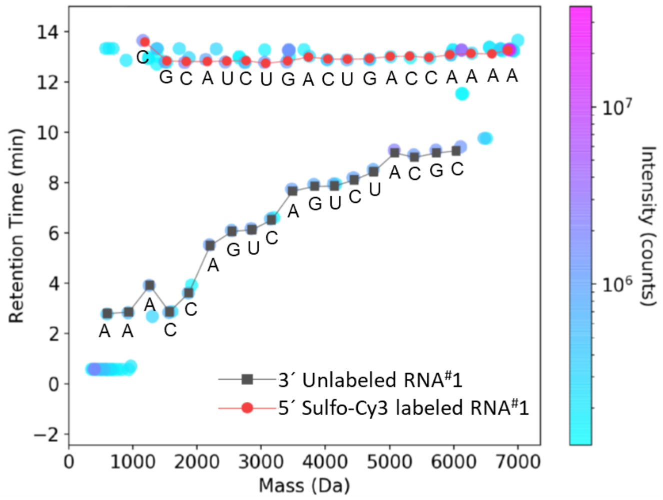


**Fig. S2.** Mass-$t_R$ plot of a sample containing complete sets of ladder fragments from 5´-sulfo-Cy3-labeled RNA #1 and its 3´-unlabeled ladder fragments. Sequences were manually read out from automatically-generated mass-$t_R$ plots containing mass-$t_R$ ladders identified from algorithm-processed data.

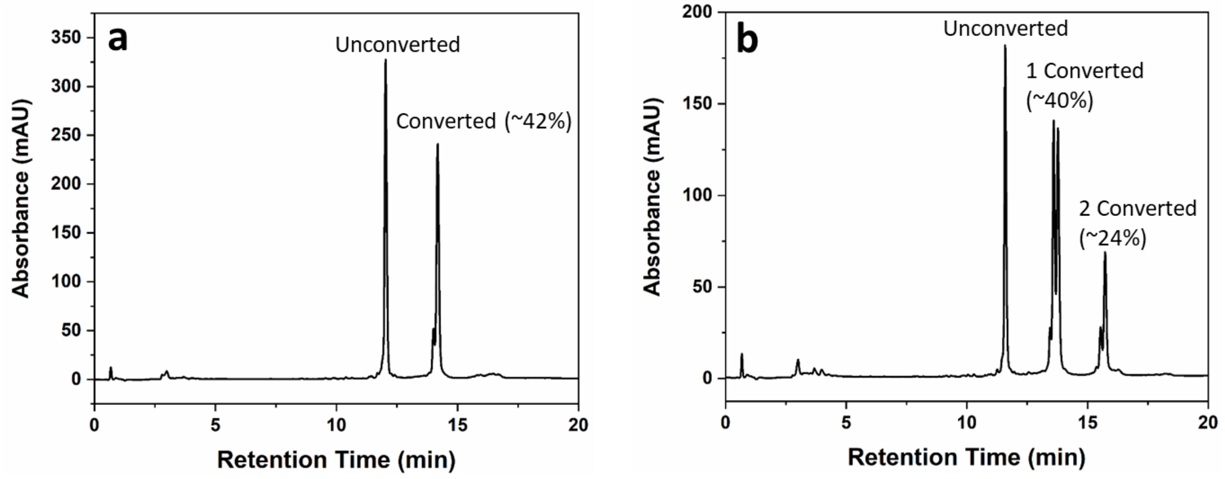


**Fig. S3.** HPLC profile of the crude products after conversion of pseudouridine (ψ) to its *N*-cyclohexyl-*N′*-(2-morpholinoethyl)-carbodiimide metho-*p*-toluenesulfonate (CMC) adduct in **a**) a 20 nt RNA (RNA #12) containing 1 ψ base and **b**) a 20 nt RNA (RNA #13) containing 2 ψ bases.

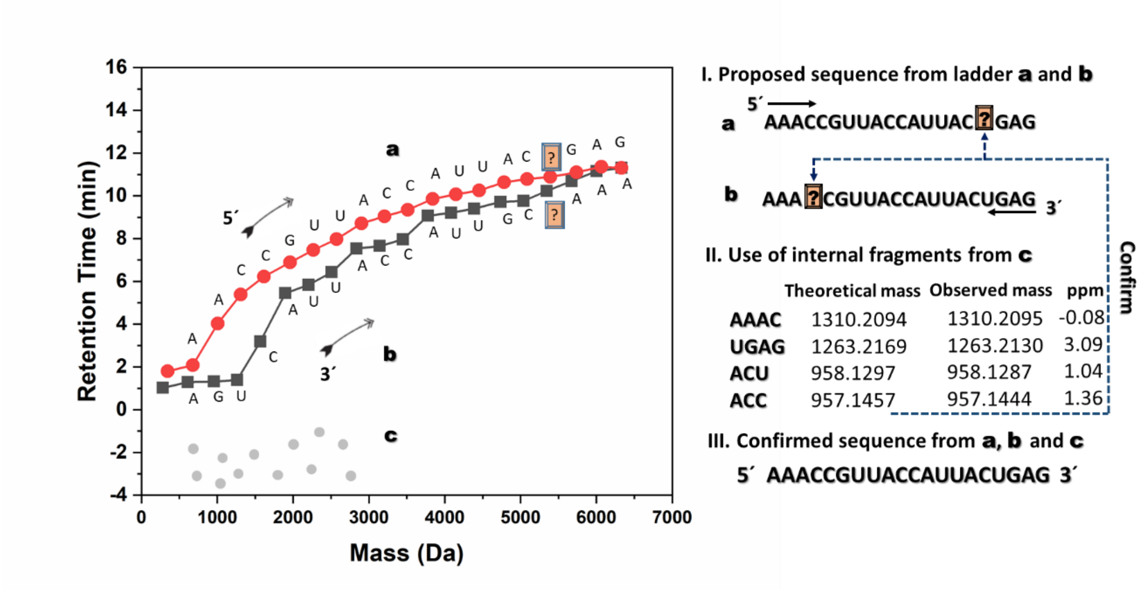


**Fig. S4.** Utilization of internal fragments without either original 5´- or 3´-end to fill gaps in the 5´-ladders ladder before reporting the final sequence of a 20 nt RNA (RNA #3) that was degraded for 60 min, thus increasing the method's accuracy by combining three pieces of information including **(a)** the 5´-ladder, **(b)** the 3´-ladder, and **(c)** internal fragments whose observed masses match with a list of theoretical masses from the proposed sequence.

**Table S1.** LC/MS analysis of 3´-biotin-labeled RNA #1 after isolation by streptavidin beads followed by subsequent chemical degradation (3´-labeled mass ladder components, RNA #1, refers to the dataset for Fig. 1b).

| Theoretical | | | | Extracted data file after LC/MS analysis | | | | Error |
|---|---|---|---|---|---|---|---|---|
| **Fragments** | **Theoretical mass** | **Base mass** | **Base** | **MFE mass** | **$t_R$** | **Volume** | **Quality Score** | **ppm** |
| 19 | 6781.0733 | 305.0413 | C | 6781.0413 | 9.752 | 16819442 | 100 | 4.72 |
| 18 | 6476.0320 | 345.0474 | G | 6475.9924 | 9.717 | 247965 | 84 | 6.11 |
| 17 | 6130.9846 | 305.0413 | C | 6130.9398 | 9.662 | 178841 | 80 | 7.31 |
| 16 | 5825.9433 | 329.0525 | A | 5825.9037 | 9.782 | 510096 | 80 | 6.80 |
| 15 | 5496.8908 | 306.0253 | U | 5496.8566 | 9.383 | 262486 | 99 | 6.22 |
| 14 | 5190.8655 | 305.0413 | C | 5190.8364 | 9.241 | 349988 | 100 | 5.61 |
| 13 | 4885.8242 | 306.0253 | U | 4885.7908 | 9.135 | 356118 | 100 | 6.84 |
| 12 | 4579.7989 | 345.0475 | G | 4579.7738 | 9.109 | 386687 | 100 | 5.48 |
| 11 | 4234.7514 | 329.0525 | A | 4234.7271 | 9.145 | 305380 | 100 | 5.74 |
| 10 | 3905.6989 | 305.0413 | C | 3905.6749 | 8.575 | 145505 | 96 | 6.14 |
| 9 | 3600.6576 | 306.0253 | U | 3600.6373 | 8.420 | 195308 | 100 | 5.64 |
| 8 | 3294.6323 | 345.0474 | G | 3294.6165 | 8.370 | 125991 | 100 | 4.80 |
| 7 | 2949.5849 | 329.0525 | A | 2949.5716 | 8.339 | 106993 | 100 | 4.51 |
| 6 | 2620.5324 | 305.0413 | C | 2620.5193 | 7.492 | 90629 | 100 | 5.00 |
| 5 | 2315.4911 | 305.0413 | C | 2315.4814 | 7.299 | 163692 | 100 | 4.19 |
| 4 | 2010.4498 | 329.0525 | A | 2010.4388 | 7.625 | 279963 | 100 | 5.47 |
| 3 | 1681.3973 | 329.0525 | A | 1681.3891 | 7.354 | 183827 | 100 | 4.88 |
| 2 | 1352.3448 | 329.0526 | A | 1352.3378 | 7.303 | 135065 | 100 | 5.18 |
| 1 | 1023.2922 | 329.0525 | A | 1023.2859 | 7.219 | 106700 | 100 | 6.16 |

**Table S2.** LC/MS analysis of 3´-biotin-labeled RNA #1 after isolation by streptavidin beads followed by subsequent chemical degradation (5´-unlabeled mass ladder components, RNA #1).

| Theoretical | | | | Extracted data file after LC/MS analysis | | | | Error |
|---|---|---|---|---|---|---|---|---|
| Fragments | Theoretical mass | Base mass | Base | MFE mass | $t_R$ | Volume | Quality Score | ppm |
| 19 | 6024.8778 | 249.0862 | A | 6024.8483 | 7.664 | 14325731 | 100 | 4.90 |
| 18 | 5775.7916 | 329.0525 | A | 5775.7522 | 7.701 | 457844 | 87 | 6.82 |
| 17 | 5446.7391 | 329.0525 | A | 5446.6965 | 7.411 | 417145 | 100 | 7.82 |
| 16 | 5117.6866 | 329.0525 | A | 5117.6572 | 7.105 | 490290 | 100 | 5.74 |
| 15 | 4788.6341 | 305.0413 | C | 4788.606 | 6.685 | 728135 | 100 | 5.87 |
| 14 | 4483.5928 | 305.0413 | C | 4483.5657 | 6.428 | 481770 | 100 | 6.04 |
| 13 | 4178.5515 | 329.0525 | A | 4178.5286 | 6.183 | 297514 | 100 | 5.48 |
| 12 | 3849.499 | 345.0475 | G | 3849.4787 | 5.653 | 518403 | 100 | 5.27 |
| 11 | 3504.4515 | 306.0253 | U | 3504.4331 | 5.238 | 614494 | 100 | 5.25 |
| 10 | 3198.4262 | 305.0413 | C | 3198.4106 | 4.785 | 524613 | 99 | 4.88 |
| 9 | 2893.3849 | 329.0525 | A | 2893.3714 | 4.341 | 373933 | 100 | 4.67 |
| 8 | 2564.3324 | 345.0474 | G | 2564.3219 | 3.458 | 509219 | 100 | 4.09 |
| 7 | 2219.285 | 306.0253 | U | 2219.2752 | 2.84 | 579139 | 100 | 4.42 |
| 6 | 1913.2597 | 305.0413 | C | 1913.2521 | 2.081 | 466058 | 100 | 3.97 |
| 5 | 1608.2184 | 306.0253 | U | 1608.2123 | 1.375 | 372038 | 80 | 3.79 |
| 4 | 1302.1931 | 329.0525 | A | 1302.1878 | 0.925 | 240613 | 100 | 4.07 |
| 3 | 973.1406 | 305.0413 | C | 973.1367 | 0.765 | 208989 | 100 | 4.01 |
| 2 | 668.0993 | 345.0474 | G | 668.0955 | 0.652 | 26061 | 100 | 5.69 |
| 1 | 323.0519 | 305.0413 | C | NA* | NA | NA | NA | NA |

* NA: Not Analyzed. The 350 Da threshold was set to minimize background ions from the elution buffers. Otherwise, we would predominantly detect HFIP and DPA ions. Thus, the masses which are smaller than 350 Da were not detected.

**Table S3.** LC/MS analysis of 5´-biotin-labeled RNA #1 (5´-labeled mass ladder components, RNA #1).

| Theoretical | | | | Extracted data file after LC/MS analysis | | | | Error |
|---|---|---|---|---|---|---|---|---|
| **Fragments** | **Theoretical mass** | **Base mass** | **Base** | **MFE mass** | **$t_R$** | **Volume** | **Quality Score** | **ppm** |
| 19 | 6600.0415 | 249.0862 | A | 6600.0153 | 10.113 | 1468018 | 100 | 3.97 |
| 18 | 6350.9553 | 329.0525 | A | 6350.9006 | 10.094 | 139388 | 80 | 8.61 |
| 17 | 6021.9028 | 329.0525 | A | 6021.8665 | 9.957 | 152155 | 80 | 6.03 |
| 16 | 5692.8503 | 329.0525 | A | 5692.8225 | 9.806 | 122377 | 84 | 4.88 |
| 15 | 5363.7978 | 305.0413 | C | 5363.7567 | 9.594 | 255396 | 100 | 7.66 |
| 14 | 5058.7565 | 305.0413 | C | 5058.732 | 9.508 | 169499 | 80 | 4.84 |
| 13 | 4753.7152 | 329.0525 | A | 4753.6944 | 9.449 | 121869 | 96 | 4.38 |
| 12 | 4424.6627 | 345.0475 | G | 4424.6389 | 9.204 | 222046 | 100 | 5.38 |
| 11 | 4079.6152 | 306.0253 | U | 4079.5902 | 9.067 | 296271 | 100 | 6.13 |
| 10 | 3773.5899 | 305.0413 | C | 3773.5679 | 8.937 | 249085 | 100 | 5.83 |
| 9 | 3468.5486 | 329.0525 | A | 3468.5308 | 8.838 | 185624 | 100 | 5.13 |
| 8 | 3139.4961 | 345.0474 | G | 3139.4834 | 8.507 | 319911 | 100 | 4.05 |
| 7 | 2794.4487 | 306.0253 | U | 2794.436 | 8.288 | 380189 | 100 | 4.54 |
| 6 | 2488.4234 | 305.0413 | C | 2488.4134 | 8.073 | 317954 | 100 | 4.02 |
| 5 | 2183.3821 | 306.0253 | U | 2183.3725 | 7.863 | 305479 | 100 | 4.40 |
| 4 | 1877.3568 | 329.0525 | A | 1877.3489 | 7.642 | 222446 | 100 | 4.21 |
| 3 | 1548.3043 | 305.0413 | C | 1548.2982 | 7.088 | 361254 | 100 | 3.94 |
| 2 | 1243.263 | 345.0474 | G | 1243.2575 | 6.798 | 162972 | 100 | 4.42 |
| 1 | 898.2156 | 305.0413 | C | 898.2105 | 6.880 | 88421 | 100 | 5.68 |

**Table S4.** LC/MS analysis of 5´-biotin-labeled RNA #2 (5´-labeled mass ladder components, RNA #2).

| Theoretical | | | | Extracted data file after LC/MS analysis | | | | Error |
|---|---|---|---|---|---|---|---|---|
| **Fragments** | **Theoretical mass** | **Base mass** | **Base** | **MFE mass** | **$t_R$** | **Volume** | **Quality Score** | **ppm** |
| 20 | 6898.0505 | 225.075 | C | 6898.0210 | 10.014 | 3995416 | 100 | 4.28 |
| 19 | 6672.9755 | 345.0474 | G | 6673.4755 | 10.115 | 92706 | 80 | -74.93 |
| 18 | 6327.9281 | 305.0413 | C | 6327.8894 | 10.117 | 108088 | 80 | 6.12 |
| 17 | 6022.8868 | 329.0525 | A | 6022.8313 | 10.104 | 133027 | 100 | 9.21 |
| 16 | 5693.8343 | 306.0253 | U | 5693.7870 | 9.920 | 68281 | 80 | 8.31 |
| 15 | 5387.809 | 305.0413 | C | 5387.7785 | 9.850 | 167081 | 80 | 5.66 |
| 14 | 5082.7677 | 306.0253 | U | 5082.7314 | 9.784 | 170198 | 100 | 7.14 |
| 13 | 4776.7424 | 345.0474 | G | 4776.7210 | 9.695 | 114657 | 99 | 4.48 |
| 12 | 4431.695 | 329.0526 | A | 4431.6685 | 9.629 | 143358 | 92 | 5.98 |
| 11 | 4102.6424 | 305.0412 | C | 4102.6199 | 9.367 | 245033 | 100 | 5.48 |
| 10 | 3797.6012 | 306.0253 | U | 3797.5819 | 9.264 | 184127 | 100 | 5.08 |
| 9 | 3491.5759 | 345.0475 | G | 3491.5567 | 9.131 | 91691 | 100 | 5.50 |
| 8 | 3146.5284 | 329.0525 | A | 3146.5054 | 9.028 | 187937 | 100 | 7.31 |
| 7 | 2817.4759 | 305.0413 | C | 2817.4633 | 8.675 | 288050 | 100 | 4.47 |
| 6 | 2512.4346 | 305.0413 | C | 2512.4233 | 8.509 | 138698 | 100 | 4.50 |
| 5 | 2207.3933 | 305.0413 | C | 2207.3835 | 8.335 | 192998 | 100 | 4.44 |
| 4 | 1902.352 | 345.0474 | G | 1902.3433 | 8.161 | 149466 | 100 | 4.57 |
| 3 | 1557.3046 | 329.0525 | A | 1557.2976 | 8.042 | 133349 | 100 | 4.49 |
| 2 | 1228.2521 | 306.0253 | U | 1228.2455 | 7.618 | 188828 | 100 | 5.37 |
| 1 | 922.2268 | 329.0525 | A | 922.2213 | 7.434 | 86674 | 100 | 5.96 |

**Table S5.** LC/MS analysis of 3´-biotin-labeled RNA#1 (3´-labeled mass ladder components, RNA #1, refers to the dataset for Fig. 6a, among a mixture of 12 RNA strands).

| Theoretical | | | | Extracted data file after LC/MS analysis | | | | Error |
|---|---|---|---|---|---|---|---|---|
| **Fragments** | **Theoretical mass** | **Base mass** | **Base** | **MFE mass** | **$t_R$** | **Volume** | **Quality Score** | **ppm** |
| 19 | 6781.0733 | 305.0413 | C | 6781.0476 | 9.552 | 1439108 | 100 | 3.79 |
| 18 | 6476.0320 | 345.0474 | G | 6475.9807 | 9.525 | 256582 | 90 | 7.92 |
| 17 | 6130.9846 | 305.0413 | C | 6130.9052 | 9.466 | 208256 | 80 | 12.95 |
| 16 | 5825.9433 | 329.0525 | A | 5825.8968 | 9.593 | 309638 | 99 | 7.98 |
| 15 | 5496.8908 | 306.0253 | U | 5496.8429 | 9.198 | 241141 | 95 | 8.71 |
| 14 | 5190.8655 | 305.0413 | C | 5190.8331 | 9.058 | 407162 | 100 | 6.24 |
| 13 | 4885.8242 | 306.0253 | U | 4885.7984 | 8.959 | 408024 | 100 | 5.28 |
| 12 | 4579.7989 | 345.0475 | G | 4579.7712 | 8.937 | 431600 | 100 | 6.05 |
| 11 | 4234.7514 | 329.0525 | A | 4234.7262 | 8.976 | 490860 | 100 | 5.95 |
| 10 | 3905.6989 | 305.0413 | C | 3905.6751 | 8.419 | 257315 | 100 | 6.09 |
| 9 | 3600.6576 | 306.0253 | U | 3600.638 | 8.271 | 336323 | 100 | 5.44 |
| 8 | 3294.6323 | 345.0474 | G | 3294.6175 | 8.228 | 433533 | 100 | 4.49 |
| 7 | 2949.5849 | 329.0525 | A | 2949.5701 | 8.205 | 431168 | 100 | 5.02 |
| 6 | 2620.5324 | 305.0413 | C | 2620.5193 | 7.374 | 163100 | 100 | 5.00 |
| 5 | 2315.4911 | 305.0413 | C | 2315.4814 | 7.192 | 366354 | 100 | 4.19 |
| 4 | 2010.4498 | 329.0525 | A | 2010.4386 | 7.528 | 703696 | 100 | 5.57 |
| 3 | 1681.3973 | 329.0525 | A | 1681.3894 | 7.274 | 439312 | 100 | 4.70 |
| 2 | 1352.3448 | 329.0526 | A | 1352.3375 | 7.236 | 326818 | 100 | 5.40 |
| 1 | 1023.2922 | 1023.2922 | A | 1023.2871 | 7.156 | 229472 | 100 | 4.98 |

**Table S6.** LC/MS analysis of 5´-Cy3-labeled RNA#1 (5´-labeled mass ladder components, RNA #1).

| Theoretical | | | | Extracted data file after LC/MS analysis | | | | Error |
|---|---|---|---|---|---|---|---|---|
| **Fragments** | **Theoretical mass** | **Base mass** | **Base** | **MFE mass** | **$t_R$** | **Volume** | **Quality Score** | **ppm** |
| 19 | 6699.1470 | 249.0862 | A | 6699.1256 | 18.524 | 5427844 | 100 | 3.19 |
| 18 | 6450.0608 | 329.0525 | A | 6449.9835 | 18.332 | 53422 | 63 | 11.98 |
| 17 | 6121.0083 | 329.0525 | A | 6120.8891 | 18.514 | 169274 | 65 | 19.47 |
| 16 | 5791.9558 | 329.0525 | A | 5791.9216 | 18.714 | 144098 | 80 | 5.90 |
| 15 | 5462.9033 | 305.0413 | C | 5462.8752 | 18.912 | 209335 | 80 | 5.14 |
| 14 | 5157.8620 | 305.0413 | C | 5157.8321 | 19.171 | 126348 | 88 | 5.80 |
| 13 | 4852.8207 | 329.0525 | A | 4852.7935 | 19.463 | 73470 | 77 | 5.60 |
| 12 | 4523.7682 | 345.0475 | G | 4523.7443 | 19.727 | 116108 | 80 | 5.28 |
| 11 | 4178.7207 | 306.0253 | U | 4178.7014 | 20.053 | 150111 | 79 | 4.62 |
| 10 | 3872.6954 | 305.0413 | C | 3872.6719 | 20.452 | 67114 | 60 | 6.07 |
| 9 | 3567.6541 | 329.0525 | A | 3567.6422 | 20.91 | 36809 | 56 | 3.34 |
| 8 | 3238.6016 | 345.0474 | G | 3238.5865 | 21.394 | 96534 | 93 | 4.66 |
| 7 | 2893.5542 | 306.0253 | U | 2893.5415 | 22.048 | 102530 | 80 | 4.39 |
| 6 | 2587.5289 | 305.0413 | C | 2587.5194 | 22.816 | 35118 | 61 | 3.67 |
| 5 | 2282.4876 | 306.0253 | U | 2282.4795 | 23.767 | 35793 | 86 | 3.55 |
| 4 | 1976.4623 | 329.0525 | A | 1976.4542 | 24.828 | 202040 | 100 | 4.10 |
| 3 | 1647.4098 | 305.0413 | C | 1647.4021 | 26.428 | 220072 | 100 | 4.67 |
| 2 | 1342.3685 | 345.0474 | G | 1342.3610 | 28.326 | 110504 | 100 | 5.59 |
| 1 | 997.3210 | 305.0413 | C | NA* | NA | NA | NA | NA |

* NA: Not Analyzed. The 350 Da threshold was set to minimize background ions from the elution buffers. Otherwise, we would predominantly detect HFIP and DPA ions. Thus, the masses which are smaller than 350 Da were not detected.

**Table S7.** LC/MS analysis of a 1 ψ-containing RNA #12 (ψ unconverted mass ladder components from 5´ to 3´, RNA #12).

| Theoretical | | | | Extracted data file after LC/MS analysis | | | | Error |
|---|---|---|---|---|---|---|---|---|
| **Fragments** | **Theoretical mass** | **Base mass** | **Base** | **MFE mass** | **$t_R$** | **Volume** | **Quality Score** | **ppm** |
| 20 | 6345.9028 | 265.0811 | G | 6345.9217 | 11.736 | 41088112 | 100 | -2.98 |
| 19 | 6080.8217 | 329.0525 | A | 6080.8255 | 11.769 | 2582596 | 100 | -0.62 |
| 18 | 5751.7692 | 345.0474 | G | 5751.7749 | 11.496 | 2169051 | 100 | -0.99 |
| 17 | 5406.7218 | 306.0253 | U | 5406.7209 | 11.315 | 2126771 | 100 | 0.17 |
| 16 | 5100.6965 | 319.057 | $m^5C$ | 5100.6941 | 11.167 | 1149416 | 100 | 0.47 |
| 15 | 4781.6395 | 329.0525 | A | 4781.6402 | 10.970 | 2692877 | 100 | -0.15 |
| 14 | 4452.5870 | 306.0253 | U | 4452.5866 | 10.566 | 5448251 | 100 | 0.09 |
| 13 | 4146.5617 | 306.0253 | U | 4146.5603 | 10.343 | 4115258 | 100 | 0.34 |
| 12 | 3840.5364 | 329.0526 | A | 3840.5352 | 10.141 | 2038738 | 100 | 0.31 |
| 11 | 3511.4838 | 305.0413 | C | 3511.4836 | 9.610 | 1167942 | 100 | 0.06 |
| 10 | 3206.4425 | 305.0412 | C | 3206.4401 | 9.331 | 3422282 | 100 | 0.75 |
| 9 | 2901.4013 | 329.0526 | A | 2901.3988 | 9.067 | 2391922 | 100 | 0.86 |
| 8 | 2572.3487 | 306.0253 | Unconverted ψ | 2572.3468 | 8.328 | 4952174 | 100 | 0.74 |
| 7 | 2266.3234 | 306.0253 | U | 2266.3215 | 7.944 | 4534905 | 100 | 0.84 |
| 6 | 1960.2981 | 345.0474 | G | 1960.2956 | 7.360 | 3437270 | 100 | 1.28 |
| 5 | 1615.2507 | 305.0413 | C | 1615.2481 | 6.693 | 4151449 | 100 | 1.61 |
| 4 | 1310.2094 | 305.0413 | C | 1310.2062 | 5.915 | 1289241 | 87 | 2.44 |
| 3 | 1005.1681 | 329.0525 | A | 1005.1655 | 4.416 | 913589 | 100 | 2.59 |
| 2 | 676.1156 | 329.0525 | A | 676.1140 | 3.321 | 748977 | 100 | 2.37 |
| 1 | 347.0631 | 329.0525 | A | NA* | NA | NA | NA | NA |

* NA: Not Analyzed. The 350 Da threshold was set to minimize background ions from the elution buffers. Otherwise, we would predominantly detect HFIP and DPA ions. Thus, the masses which are smaller than 350 Da were not detected.

**Table S8.** LC/MS analysis of a 1 ψ-containing RNA #12 (ψ unconverted mass ladder components from 3´ to 5´, RNA #12).

| Theoretical | | | | Extracted data file after LC/MS analysis | | | | Error |
|---|---|---|---|---|---|---|---|---|
| **Fragments** | **Theoretical mass** | **Base mass** | **Base** | **MFE mass** | **$t_R$** | **Volume** | **Quality Score** | **ppm** |
| 20 | 6345.9028 | 329.0525 | A | 6345.9069 | 11.361 | 91693 | 61 | -0.65 |
| 19 | 6016.8503 | 329.0525 | A | 6016.856 | 11.603 | 2102227 | 96 | -0.95 |
| 18 | 5687.7978 | 329.0525 | A | 5687.8032 | 11.149 | 1349414 | 100 | -0.95 |
| 17 | 5358.7453 | 305.0413 | C | 5358.7538 | 10.493 | 1095672 | 100 | -1.59 |
| 16 | 5053.7040 | 305.0413 | C | 5053.7053 | 10.247 | 1906586 | 100 | -0.26 |
| 15 | 4748.6627 | 345.0475 | G | 4748.6638 | 10.082 | 2832083 | 100 | -0.23 |
| 14 | 4403.6152 | 306.0253 | U | 4403.6162 | 9.655 | 1017645 | 100 | -0.23 |
| 13 | 4097.5899 | 306.0253 | Unconverted ψ | 4097.5897 | 9.281 | 2438044 | 100 | 0.05 |
| 12 | 3791.5646 | 329.0525 | A | 3791.5638 | 9.613 | 6450776 | 100 | 0.21 |
| 11 | 3462.5121 | 305.0413 | C | 3462.511 | 8.533 | 2959433 | 100 | 0.32 |
| 10 | 3157.4708 | 305.0413 | C | 3157.4687 | 8.247 | 4281684 | 100 | 0.67 |
| 9 | 2852.4295 | 329.0525 | A | 2852.4279 | 8.384 | 6732016 | 100 | 0.56 |
| 8 | 2523.3770 | 306.0253 | U | 2523.3752 | 7.06 | 3639095 | 100 | 0.71 |
| 7 | 2217.3517 | 306.0253 | U | 2217.3496 | 6.547 | 5142524 | 100 | 0.95 |
| 6 | 1911.3264 | 329.0525 | A | 1911.3234 | 5.628 | 148978 | 100 | 1.57 |
| 5 | 1582.2739 | 319.057 | $m^5C$ | 1582.271 | 4.694 | 2365111 | 100 | 1.83 |
| 4 | 1263.2169 | 306.0253 | U | 1263.216 | 1.392 | 1025750 | 100 | 0.71 |
| 3 | 957.1916 | 345.0474 | G | 957.1909 | 1.354 | 1030368 | 100 | 0.73 |
| 2 | 612.1442 | 329.0525 | A | 612.1432 | 1.334 | 609338 | 100 | 1.63 |
| 1 | 283.0917 | 345.0475 | G | NA* | NA | NA | NA | NA |

* NA: Not Analyzed. The 350 Da threshold was set to minimize background ions from the elution buffers. Otherwise, we would predominantly detect HFIP and DPA ions. Thus, the masses which are smaller than 350 Da were not detected.

**Table S9.** LC/MS analysis of a 1 ψ-containing RNA #12 (mass ladder components with CMC-converted ψ from 5′ to 3′, 20 nt RNA)

| Theoretical | | | | Extracted data file after LC/MS analysis | | | | Error |
|---|---|---|---|---|---|---|---|---|
| **Fragments** | **Theoretical mass** | **Base mass** | **Base** | **MFE mass** | **$t_R$** | **Volume** | **Quality Score** | **ppm** |
| 20 | 6597.1025 | 265.0811 | G | 6597.1125 | 13.985 | 60627484 | 100 | -1.52 |
| 19 | 6332.0214 | 329.0525 | A | 6332.0201 | 13.979 | 1541470 | 100 | 0.21 |
| 18 | 6002.9689 | 345.0474 | G | 6002.9756 | 13.816 | 2147847 | 89 | -1.12 |
| 17 | 5657.9215 | 306.0253 | U | 5657.9243 | 13.742 | 2608610 | 100 | -0.49 |
| 16 | 5351.8962 | 319.057 | $m^5C$ | 5351.8960 | 13.695 | 2110248 | 100 | 0.04 |
| 15 | 5032.8392 | 329.0525 | A | 5032.8400 | 13.633 | 1907945 | 100 | -0.16 |
| 14 | 4703.7867 | 306.0253 | U | 4703.7861 | 13.394 | 4110706 | 88 | 0.13 |
| 13 | 4397.7614 | 306.0253 | U | 4397.7599 | 13.320 | 2867370 | 100 | 0.34 |
| 12 | 4091.7361 | 329.0526 | A | 4091.7361 | 13.283 | 1855682 | 100 | 0.00 |
| 11 | 3762.6835 | 305.0413 | C | 3762.6830 | 12.962 | 2817838 | 100 | 0.13 |
| 10 | 3457.6422 | 305.0412 | C | 3457.6396 | 12.878 | 1149319 | 100 | 0.75 |
| 9 | 3152.6010 | 329.0526 | A | 3152.5974 | 12.934 | 746862 | 100 | 1.14 |
| 8 | 2823.5485 | 557.2251 | Converted ψ | 2823.5455 | 12.380 | 2149383 | 100 | 1.06 |
| 7 | 2266.3234 | 306.0253 | U | 2266.3213 | 7.944 | 4767282 | 100 | 0.93 |
| 6 | 1960.2981 | 345.0474 | G | 1960.2956 | 7.360 | 3433416 | 100 | 1.28 |
| 5 | 1615.2507 | 305.0413 | C | 1615.2481 | 6.694 | 4174772 | 100 | 1.61 |
| 4 | 1310.2094 | 305.0413 | C | 1310.2071 | 5.917 | 806139 | 87 | 1.76 |
| 3 | 1005.1681 | 329.0525 | A | 1005.1655 | 4.416 | 913589 | 100 | 2.59 |
| 2 | 676.1156 | 329.0525 | A | 676.1140 | 3.321 | 743305 | 100 | 2.37 |
| 1 | 347.0631 | 329.0525 | A | NA* | NA | NA | NA | NA |

* NA: Not Analyzed. The 350 Da threshold was set to minimize background ions from the elution buffers. Otherwise, we would predominantly detect HFIP and DPA ions. Thus, the masses which are smaller than 350 Da were not detected.

**Table S10.** LC/MS analysis of a 1 ψ-containing RNA #12 (mass ladder components with CMC-converted ψ from 3′ to 5′, RNA #12)

| Theoretical | | | | Extracted data file after LC/MS analysis | | | | Error |
|---|---|---|---|---|---|---|---|---|
| Fragments | Theoretical mass | Base mass | Base | MFE mass | $t_R$ | Volume | Quality Score | ppm |
| 20 | 6597.1025 | 329.0525 | A | 6597.1125 | 13.985 | 60627484 | 100 | -1.52 |
| 19 | 6268.0500 | 329.0525 | A | 6268.0571 | 13.936 | 2514888 | 96 | -1.13 |
| 18 | 5938.9975 | 329.0525 | A | 5939.0021 | 13.618 | 919334 | 80 | -0.77 |
| 17 | 5609.9450 | 305.0413 | C | 5609.9509 | 13.027 | 550752 | 100 | -1.05 |
| 16 | 5304.9037 | 305.0413 | C | 5304.9018 | 12.95 | 1145236 | 100 | 0.36 |
| 15 | 4999.8624 | 345.0475 | G | 4999.8628 | 13.09 | 1603456 | 100 | -0.08 |
| 14 | 4654.8150 | 306.0253 | U | 4654.8165 | 12.976 | 1028627 | 100 | -0.32 |
| 13 | 4348.7897 | 557.2251 | Converted ψ | 4348.7878 | 12.747 | 1061149 | 100 | 0.44 |
| 12 | 3791.5646 | 329.0525 | A | 3791.5638 | 9.613 | 6450776 | 100 | 0.21 |
| 11 | 3462.5121 | 305.0413 | C | 3462.511 | 8.533 | 2959433 | 100 | 0.32 |
| 10 | 3157.4708 | 305.0413 | C | 3157.4687 | 8.247 | 4281684 | 100 | 0.67 |
| 9 | 2852.4295 | 329.0525 | A | 2852.4279 | 8.384 | 6732016 | 100 | 0.56 |
| 8 | 2523.3770 | 306.0253 | U | 2523.3752 | 7.06 | 3639095 | 100 | 0.71 |
| 7 | 2217.3517 | 306.0253 | U | 2217.3496 | 6.547 | 5142524 | 100 | 0.95 |
| 6 | 1911.3264 | 329.0525 | A | 1911.3234 | 5.628 | 148978 | 100 | 1.57 |
| 5 | 1582.2739 | 319.057 | $m^5C$ | 1582.271 | 4.694 | 2365111 | 100 | 1.83 |
| 4 | 1263.2169 | 306.0253 | U | 1263.216 | 1.392 | 1025750 | 100 | 0.71 |
| 3 | 957.1916 | 345.0474 | G | 957.1909 | 1.355 | 1052036 | 100 | 0.73 |
| 2 | 612.1442 | 329.0525 | A | 612.1432 | 1.334 | 609338 | 100 | 1.63 |
| 1 | 283.0917 | 345.0475 | G | NA* | NA | NA | NA | NA |

* NA: Not Analyzed. The 350 Da threshold was set to minimize background ions from the elution buffers. Otherwise, we would predominantly detect HFIP and DPA ions. Thus, the masses which are smaller than 350 Da were not detected.

**Table S11.** LC/MS analysis of a 2 ψ-containing RNA #13 (ψ unconverted mass ladder components from 5´ to 3´, RNA #13).

| Theoretical | | | | Extracted data file after LC/MS analysis | | | | Error |
|---|---|---|---|---|---|---|---|---|
| **Fragments** | **Theoretical mass** | **Base mass** | **Base** | **MFE mass** | **$t_R$** | **Volume** | **Quality Score** | **ppm** |
| 20 | 6331.8871 | 265.0811 | G | 6331.9010 | 11.627 | 20815662 | 100 | -2.20 |
| 19 | 6066.8060 | 329.0525 | A | 6066.8121 | 11.661 | 1640168 | 99 | -1.01 |
| 18 | 5737.7535 | 345.0474 | G | 5737.7570 | 11.382 | 885613 | 80 | -0.61 |
| 17 | 5392.7061 | 306.0253 | Unconverted ψ | 5392.7060 | 11.212 | 617277 | 100 | 0.02 |
| 16 | 5086.6808 | 305.0413 | C | 5086.6829 | 11.082 | 2141353 | 100 | -0.41 |
| 15 | 4781.6395 | 329.0525 | A | 4781.6267 | 10.759 | 26031 | 75 | 2.68 |
| 14 | 4452.5870 | 306.0253 | U | 4452.5872 | 10.522 | 3256295 | 100 | -0.04 |
| 13 | 4146.5617 | 306.0253 | U | 4146.5608 | 10.294 | 2867802 | 100 | 0.22 |
| 12 | 3840.5364 | 329.0526 | A | 3840.5345 | 10.089 | 1804456 | 100 | 0.49 |
| 11 | 3511.4838 | 305.0413 | C | 3511.4825 | 9.545 | 3618243 | 100 | 0.37 |
| 10 | 3206.4425 | 305.0412 | C | 3206.4408 | 9.254 | 2325449 | 100 | 0.53 |
| 9 | 2901.4013 | 329.0526 | A | 2901.3978 | 8.965 | 1647914 | 100 | 1.21 |
| 8 | 2572.3487 | 306.0253 | Unconverted ψ | 2572.3461 | 8.205 | 3697493 | 100 | 1.01 |
| 7 | 2266.3234 | 306.0253 | U | 2266.3205 | 7.822 | 3317588 | 100 | 1.28 |
| 6 | 1960.2981 | 345.0474 | G | 1960.2952 | 7.245 | 2415197 | 100 | 1.48 |
| 5 | 1615.2507 | 305.0413 | C | 1615.2480 | 6.605 | 2827204 | 100 | 1.67 |
| 4 | 1310.2094 | 305.0413 | C | 1310.2060 | 5.804 | 1306273 | 80 | 2.60 |
| 3 | 1005.1681 | 329.0525 | A | 1005.1658 | 4.496 | 867786 | 100 | 2.29 |
| 2 | 676.1156 | 329.0525 | A | 676.1140 | 3.231 | 662092 | 100 | 2.37 |
| 1 | 347.0630 | 329.0525 | A | NA* | NA | NA | NA | NA |

* NA: Not Analyzed. The 350 Da threshold was set to minimize background ions from the elution buffers. Otherwise, we would predominantly detect HFIP and DPA ions. Thus, the masses which are smaller than 350 Da were not detected.

**Table S12.** LC/MS analysis of a 2 ψ-containing RNA #13 (mass ladder components with 1 CMC-converted ψ from 5´ to 3´, 20 nt RNA #13).

| Theoretical | | | | Extracted data file after LC/MS analysis | | | | Error |
|---|---|---|---|---|---|---|---|---|
| **Fragments** | **Theoretical mass** | **Base mass** | **Base** | **MFE mass** | **$t_R$** | **Volume** | **Quality Score** | **ppm** |
| 20 | 6583.0869 | 265.0811 | G | 6583.0981 | 13.829 | 35962424 | 100 | -1.70 |
| 19 | 6318.0058 | 329.0525 | A | 6318.0114 | 13.829 | 938044 | 100 | -0.89 |
| 18 | 5988.9533 | 345.0474 | G | 5988.9552 | 13.654 | 602824 | 96 | -0.32 |
| 17 | 5643.9059 | 306.0253 | Unconverted ψ | 5643.9107 | 13.573 | 1578612 | 80 | -0.85 |
| 16 | 5337.8806 | 305.0413 | C | 5337.8852 | 13.573 | 1563724 | 100 | -0.86 |
| 15 | 5032.8393 | 329.0525 | A | 5032.8468 | 13.541 | 991863 | 100 | -1.49 |
| 14 | 4703.7868 | 306.0253 | U | 4703.7876 | 13.308 | 1970261 | 100 | -0.17 |
| 13 | 4397.7615 | 306.0253 | U | 4397.7601 | 13.230 | 817755 | 100 | 0.32 |
| 12 | 4091.7362 | 329.0526 | A | 4091.7338 | 13.190 | 330683 | 98 | 0.59 |
| 11 | 3762.6836 | 305.0413 | C | 3762.6827 | 12.884 | 1591068 | 100 | 0.24 |
| 10 | 3457.6423 | 305.0412 | C | 3457.6403 | 12.806 | 1110204 | 99 | 0.58 |
| 9 | 3152.6011 | 329.0526 | A | 3152.5988 | 12.857 | 512332 | 100 | 0.73 |
| 8 | 2823.5485 | 557.2251 | Converted ψ | 2823.5457 | 12.325 | 1193480 | 100 | 0.99 |
| 7 | 2266.3234 | 306.0253 | U | 2266.3205 | 7.822 | 3317588 | 100 | 1.28 |
| 6 | 1960.2981 | 345.0474 | G | 1960.2952 | 7.245 | 2415197 | 100 | 1.48 |
| 5 | 1615.2507 | 305.0413 | C | 1615.2480 | 6.605 | 2827204 | 100 | 1.67 |
| 4 | 1310.2094 | 305.0413 | C | 1310.206 | 5.804 | 1306273 | 80 | 2.60 |
| 3 | 1005.1681 | 329.0525 | A | 1005.1658 | 4.496 | 867786 | 100 | 2.29 |
| 2 | 676.1156 | 329.0525 | A | 676.1140 | 3.231 | 662092 | 100 | 2.37 |
| 1 | 347.0630 | 329.0525 | A | NA* | NA | NA | NA | NA |

* NA: Not Analyzed. The 350 Da threshold was set to minimize background ions from the elution buffers. Otherwise, we would predominantly detect HFIP and DPA ions. Thus, the masses which are smaller than 350 Da were not detected.

**Table S13.** LC/MS analysis of a 2 ψ-containing RNA #13 (mass ladder components with 1 CMC-converted ψ from 5´ to 3´, RNA #13).

| Theoretical | | | | Extracted data file after LC/MS analysis | | | | Error |
|---|---|---|---|---|---|---|---|---|
| **Fragments** | **Theoretical mass** | **Base mass** | **Base** | **MFE mass** | **$t_R$** | **Volume** | **Quality Score** | **ppm** |
| 20 | 6583.0869 | 265.0811 | G | 6583.0981 | 13.829 | 35962424 | 100 | -1.70 |
| 19 | 6318.0058 | 329.0525 | A | 6318.0114 | 13.829 | 938044 | 100 | -0.89 |
| 18 | 5988.9533 | 345.0474 | G | 5988.9552 | 13.654 | 602824 | 96 | -0.32 |
| 17 | 5643.9059 | 557.2251 | Converted ψ | 5643.9107 | 13.573 | 1578612 | 80 | -0.85 |
| 16 | 5086.6808 | 305.0413 | C | 5086.6827 | 11.08 | 1427810 | 100 | -0.37 |
| 15 | 4781.6395 | 329.0525 | A | 4781.6412 | 10.926 | 1523517 | 100 | -0.36 |
| 14 | 4452.587 | 306.0253 | U | 4452.588 | 10.522 | 2085205 | 100 | -0.22 |
| 13 | 4146.5617 | 306.0253 | U | 4146.5609 | 10.294 | 2788426 | 100 | 0.19 |
| 12 | 3840.5364 | 329.0526 | A | 3840.5345 | 10.084 | 1938977 | 100 | 0.49 |
| 11 | 3511.4838 | 305.0413 | C | 3511.4816 | 9.546 | 3088818 | 100 | 0.63 |
| 10 | 3206.4425 | 305.0412 | C | 3206.4409 | 9.253 | 2028277 | 100 | 0.50 |
| 9 | 2901.4013 | 329.0526 | A | 2901.3977 | 8.965 | 1489932 | 100 | 1.24 |
| 8 | 2572.3487 | 306.0253 | Unconverted ψ | 2572.3461 | 8.205 | 3716588 | 100 | 1.01 |
| 7 | 2266.3234 | 306.0253 | U | 2266.3205 | 7.822 | 3317588 | 100 | 1.28 |
| 6 | 1960.2981 | 345.0474 | G | 1960.2952 | 7.245 | 2415197 | 100 | 1.48 |
| 5 | 1615.2507 | 305.0413 | C | 1615.248 | 6.605 | 2827204 | 100 | 1.67 |
| 4 | 1310.2094 | 305.0413 | C | 1310.206 | 5.804 | 1306273 | 80 | 2.60 |
| 3 | 1005.1681 | 329.0525 | A | 1005.1658 | 4.496 | 867786 | 100 | 2.29 |
| 2 | 676.1156 | 329.0525 | A | 676.114 | 3.231 | 662092 | 100 | 2.37 |
| 1 | 347.0631 | 329.0525 | A | NA* | NA | NA | NA | NA |

* NA: Not Analyzed. The 350 Da threshold was set to minimize background ions from the elution buffers. Otherwise, we would predominantly detect HFIP and DPA ions. Thus, the masses which are smaller than 350 Da were not detected.

**Table S14.** LC/MS analysis of a 2 ψ-containing RNA #13 (mass ladder components with 2 CMC-converted ψ from 5´ to RNA #13).

| Theoretical | | | | Extracted data file after LC/MS analysis | | | | Error |
|---|---|---|---|---|---|---|---|---|
| Fragments | Theoretical mass | Base mass | Base | MFE mass | $t_R$ | Volume | Quality Score | ppm |
| 20 | 6834.2866 | 265.0811 | G | 6834.2945 | 15.887 | 10647840 | 100 | -1.16 |
| 19 | 6569.2055 | 329.0525 | A | 6569.2283 | 15.694 | 22547 | 73 | -3.47 |
| 18 | 6240.1530 | 345.0474 | G | 6241.1635 | 15.787 | 151235 | 79 | -161.94 |
| 17 | 5895.1056 | 557.2251 | Converted ψ | 5895.0646 | 15.870 | 3373 | 53 | 6.95 |
| 16 | 5337.8805 | 305.0413 | C | 5337.8852 | 13.573 | 1563724 | 100 | -0.88 |
| 15 | 5032.8392 | 329.0525 | A | 5032.8468 | 13.541 | 991863 | 100 | -1.51 |
| 14 | 4703.7867 | 306.0253 | U | 4703.7876 | 13.308 | 1970261 | 100 | -0.19 |
| 13 | 4397.7614 | 306.0253 | U | 4397.7601 | 13.230 | 817755 | 100 | 0.30 |
| 12 | 4091.7361 | 329.0526 | A | 4091.7338 | 13.190 | 330683 | 98 | 0.56 |
| 11 | 3762.6835 | 305.0413 | C | 3762.6827 | 12.884 | 1591068 | 100 | 0.21 |
| 10 | 3457.6422 | 305.0412 | C | 3457.6403 | 12.806 | 1110204 | 99 | 0.55 |
| 9 | 3152.6010 | 329.0526 | A | 3152.5988 | 12.857 | 512332 | 100 | 0.70 |
| 8 | 2823.5484 | 557.2251 | Converted ψ | 2823.5457 | 12.325 | 1193480 | 100 | 0.96 |
| 7 | 2266.3233 | 306.0253 | U | 2266.3205 | 7.822 | 3317588 | 100 | 1.24 |
| 6 | 1960.2980 | 345.0474 | G | 1960.2952 | 7.245 | 2415197 | 100 | 1.43 |
| 5 | 1615.2506 | 305.0413 | C | 1615.248 | 6.605 | 2827204 | 100 | 1.61 |
| 4 | 1310.2093 | 305.0413 | C | 1310.206 | 5.804 | 1306273 | 80 | 2.52 |
| 3 | 1005.1680 | 329.0525 | A | 1005.1658 | 4.496 | 867786 | 100 | 2.19 |
| 2 | 676.1155 | 329.0525 | A | 676.1140 | 3.231 | 662092 | 100 | 2.22 |
| 1 | 347.0630 | 329.0525 | A | NA* | NA | NA | NA | NA |

* NA: Not Analyzed. The 350 Da threshold was set to minimize background ions from the elution buffers. Otherwise, we would predominantly detect HFIP and DPA ions. Thus, the masses which are smaller than 350 Da were not detected.

**Table S15.** LC/MS analysis of 3´-biotin-labeled RNA #1, showing its mass ladder components.

| Theoretical | | | | Extracted data file after LC/MS analysis | | | | Error |
|---|---|---|---|---|---|---|---|---|
| Fragments | Theoretical mass | Base mass | Base | MFE mass | $t_R$ | Volume | Quality Score | ppm |
| 19 | 6781.0733 | 305.0413 | C | 6781.0413 | 9.752 | 16819442 | 100 | 4.72 |
| 18 | 6476.0320 | 345.0474 | G | 6475.9924 | 9.717 | 247965 | 84 | 6.11 |
| 17 | 6130.9846 | 305.0413 | C | 6130.9398 | 9.662 | 178841 | 80 | 7.31 |
| 16 | 5825.9433 | 329.0525 | A | 5825.9037 | 9.782 | 510096 | 80 | 6.80 |
| 15 | 5496.8908 | 306.0253 | U | 5496.8566 | 9.383 | 262486 | 99 | 6.22 |
| 14 | 5190.8655 | 305.0413 | C | 5190.8364 | 9.241 | 349988 | 100 | 5.61 |
| 13 | 4885.8242 | 306.0253 | U | 4885.7908 | 9.135 | 356118 | 100 | 6.84 |
| 12 | 4579.7989 | 345.0475 | G | 4579.7738 | 9.109 | 386687 | 100 | 5.48 |
| 11 | 4234.7514 | 329.0525 | A | 4234.7271 | 9.145 | 305380 | 100 | 5.74 |
| 10 | 3905.6989 | 305.0413 | C | 3905.6749 | 8.575 | 145505 | 96 | 6.14 |
| 9 | 3600.6576 | 306.0253 | U | 3600.6373 | 8.420 | 195308 | 100 | 5.64 |
| 8 | 3294.6323 | 345.0474 | G | 3294.6165 | 8.370 | 125991 | 100 | 4.80 |
| 7 | 2949.5849 | 329.0525 | A | 2949.5716 | 8.339 | 106993 | 100 | 4.51 |
| 6 | 2620.5324 | 305.0413 | C | 2620.5193 | 7.492 | 90629 | 100 | 5.00 |
| 5 | 2315.4911 | 305.0413 | C | 2315.4814 | 7.299 | 163692 | 100 | 4.19 |
| 4 | 2010.4498 | 329.0525 | A | 2010.4388 | 7.625 | 279963 | 100 | 5.47 |
| 3 | 1681.3973 | 329.0525 | A | 1681.3891 | 7.354 | 183827 | 100 | 4.88 |
| 2 | 1352.3448 | 329.0526 | A | 1352.3378 | 7.303 | 135065 | 100 | 5.18 |
| 1 | 1023.2922 | 329.0525 | A | 1023.2859 | 7.219 | 106700 | 100 | 6.16 |

**Table S16.** LC/MS analysis of 3´-biotin-labeled RNA #2, showing its mass ladder components.

| Theoretical | | | | Extracted data file after LC/MS analysis | | | | Error |
|---|---|---|---|---|---|---|---|---|
| **Fragments** | **Theoretical mass** | **Base mass** | **Base** | **MFE mass** | **$t_R$** | **Volume** | **Quality Score** | **ppm** |
| 20 | 7079.0823 | 329.2088 | A | 7079.0519 | 9.695 | 15887400 | 100 | 4.29 |
| 19 | 6750.0298 | 306.1667 | U | 6749.9576 | 9.422 | 103400 | 80 | 10.70 |
| 18 | 6444.0045 | 329.2088 | A | 6443.9541 | 9.504 | 292394 | 91 | 7.82 |
| 17 | 6114.9519 | 345.2077 | G | 6114.9026 | 9.156 | 99684 | 87 | 8.06 |
| 16 | 5769.9045 | 305.1828 | C | 5769.8585 | 9.020 | 146499 | 80 | 7.97 |
| 15 | 5464.8632 | 305.1828 | C | 5464.8200 | 8.887 | 63438 | 80 | 7.91 |
| 14 | 5159.8219 | 305.1827 | C | 5159.8026 | 8.769 | 284881 | 100 | 3.74 |
| 13 | 4854.7806 | 329.2088 | A | 4854.7562 | 8.879 | 336079 | 100 | 5.03 |
| 12 | 4525.7281 | 345.2078 | G | 4525.7034 | 8.413 | 242815 | 100 | 5.46 |
| 11 | 4180.6807 | 306.1667 | U | 4180.6582 | 8.181 | 208097 | 100 | 5.38 |
| 10 | 3874.6554 | 305.1828 | C | 3874.6356 | 7.962 | 274449 | 100 | 5.11 |
| 9 | 3569.6141 | 329.2087 | A | 3569.5960 | 8.083 | 385282 | 100 | 5.07 |
| 8 | 3240.5616 | 345.2078 | G | 3240.5467 | 7.440 | 238714 | 100 | 4.60 |
| 7 | 2895.5141 | 306.1668 | U | 2895.4995 | 7.096 | 215938 | 100 | 5.04 |
| 6 | 2589.4888 | 305.1827 | C | 2589.4766 | 6.736 | 291557 | 100 | 4.71 |
| 5 | 2284.4476 | 306.1668 | U | 2284.4371 | 6.523 | 322833 | 100 | 4.60 |
| 4 | 1978.4223 | 329.2088 | A | 1978.4135 | 6.382 | 360972 | 100 | 4.45 |
| 3 | 1649.3697 | 305.1827 | C | 1649.3626 | 5.238 | 129210 | 100 | 4.30 |
| 2 | 1344.3284 | 345.2078 | G | 1344.3224 | 5.191 | 163260 | 100 | 4.46 |
| 1 | 999.2810 | 305.1827 | C | 999.2753 | 5.323 | 81388 | 100 | 5.70 |

**Table S17.** LC/MS analysis of 3´-biotin-labeled RNA #3, showing its mass ladder components.

| Theoretical | | | | Extracted data file after LC/MS analysis | | | | Error |
|---|---|---|---|---|---|---|---|---|
| **Fragments** | **Theoretical mass** | **Base mass** | **Base** | **MFE mass** | **$t_R$** | **Volume** | **Quality Score** | **ppm** |
| 20 | 7088.0826 | 329.0525 | A | 7088.0481 | 9.912 | 20041130 | 100 | 4.87 |
| 19 | 6759.0301 | 329.0525 | A | 6758.9787 | 9.827 | 312216 | 85 | 7.60 |
| 18 | 6429.9776 | 329.0525 | A | 6429.9267 | 9.575 | 270720 | 80 | 7.92 |
| 17 | 6100.9251 | 305.0413 | C | 6100.8781 | 9.174 | 239340 | 80 | 7.70 |
| 16 | 5795.8838 | 305.0413 | C | 5795.8548 | 9.071 | 488843 | 100 | 5.00 |
| 15 | 5490.8425 | 345.0475 | G | 5490.8073 | 9.044 | 673490 | 98 | 6.41 |
| 14 | 5145.7950 | 306.0253 | U | 5145.7622 | 8.944 | 583546 | 100 | 6.37 |
| 13 | 4839.7697 | 306.0253 | U | 4839.7411 | 8.870 | 671098 | 100 | 5.91 |
| 12 | 4533.7444 | 329.0525 | A | 4533.7210 | 8.874 | 1044860 | 100 | 5.16 |
| 11 | 4204.6919 | 305.0413 | C | 4204.6731 | 8.297 | 513780 | 100 | 4.47 |
| 10 | 3899.6506 | 305.0413 | C | 3899.6292 | 8.185 | 650568 | 100 | 5.49 |
| 9 | 3594.6093 | 329.0525 | A | 3594.5921 | 8.321 | 1203072 | 100 | 4.78 |
| 8 | 3265.5568 | 306.0253 | U | 3265.5424 | 7.668 | 797335 | 100 | 4.41 |
| 7 | 2959.5315 | 306.0253 | U | 2959.5198 | 7.482 | 1166317 | 100 | 3.95 |
| 6 | 2653.5062 | 329.0525 | A | 2653.4961 | 7.449 | 1461689 | 100 | 3.81 |
| 5 | 2324.4537 | 305.0413 | C | 2324.4451 | 6.497 | 759285 | 100 | 3.70 |
| 4 | 2019.4124 | 306.0253 | U | 2019.4051 | 6.113 | 869967 | 100 | 3.61 |
| 3 | 1713.3871 | 345.0474 | G | 1713.3810 | 5.978 | 955386 | 100 | 3.56 |
| 2 | 1368.3397 | 329.0525 | A | 1368.3338 | 6.370 | 586922 | 100 | 4.31 |
| 1 | 1039.2872 | 1039.2872 | G | 1039.2825 | 5.660 | 342316 | 100 | 4.52 |

**Table S18.** LC/MS analysis of 3´-biotin-labeled RNA #4, showing its mass ladder components.

| Theoretical | | | | Extracted data file after LC/MS analysis | | | | Error |
|---|---|---|---|---|---|---|---|---|
| **Fragments** | **Theoretical mass** | **Base mass** | **Base** | **MFE mass** | **$t_R$** | **Volume** | **Quality Score** | **ppm** |
| 20 | 6985.0431 | 306.0253 | U | 6985.0207 | 11.625 | 58498820 | 100 | 3.21 |
| 19 | 6679.0178 | 345.0474 | G | 6679.9864 | 11.557 | 78870 | 100 | -145.02 |
| 18 | 6333.9704 | 306.0253 | U | 6333.9577 | 11.514 | 1165403 | 95 | 2.01 |
| 17 | 6027.9451 | 329.0526 | A | 6027.9150 | 11.707 | 3055438 | 100 | 4.99 |
| 16 | 5698.8925 | 329.0525 | A | 5698.8699 | 11.588 | 2305641 | 85 | 3.97 |
| 15 | 5369.8400 | 329.0525 | A | 5369.8145 | 11.201 | 1931925 | 100 | 4.75 |
| 14 | 5040.7875 | 305.0413 | C | 5040.7605 | 10.777 | 1506142 | 100 | 5.36 |
| 13 | 4735.7462 | 329.0525 | A | 4735.7232 | 11.042 | 3132367 | 100 | 4.86 |
| 12 | 4406.6937 | 306.0253 | U | 4406.6725 | 10.372 | 1761089 | 100 | 4.81 |
| 11 | 4100.6684 | 305.0413 | C | 4100.6501 | 10.171 | 2219510 | 100 | 4.46 |
| 10 | 3795.6271 | 305.0413 | C | 3795.6100 | 10.043 | 2529132 | 100 | 4.51 |
| 9 | 3490.5858 | 306.0253 | U | 3490.5716 | 10.035 | 2441434 | 100 | 4.07 |
| 8 | 3184.5605 | 329.0525 | A | 3184.5476 | 10.052 | 3440631 | 100 | 4.05 |
| 7 | 2855.5080 | 305.0413 | C | 2855.4962 | 9.308 | 1722723 | 100 | 4.13 |
| 6 | 2550.4667 | 329.0525 | A | 2550.4587 | 9.605 | 2447222 | 98 | 3.14 |
| 5 | 2221.4142 | 305.0413 | C | 2221.4058 | 8.474 | 1901654 | 100 | 3.78 |
| 4 | 1916.3729 | 306.0253 | U | 1916.3661 | 8.222 | 2469329 | 100 | 3.55 |
| 3 | 1610.3476 | 305.0413 | C | 1610.3419 | 7.922 | 2259370 | 100 | 3.54 |
| 2 | 1305.3063 | 306.0253 | U | 1305.3016 | 7.899 | 1603980 | 100 | 3.60 |
| 1 | 999.2810 | 305.0413 | C | 999.2770 | 8.131 | 1272190 | 100 | 4.00 |

**Table S19.** LC/MS analysis of 3´-biotin-labeled RNA #5, showing its mass ladder components.

| Theoretical | | | | Extracted data file after LC/MS analysis | | | | Error |
|---|---|---|---|---|---|---|---|---|
| **Fragments** | **Theoretical mass** | **Base mass** | **Base** | **MFE mass** | $t_R$ | **Volume** | **Quality Score** | **ppm** |
| 20 | 7073.0717 | 306.0253 | U | 7073.0472 | 12.156 | 82887552 | 100 | 3.46 |
| 19 | 6767.0464 | 329.0525 | A | 6767.0137 | 12.296 | 4576892 | 100 | 4.83 |
| 18 | 6437.9939 | 306.0253 | U | 6437.9769 | 11.923 | 2315485 | 100 | 2.64 |
| 17 | 6131.9686 | 306.0253 | U | 6131.9364 | 11.837 | 3353132 | 100 | 5.25 |
| 16 | 5825.9433 | 305.0413 | C | 5825.9117 | 11.751 | 3225051 | 100 | 5.42 |
| 15 | 5520.9020 | 329.0525 | A | 5520.8693 | 11.976 | 4163809 | 100 | 5.92 |
| 14 | 5191.8495 | 329.0525 | A | 5191.8215 | 11.751 | 3066109 | 100 | 5.39 |
| 13 | 4862.7970 | 345.0475 | G | 4862.7739 | 11.305 | 2677901 | 100 | 4.75 |
| 12 | 4517.7495 | 306.0253 | U | 4517.7279 | 11.153 | 2051199 | 100 | 4.78 |
| 11 | 4211.7242 | 306.0253 | U | 4211.7051 | 11.108 | 3646647 | 100 | 4.53 |
| 10 | 3905.6989 | 329.0525 | A | 3905.6825 | 11.163 | 4185511 | 100 | 4.20 |
| 9 | 3576.6464 | 305.0413 | C | 3576.6311 | 10.626 | 2134080 | 100 | 4.28 |
| 8 | 3271.6051 | 329.0525 | A | 3271.5909 | 10.892 | 4157558 | 100 | 4.34 |
| 7 | 2942.5526 | 305.0413 | C | 2942.5413 | 10.114 | 2150759 | 100 | 3.84 |
| 6 | 2637.5113 | 306.0253 | U | 2637.5010 | 9.986 | 2806597 | 100 | 3.91 |
| 5 | 2331.4860 | 305.0413 | C | 2331.4714 | 10.257 | 10052 | 100 | 6.26 |
| 4 | 2026.4447 | 329.0525 | A | 2026.4377 | 10.176 | 3408728 | 100 | 3.45 |
| 3 | 1697.3922 | 329.0525 | A | 1697.3861 | 9.691 | 2143607 | 100 | 3.59 |
| 2 | 1368.3397 | 345.0475 | G | 1368.3344 | 9.292 | 1254041 | 100 | 3.87 |
| 1 | 1023.2922 | 329.0525 | A | 1023.2882 | 10.603 | 1407833 | 100 | 3.91 |

**Table S20.** LC/MS analysis of 3´-biotin-labeled RNA #6, showing its mass ladder components.

| Theoretical | | | | Extracted data file after LC/MS analysis | | | | Error |
|---|---|---|---|---|---|---|---|---|
| **Fragments** | **Theoretical mass** | **Base mass** | **Base** | **MFE mass** | **$t_R$** | **Volume** | **Quality Score** | **ppm** |
| 20 | 6954.9836 | 345.0475 | G | 6954.9496 | 9.252 | 19518152 | 100 | 4.89 |
| 19 | 6609.9361 | 305.0412 | C | 6609.8895 | 9.140 | 186061 | 80 | 7.05 |
| 18 | 6304.8949 | 345.0475 | G | 6304.8493 | 9.116 | 570614 | 95 | 7.23 |
| 17 | 5959.8474 | 306.0253 | U | 5959.8016 | 9.064 | 430830 | 100 | 7.68 |
| 16 | 5653.8221 | 329.0525 | A | 5653.7951 | 9.068 | 845499 | 100 | 4.78 |
| 15 | 5324.7696 | 305.0413 | C | 5324.7375 | 8.714 | 497200 | 100 | 6.03 |
| 14 | 5019.7283 | 329.0525 | A | 5019.7206 | 8.862 | 69267 | 80 | 1.53 |
| 13 | 4690.6758 | 306.0253 | U | 4690.6508 | 8.360 | 624270 | 100 | 5.33 |
| 12 | 4384.6505 | 305.0413 | C | 4384.6287 | 8.202 | 905111 | 100 | 4.97 |
| 11 | 4079.6092 | 306.0253 | U | 4079.5872 | 8.088 | 934627 | 100 | 5.39 |
| 10 | 3773.5839 | 306.0253 | U | 3773.5610 | 7.898 | 865362 | 100 | 6.07 |
| 9 | 3467.5586 | 305.0413 | C | 3467.5420 | 7.648 | 551801 | 100 | 4.79 |
| 8 | 3162.5173 | 305.0413 | C | 3162.5033 | 7.427 | 763065 | 100 | 4.43 |
| 7 | 2857.4760 | 305.0413 | C | 2857.4632 | 7.176 | 934459 | 100 | 4.48 |
| 6 | 2552.4347 | 305.0412 | C | 2552.4237 | 6.942 | 1266516 | 100 | 4.31 |
| 5 | 2247.3935 | 306.0253 | U | 2247.3841 | 6.711 | 1457982 | 100 | 4.18 |
| 4 | 1941.3682 | 306.0253 | U | 1941.3606 | 6.369 | 1784912 | 100 | 3.91 |
| 3 | 1635.3429 | 306.0254 | U | 1635.3358 | 6.162 | 1549510 | 100 | 4.34 |
| 2 | 1329.3175 | 329.0525 | A | 1329.3122 | 6.619 | 1621370 | 100 | 3.99 |
| 1 | 1000.2650 | 306.0253 | U | 1000.2615 | 5.284 | 24083 | 100 | 3.50 |

**Table S21.** LC/MS analysis of 3´-biotin-labeled RNA #7, showing its mass ladder components.

| Theoretical | | | | Extracted data file after LC/MS analysis | | | | Error |
|---|---|---|---|---|---|---|---|---|
| Fragments | Theoretical mass | Base mass | Base | MFE mass | $t_R$ | Volume | Quality Score | ppm |
| 20 | 7110.0881 | 305.0413 | C | 7110.0622 | 11.550 | 40533036 | 100 | 3.64 |
| 19 | 6805.0468 | 345.0474 | G | 6805.0163 | 11.536 | 1377944 | 100 | 4.48 |
| 18 | 6459.9994 | 305.0413 | C | 6459.9655 | 11.383 | 515259 | 96 | 5.25 |
| 17 | 6154.9581 | 305.0413 | C | 6154.9267 | 11.333 | 915022 | 100 | 5.10 |
| 16 | 5849.9168 | 329.0525 | A | 5849.8891 | 11.425 | 2491248 | 99 | 4.74 |
| 15 | 5520.8643 | 306.0253 | U | 5520.8364 | 10.963 | 957615 | 100 | 5.05 |
| 14 | 5214.8390 | 345.0475 | G | 5214.8129 | 10.913 | 1607534 | 100 | 5.00 |
| 13 | 4869.7915 | 306.0253 | U | 4869.7663 | 10.706 | 1002213 | 100 | 5.17 |
| 12 | 4563.7662 | 345.0474 | G | 4563.7450 | 10.786 | 872578 | 100 | 4.65 |
| 11 | 4218.7188 | 329.0525 | A | 4218.6990 | 10.933 | 1284822 | 100 | 4.69 |
| 10 | 3889.6663 | 306.0253 | U | 3889.6549 | 10.212 | 786209 | 100 | 2.93 |
| 9 | 3583.6410 | 305.0413 | C | 3583.6265 | 9.978 | 940944 | 100 | 4.05 |
| 8 | 3278.5997 | 305.0413 | C | 3278.5866 | 9.685 | 809912 | 100 | 4.00 |
| 7 | 2973.5584 | 305.0413 | C | 2973.5474 | 9.381 | 679854 | 100 | 3.70 |
| 6 | 2668.5171 | 345.0474 | G | 2668.5070 | 9.315 | 819030 | 100 | 3.78 |
| 5 | 2323.4697 | 345.0474 | G | 2323.4614 | 9.329 | 646645 | 100 | 3.57 |
| 4 | 1978.4223 | 329.0526 | A | 1978.4141 | 9.272 | 715798 | 100 | 4.14 |
| 3 | 1649.3697 | 305.0413 | C | 1649.3622 | 7.894 | 182946 | 100 | 4.55 |
| 2 | 1344.3284 | 305.0412 | C | 1344.3226 | 7.901 | 369846 | 100 | 4.31 |
| 1 | 1039.2872 | 345.0475 | G | 1039.2824 | 8.816 | 397016 | 100 | 4.62 |

**Table S22.** LC/MS analysis of 3´-biotin-labeled RNA #8, showing its mass ladder components.

| Theoretical | | | | Extracted data file after LC/MS analysis | | | | Error |
|---|---|---|---|---|---|---|---|---|
| Fragments | Theoretical mass | Base mass | Base | MFE mass | $t_R$ | Volume | Quality Score | ppm |
| 20 | 7151.1160 | 329.0525 | A | 7151.0928 | 12.317 | 87850496 | 100 | 3.24 |
| 19 | 6822.0635 | 305.0413 | C | 6822.0257 | 12.046 | 836640 | 100 | 5.54 |
| 18 | 6517.0222 | 329.0526 | A | 6516.9906 | 12.178 | 1896420 | 100 | 4.85 |
| 17 | 6187.9696 | 305.0412 | C | 6187.9538 | 11.973 | 51293 | 98 | 2.55 |
| 16 | 5882.9284 | 306.0253 | U | 5882.8973 | 11.690 | 2436562 | 100 | 5.29 |
| 15 | 5576.9031 | 345.0475 | G | 5576.8745 | 11.763 | 2954102 | 100 | 5.13 |
| 14 | 5231.8556 | 329.0525 | A | 5231.8307 | 11.780 | 1503563 | 100 | 4.76 |
| 13 | 4902.8031 | 305.0413 | C | 4902.7787 | 11.376 | 1728477 | 100 | 4.98 |
| 12 | 4597.7618 | 329.0525 | A | 4597.7384 | 11.440 | 3528610 | 100 | 5.09 |
| 11 | 4268.7093 | 306.0253 | U | 4268.6887 | 10.855 | 1721343 | 100 | 4.83 |
| 10 | 3962.6840 | 345.0474 | G | 3962.6651 | 10.805 | 2353609 | 100 | 4.77 |
| 9 | 3617.6366 | 345.0475 | G | 3617.6199 | 10.832 | 1863580 | 100 | 4.62 |
| 8 | 3272.5891 | 329.0525 | A | 3272.5764 | 10.649 | 230927 | 100 | 3.88 |
| 7 | 2943.5366 | 305.0413 | C | 2943.5235 | 10.040 | 1417986 | 100 | 4.45 |
| 6 | 2638.4953 | 306.0253 | U | 2638.4844 | 9.867 | 2035557 | 100 | 4.13 |
| 5 | 2332.4700 | 345.0474 | G | 2332.4613 | 9.878 | 2467172 | 100 | 3.73 |
| 4 | 1987.4226 | 329.0525 | A | 1987.4147 | 10.359 | 2158002 | 100 | 3.97 |
| 3 | 1658.3701 | 329.0526 | A | 1658.3625 | 9.410 | 70871 | 100 | 4.58 |
| 2 | 1329.3175 | 306.0253 | U | 1329.3130 | 8.639 | 37300 | 100 | 3.39 |
| 1 | 1023.2922 | 329.0525 | A | 1023.2883 | 10.597 | 1731424 | 100 | 3.81 |

**Table S23.** LC/MS analysis of 3´-biotin-labeled RNA #9, showing its mass ladder components.

| Theoretical | | | | Extracted data file after LC/MS analysis | | | | Error |
|---|---|---|---|---|---|---|---|---|
| Fragments | Theoretical mass | Base mass | Base | MFE mass | $t_R$ | Volume | Quality Score | ppm |
| 20 | 7193.0524 | 345.0474 | G | 7193.0274 | 11.807 | 55442324 | 100 | 3.48 |
| 19 | 6848.0050 | 305.0413 | C | 6847.9696 | 11.629 | 605416 | 99 | 5.17 |
| 18 | 6542.9637 | 345.0474 | G | 6542.9287 | 11.612 | 1153241 | 100 | 5.35 |
| 17 | 6197.9163 | 345.0475 | G | 6197.8868 | 11.627 | 1710951 | 100 | 4.76 |
| 16 | 5852.8688 | 329.0525 | A | 5852.8355 | 11.750 | 1889983 | 100 | 5.69 |
| 15 | 5523.8163 | 306.0253 | U | 5523.7916 | 11.276 | 1055262 | 100 | 4.47 |
| 14 | 5217.7910 | 306.0253 | U | 5217.7646 | 11.181 | 2644440 | 100 | 5.06 |
| 13 | 4911.7657 | 306.0253 | U | 4911.7562 | 11.195 | 2901850 | 100 | 1.93 |
| 12 | 4605.7404 | 329.0525 | A | 4605.7117 | 10.639 | 54327 | 100 | 6.23 |
| 11 | 4276.6879 | 345.0474 | G | 4276.6684 | 12.237 | 1747514 | 100 | 4.56 |
| 10 | 3931.6405 | 305.0413 | C | 3931.6227 | 10.370 | 1744474 | 100 | 4.53 |
| 9 | 3626.5992 | 306.0253 | U | 3626.5834 | 10.080 | 2028011 | 100 | 4.36 |
| 8 | 3320.5739 | 305.0413 | C | 3320.5607 | 9.905 | 1675877 | 100 | 3.98 |
| 7 | 3015.5326 | 329.0525 | A | 3015.5209 | 10.128 | 2926950 | 100 | 3.88 |
| 6 | 2686.4801 | 345.0475 | G | 2686.4700 | 9.355 | 1768713 | 100 | 3.76 |
| 5 | 2341.4326 | 306.0253 | U | 2341.4237 | 8.811 | 1667926 | 100 | 3.80 |
| 4 | 2035.4073 | 306.0253 | U | 2035.3998 | 8.419 | 1823836 | 100 | 3.68 |
| 3 | 1729.3820 | 345.0474 | G | 1729.3764 | 8.342 | 1574679 | 100 | 3.24 |
| 2 | 1384.3346 | 345.0474 | G | 1384.3290 | 8.383 | 897954 | 100 | 4.05 |
| 1 | 1039.2872 | 345.0475 | G | 1039.2827 | 8.811 | 725527 | 100 | 4.33 |

**Table S24.** LC/MS analysis of 3´-biotin-labeled RNA #10, showing its mass ladder components.

| Theoretical | | | | Extracted data file after LC/MS analysis | | | | Error |
|---|---|---|---|---|---|---|---|---|
| Fragments | Theoretical mass | Base mass | Base | MFE mass | $t_R$ | Volume | Quality Score | ppm |
| 20 | 7088.0826 | 305.0413 | C | 7088.0613 | 11.883 | 83257784 | 100 | 3.01 |
| 19 | 6783.0413 | 329.0525 | A | 6783.0061 | 11.975 | 2374953 | 100 | 5.19 |
| 18 | 6453.9888 | 305.0413 | C | 6453.9468 | 11.681 | 1388931 | 100 | 6.51 |
| 17 | 6148.9475 | 329.0525 | A | 6148.9140 | 11.935 | 1819504 | 100 | 5.45 |
| 16 | 5819.8950 | 329.0525 | A | 5819.8674 | 11.838 | 1894041 | 100 | 4.74 |
| 15 | 5490.8425 | 329.0525 | A | 5490.8152 | 11.586 | 2817326 | 100 | 4.97 |
| 14 | 5161.7900 | 306.0253 | U | 5161.7648 | 11.083 | 2176473 | 100 | 4.88 |
| 13 | 4855.7647 | 306.0253 | U | 4855.7413 | 10.915 | 3237261 | 100 | 4.82 |
| 12 | 4549.7394 | 305.0413 | C | 4549.7141 | 10.730 | 2960106 | 100 | 5.56 |
| 11 | 4244.6981 | 345.0475 | G | 4244.6787 | 10.741 | 3118826 | 100 | 4.57 |
| 10 | 3899.6506 | 345.0474 | G | 3899.6401 | 10.625 | 2939016 | 100 | 2.69 |
| 9 | 3554.6032 | 306.0253 | U | 3554.5892 | 10.396 | 2535213 | 100 | 3.94 |
| 8 | 3248.5779 | 306.0253 | U | 3248.5652 | 9.955 | 114648 | 100 | 3.91 |
| 7 | 2942.5526 | 305.0413 | C | 2942.5417 | 9.980 | 2735803 | 100 | 3.70 |
| 6 | 2637.5113 | 306.0253 | U | 2637.5011 | 9.974 | 2936338 | 100 | 3.87 |
| 5 | 2331.4860 | 329.0525 | A | 2331.4784 | 9.985 | 2893702 | 100 | 3.26 |
| 4 | 2002.4335 | 305.0413 | C | 2002.4268 | 10.028 | 41002 | 94 | 3.35 |
| 3 | 1697.3922 | 329.0525 | A | 1697.3866 | 9.786 | 3139447 | 100 | 3.30 |
| 2 | 1368.3397 | 329.0525 | A | 1368.3343 | 9.551 | 1604226 | 100 | 3.95 |
| 1 | 1039.2872 | 345.0475 | G | 1039.2827 | 8.817 | 981598 | 100 | 4.33 |

**Table S25.** LC/MS analysis of 3´-biotin-labeled RNA #11, showing its mass ladder components.

| Theoretical | | | | Extracted data file after LC/MS analysis | | | | Error |
|---|---|---|---|---|---|---|---|---|
| **Fragments** | **Theoretical mass** | **Base mass** | **Base** | **MFE mass** | **$t_R$** | **Volume** | **Quality Score** | **ppm** |
| 21 | 7522.1050 | 345.0475 | G | 7522.0727 | 9.677 | 10294695 | 100 | 4.29 |
| 20 | 7177.0575 | 305.0413 | C | 7176.9932 | 9.555 | 20884 | 74 | 8.96 |
| 19 | 6872.0162 | 345.0474 | G | 6871.9425 | 9.511 | 77150 | 80 | 10.72 |
| 18 | 6526.9688 | 345.0474 | G | 6526.9038 | 9.494 | 106806 | 97 | 9.96 |
| 17 | 6181.9214 | 329.0526 | A | 6181.8843 | 9.576 | 410798 | 93 | 6.00 |
| 16 | 5852.8688 | 306.0253 | U | 5852.8373 | 9.197 | 106865 | 90 | 5.38 |
| 15 | 5546.8435 | 306.0253 | U | 5546.8112 | 9.073 | 412694 | 98 | 5.82 |
| 14 | 5240.8182 | 306.0253 | U | 5240.7832 | 8.977 | 298557 | 99 | 6.68 |
| 13 | 4934.7929 | 329.0525 | A | 4934.7688 | 9.053 | 289020 | 100 | 4.88 |
| 12 | 4605.7404 | 345.0474 | G | 4605.7156 | 8.603 | 217621 | 100 | 5.38 |
| 11 | 4260.6930 | 305.0413 | C | 4260.6680 | 8.429 | 242965 | 100 | 5.87 |
| 10 | 3955.6517 | 306.0253 | U | 3955.6316 | 8.244 | 345563 | 100 | 5.08 |
| 9 | 3649.6264 | 305.0413 | C | 3649.6065 | 8.024 | 410186 | 100 | 5.45 |
| 8 | 3344.5851 | 329.0525 | A | 3344.5699 | 8.115 | 552137 | 100 | 4.54 |
| 7 | 3015.5326 | 345.0474 | G | 3015.5116 | 7.460 | 373904 | 100 | 6.96 |
| 6 | 2670.4852 | 306.0253 | U | 2670.4729 | 7.068 | 332059 | 100 | 4.61 |
| 5 | 2364.4599 | 306.0253 | U | 2364.4490 | 6.658 | 358553 | 100 | 4.61 |
| 4 | 2058.4346 | 345.0475 | G | 2058.4263 | 6.345 | 313197 | 100 | 4.03 |
| 3 | 1713.3871 | 345.0474 | G | 1713.3799 | 6.069 | 197197 | 100 | 4.20 |
| 2 | 1368.3397 | 345.0475 | G | 1368.3325 | 6.214 | 146520 | 100 | 5.26 |
| 1 | 1023.2922 | 329.0525 | A | 1023.2863 | 7.215 | 150107 | 100 | 5.77 |

**Table S26.** LC/MS analysis of 5´-sulfo-Cy3-labeled RNA #1, showing its mass ladder components.

| Theoretical | | | | Extracted data file after LC/MS analysis | | | | Error |
|---|---|---|---|---|---|---|---|---|
| **Fragments** | **Theoretical mass** | **Base mass** | **Base** | **MFE mass** | **$t_R$** | **Volume** | **Quality Score** | **ppm** |
| 19 | 6859.0606 | 249.0862 | A | 6859.0293 | 13.278 | 29162746 | 100 | 4.56 |
| 18 | 6609.9744 | 329.0525 | A | 6610.9354 | 13.116 | 69218 | 73 | -145.39 |
| 17 | 6280.9219 | 329.0525 | A | 6280.8859 | 13.138 | 299442 | 89 | 5.73 |
| 16 | 5951.8694 | 329.0525 | A | 5951.8447 | 13.077 | 150172 | 80 | 4.15 |
| 15 | 5622.8169 | 305.0413 | C | 5622.7793 | 12.955 | 260581 | 80 | 6.69 |
| 14 | 5317.7756 | 305.0413 | C | 5318.7555 | 13.012 | 19172 | 68 | -184.27 |
| 13 | 5012.7343 | 329.0525 | A | 5012.7020 | 12.996 | 242326 | 94 | 6.44 |
| 12 | 4683.6818 | 345.0475 | G | 4683.6584 | 12.898 | 685126 | 87 | 5.00 |
| 11 | 4338.6343 | 306.0253 | U | 4338.6115 | 12.875 | 640041 | 100 | 5.26 |
| 10 | 4032.6090 | 305.0413 | C | 4032.5867 | 12.881 | 306999 | 96 | 5.53 |
| 9 | 3727.5677 | 329.0525 | A | 3727.5518 | 12.967 | 86034 | 81 | 4.27 |
| 8 | 3398.5152 | 345.0474 | G | 3398.5004 | 12.795 | 1050778 | 99 | 4.35 |
| 7 | 3053.4678 | 306.0253 | U | 3053.4581 | 12.691 | 33763 | 83 | 3.18 |
| 6 | 2747.4425 | 305.0413 | C | 2747.4301 | 12.803 | 244796 | 80 | 4.51 |
| 5 | 2442.4012 | 306.0253 | U | 2442.3910 | 12.791 | 1013984 | 100 | 4.18 |
| 4 | 2136.3759 | 329.0525 | A | 2136.3676 | 12.769 | 184183 | 87 | 3.89 |
| 3 | 1807.3234 | 305.0413 | C | 1807.3165 | 12.770 | 1840549 | 100 | 3.82 |
| 2 | 1502.2821 | 345.0474 | G | 1502.2765 | 12.794 | 965042 | 100 | 3.73 |
| 1 | 1157.2347 | 305.0413 | C | 1157.2297 | 13.642 | 913331 | 100 | 4.32 |

**Table S27.** LC/MS analysis of 5´-sulfo-Cy3-labeled RNA #2, showing its mass ladder components.

| Theoretical | | | | Extracted data file after LC/MS analysis | | | | Error |
|---|---|---|---|---|---|---|---|---|
| Fragments | Theoretical mass | Base mass | Base | MFE mass | $t_R$ | Volume | Quality Score | ppm |
| 20 | 7157.0696 | 225.0750 | C | 7157.0369 | 13.199 | 18994320 | 100 | 4.57 |
| 19 | 6931.9946 | 345.0474 | G | 6931.0867 | 13.257 | 166222 | 80 | 130.97 |
| 18 | 6586.9472 | 305.0413 | C | 6586.9163 | 13.247 | 373645 | 80 | 4.69 |
| 17 | 6281.9059 | 329.0525 | A | 6281.8475 | 13.280 | 216088 | 80 | 9.30 |
| 16 | 5952.8534 | 306.0253 | U | 5952.8248 | 13.205 | 931428 | 100 | 4.80 |
| 15 | 5646.8281 | 305.0413 | C | 5646.8065 | 13.222 | 627329 | 98 | 3.83 |
| 14 | 5341.7868 | 306.0253 | U | 5341.7543 | 13.240 | 209385 | 80 | 6.08 |
| 13 | 5035.7615 | 345.0474 | G | 5035.7355 | 13.256 | 355370 | 80 | 5.16 |
| 12 | 4690.7141 | 329.0526 | A | 4690.6877 | 13.288 | 293771 | 97 | 5.63 |
| 11 | 4361.6615 | 305.0412 | C | 4361.6393 | 13.183 | 624454 | 100 | 5.09 |
| 10 | 4056.6203 | 306.0253 | U | 4056.5940 | 13.154 | 22971 | 79 | 6.48 |
| 9 | 3750.5950 | 345.0475 | G | 3750.5764 | 13.218 | 392405 | 97 | 4.96 |
| 8 | 3405.5475 | 329.0525 | A | 3405.5311 | 13.266 | 376785 | 96 | 4.82 |
| 7 | 3076.4950 | 305.0413 | C | 3076.4812 | 13.144 | 764082 | 95 | 4.49 |
| 6 | 2771.4537 | 305.0413 | C | 2771.4461 | 13.182 | 576176 | 100 | 2.74 |
| 5 | 2466.4124 | 305.0413 | C | 2466.4028 | 13.212 | 258560 | 100 | 3.89 |
| 4 | 2161.3711 | 345.0474 | G | 2161.3628 | 13.277 | 548722 | 80 | 3.84 |
| 3 | 1816.3237 | 329.0525 | A | 1816.3169 | 13.474 | 783483 | 84 | 3.74 |
| 2 | 1487.2712 | 306.0253 | U | 1487.2656 | 13.532 | 1797103 | 100 | 3.77 |
| 1 | 1181.2459 | 329.0525 | A | 1181.2408 | 13.861 | 824092 | 100 | 4.32 |

**Table S28.** LC/MS analysis of 5´-sulfo-Cy3-labeled RNA #3, showing its mass ladder components.

| Theoretical | | | | Extracted data file after LC/MS analysis | | | | Error |
|---|---|---|---|---|---|---|---|---|
| Fragments | Theoretical mass | Base mass | Base | MFE mass | $t_R$ | Volume | Quality Score | ppm |
| 20 | 7166.0699 | 265.0811 | G | 7166.0350 | 13.464 | 15752947 | 100 | 4.87 |
| 19 | 6900.9888 | 329.0525 | A | 6900.9425 | 13.428 | 275366 | 82 | 6.71 |
| 18 | 6571.9363 | 345.0474 | G | 6571.8939 | 13.356 | 132733 | 74 | 6.45 |
| 17 | 6226.8889 | 306.0253 | U | 6226.8593 | 13.354 | 180552 | 78 | 4.75 |
| 16 | 5920.8636 | 305.0413 | C | 5920.8086 | 13.403 | 212136 | 80 | 9.29 |
| 15 | 5615.8223 | 329.0525 | A | 5615.7902 | 13.426 | 260478 | 80 | 5.72 |
| 14 | 5286.7698 | 306.0253 | U | 5286.7436 | 13.348 | 876722 | 91 | 4.96 |
| 13 | 4980.7445 | 306.0253 | U | 4980.7386 | 13.371 | 654236 | 100 | 1.18 |
| 12 | 4674.7192 | 329.0526 | A | 4674.6993 | 13.424 | 542251 | 80 | 4.26 |
| 11 | 4345.6666 | 305.0413 | C | 4345.6466 | 13.329 | 814417 | 100 | 4.60 |
| 10 | 4040.6253 | 305.0412 | C | 4040.6052 | 13.361 | 520867 | 98 | 4.97 |
| 9 | 3735.5841 | 329.0526 | A | 3735.5739 | 13.419 | 42982 | 59 | 2.73 |
| 8 | 3406.5315 | 306.0253 | U | 3406.5151 | 13.318 | 770893 | 96 | 4.81 |
| 7 | 3100.5062 | 306.0253 | U | 3100.4930 | 13.340 | 491826 | 100 | 4.26 |
| 6 | 2794.4809 | 345.0474 | G | 2794.4683 | 13.348 | 371969 | 94 | 4.51 |
| 5 | 2449.4335 | 305.0413 | C | 2449.4233 | 13.398 | 303466 | 80 | 4.16 |
| 4 | 2144.3922 | 305.0413 | C | 2144.3829 | 13.436 | 419905 | 87 | 4.34 |
| 3 | 1839.3509 | 329.0525 | A | 1839.3429 | 13.365 | 179583 | 86 | 4.35 |
| 2 | 1510.2984 | 329.0525 | A | 1510.2924 | 13.403 | 288879 | 80 | 3.97 |
| 1 | 1181.2459 | 329.0525 | A | 1181.2410 | 13.860 | 707398 | 100 | 4.15 |

**Table S29.** LC/MS analysis of 5´-sulfo-Cy3-labeled RNA #4, showing its mass ladder components.

| Theoretical | | | | Extracted data file after LC/MS analysis | | | | Error |
|---|---|---|---|---|---|---|---|---|
| Fragments | Theoretical mass | Base mass | Base | MFE mass | $t_R$ | Volume | Quality Score | ppm |
| 20 | 7063.0304 | 225.0749 | C | 7063.0072 | 13.390 | 11257376 | 87 | 3.28 |
| 19 | 6837.9555 | 306.0253 | U | 6837.9201 | 13.469 | 300823 | 86 | 5.18 |
| 18 | 6531.9302 | 305.0413 | C | 6531.9373 | 13.584 | 30910 | 80 | -1.09 |
| 17 | 6226.8889 | 306.0253 | U | 6226.8376 | 13.627 | 26579 | 60 | 8.24 |
| 16 | 5920.8636 | 305.0413 | C | 5920.8443 | 13.631 | 50737 | 75 | 3.26 |
| 15 | 5615.8223 | 329.0525 | A | 5615.7920 | 13.671 | 42482 | 63 | 5.40 |
| 14 | 5286.7698 | 305.0413 | C | 5286.7615 | 13.594 | 843779 | 83 | 1.57 |
| 13 | 4981.7285 | 329.0525 | A | 4981.6999 | 13.636 | 151248 | 97 | 5.74 |
| 12 | 4652.6760 | 306.0254 | U | 4652.6511 | 13.391 | 1191688 | 87 | 5.35 |
| 11 | 4346.6506 | 305.0412 | C | 4346.6371 | 13.403 | 130923 | 70 | 3.11 |
| 10 | 4041.6094 | 305.0413 | C | 4041.5867 | 13.571 | 376672 | 93 | 5.62 |
| 9 | 3736.5681 | 306.0253 | U | 3736.5502 | 13.588 | 60297 | 97 | 4.79 |
| 8 | 3430.5428 | 329.0525 | A | 3430.5239 | 13.454 | 45199 | 69 | 5.51 |
| 7 | 3101.4903 | 305.0413 | C | 3101.4769 | 13.301 | 778223 | 99 | 4.32 |
| 6 | 2796.4490 | 329.0526 | A | 2796.4353 | 13.695 | 35158 | 78 | 4.90 |
| 5 | 2467.3964 | 329.0525 | A | 2467.3855 | 13.818 | 108974 | 88 | 4.42 |
| 4 | 2138.3439 | 329.0525 | A | 2138.3355 | 13.161 | 82910 | 88 | 3.93 |
| 3 | 1809.2914 | 306.0253 | U | 1809.2846 | 13.153 | 2193742 | 87 | 3.76 |
| 2 | 1503.2661 | 345.0474 | G | 1503.2598 | 13.708 | 159632 | 100 | 4.19 |
| 1 | 1158.2187 | 306.0253 | U | 1158.2131 | 14.188 | 1574057 | 100 | 4.84 |

**Table S30.** LC/MS analysis of 5´-sulfo-Cy3-labeled RNA #5, showing its mass ladder components.

| Theoretical | | | | Extracted data file after LC/MS analysis | | | | Error |
|---|---|---|---|---|---|---|---|---|
| Fragments | Theoretical mass | Base mass | Base | MFE mass | $t_R$ | Volume | Quality Score | ppm |
| 20 | 7151.0590 | 249.0862 | A | 7151.0256 | 13.949 | 43379424 | 100 | 4.67 |
| 19 | 6901.9728 | 345.0474 | G | 6901.9226 | 13.835 | 322593 | 80 | 7.27 |
| 18 | 6556.9254 | 329.0525 | A | 6556.8915 | 13.831 | 396493 | 80 | 5.17 |
| 17 | 6227.8729 | 329.0525 | A | 6227.8645 | 13.834 | 76663 | 79 | 1.35 |
| 16 | 5898.8204 | 305.0413 | C | 5898.7973 | 13.640 | 212566 | 70 | 3.92 |
| 15 | 5593.7791 | 306.0253 | U | 5593.7475 | 13.745 | 664008 | 80 | 5.65 |
| 14 | 5287.7538 | 305.0413 | C | 5287.7257 | 13.749 | 1458044 | 100 | 5.31 |
| 13 | 4982.7125 | 329.0525 | A | 4982.6855 | 13.742 | 174109 | 80 | 5.42 |
| 12 | 4653.6600 | 305.0413 | C | 4653.6362 | 13.697 | 2006854 | 100 | 5.11 |
| 11 | 4348.6187 | 329.0525 | A | 4348.5989 | 13.647 | 73164 | 72 | 4.55 |
| 10 | 4019.5662 | 306.0253 | U | 4019.5481 | 13.535 | 920778 | 84 | 4.50 |
| 9 | 3713.5409 | 306.0253 | U | 3713.5244 | 13.672 | 120519 | 100 | 4.44 |
| 8 | 3407.5156 | 345.0475 | G | 3407.4991 | 13.681 | 168659 | 72 | 4.84 |
| 7 | 3062.4681 | 329.0525 | A | 3062.4534 | 13.557 | 126015 | 87 | 4.80 |
| 6 | 2733.4156 | 329.0525 | A | 2733.4027 | 13.604 | 327314 | 98 | 4.72 |
| 5 | 2404.3631 | 305.0413 | C | 2404.3530 | 13.326 | 2389580 | 87 | 4.20 |
| 4 | 2099.3218 | 306.0253 | U | 2099.3075 | 13.169 | 17723 | 91 | 6.81 |
| 3 | 1793.2965 | 306.0253 | U | 1793.2897 | 13.865 | 217546 | 99 | 3.79 |
| 2 | 1487.2712 | 329.0525 | A | 1487.2657 | 13.677 | 2638249 | 100 | 3.70 |
| 1 | 1158.2187 | 306.0253 | U | 1158.2134 | 14.192 | 2172695 | 100 | 4.58 |

**Table S31.** LC/MS analysis of 5´-sulfo-Cy3-labeled RNA #6, showing its mass ladder components.

| Theoretical | | | | Extracted data file after LC/MS analysis | | | | Error |
|---|---|---|---|---|---|---|---|---|
| Fragments | Theoretical mass | Base mass | Base | MFE mass | $t_R$ | Volume | Quality Score | ppm |
| 20 | 7032.9709 | 226.0590 | U | 7032.9380 | 12.938 | 24081534 | 100 | 4.68 |
| 19 | 6806.9119 | 329.0525 | A | 6806.8565 | 12.954 | 497938 | 100 | 8.14 |
| 18 | 6477.8594 | 306.0253 | U | 6477.8296 | 12.875 | 1123636 | 100 | 4.60 |
| 17 | 6171.8341 | 306.0253 | U | 6171.7982 | 12.889 | 797659 | 100 | 5.82 |
| 16 | 5865.8088 | 306.0253 | U | 5865.8484 | 12.899 | 1419968 | 80 | -6.75 |
| 15 | 5559.7835 | 305.0413 | C | 5559.7761 | 12.919 | 249723 | 80 | 1.33 |
| 14 | 5254.7422 | 305.0413 | C | 5254.7165 | 12.944 | 1499456 | 100 | 4.89 |
| 13 | 4949.7009 | 305.0413 | C | 4949.6783 | 12.982 | 147053 | 79 | 4.57 |
| 12 | 4644.6596 | 305.0413 | C | 4644.6354 | 13.000 | 1219024 | 100 | 5.21 |
| 11 | 4339.6183 | 306.0253 | U | 4339.6137 | 13.021 | 1246558 | 100 | 1.06 |
| 10 | 4033.5930 | 306.0253 | U | 4033.5760 | 13.029 | 1640640 | 100 | 4.21 |
| 9 | 3727.5677 | 305.0412 | C | 3727.5530 | 13.039 | 726317 | 96 | 3.94 |
| 8 | 3422.5265 | 306.0253 | U | 3422.5122 | 13.068 | 1753331 | 100 | 4.18 |
| 7 | 3116.5012 | 329.0526 | A | 3116.4876 | 13.113 | 1248491 | 100 | 4.36 |
| 6 | 2787.4486 | 305.0413 | C | 2787.4373 | 12.970 | 2163746 | 95 | 4.05 |
| 5 | 2482.4073 | 329.0525 | A | 2482.3979 | 13.002 | 695135 | 100 | 3.79 |
| 4 | 2153.3548 | 306.0253 | U | 2153.3470 | 12.883 | 2141185 | 100 | 3.62 |
| 3 | 1847.3295 | 345.0474 | G | 1847.3226 | 12.935 | 1062104 | 100 | 3.74 |
| 2 | 1502.2821 | 305.0413 | C | 1502.2770 | 13.140 | 2211201 | 100 | 3.39 |
| 1 | 1197.2408 | 345.0474 | G | 1197.2362 | 13.279 | 1324255 | 98 | 3.84 |

**Table S32.** LC/MS analysis of 5´-sulfo-Cy3-labeled RNA #7, showing its mass ladder components.

| Theoretical | | | | Extracted data file after LC/MS analysis | | | | Error |
|---|---|---|---|---|---|---|---|---|
| Fragments | Theoretical mass | Base mass | Base | MFE mass | $t_R$ | Volume | Quality Score | ppm |
| 20 | 7188.0754 | 265.0811 | G | 7188.0577 | 13.257 | 198372 | 70 | 2.46 |
| 19 | 6922.9943 | 305.0413 | C | 6922.9600 | 13.374 | 1169126 | 80 | 4.95 |
| 18 | 6617.9530 | 305.0413 | C | 6617.9032 | 13.372 | 360353 | 74 | 7.52 |
| 17 | 6312.9117 | 329.0525 | A | 6312.8754 | 13.386 | 707713 | 80 | 5.75 |
| 16 | 5983.8592 | 345.0474 | G | 5983.8242 | 13.343 | 112885 | 77 | 5.85 |
| 15 | 5638.8118 | 345.0475 | G | 5638.7821 | 13.268 | 961515 | 80 | 5.27 |
| 14 | 5293.7643 | 305.0412 | C | 5293.7168 | 13.185 | 35206 | 75 | 8.97 |
| 13 | 4988.7231 | 305.0413 | C | 4988.7064 | 13.196 | 35019 | 80 | 3.35 |
| 12 | 4683.6818 | 305.0413 | C | 4683.6599 | 13.355 | 148461 | 76 | 4.68 |
| 11 | 4378.6405 | 306.0253 | U | 4378.6236 | 13.355 | 51270 | 73 | 3.86 |
| 10 | 4072.6152 | 329.0525 | A | 4072.5932 | 13.368 | 444401 | 80 | 5.40 |
| 9 | 3743.5627 | 345.0475 | G | 3743.5471 | 13.261 | 227634 | 87 | 4.17 |
| 8 | 3398.5152 | 306.0253 | U | 3398.4868 | 13.177 | 17855 | 60 | 8.36 |
| 7 | 3092.4899 | 345.0474 | G | 3092.4781 | 13.125 | 168338 | 100 | 3.82 |
| 6 | 2747.4425 | 306.0253 | U | 2747.4316 | 13.187 | 1180398 | 80 | 3.97 |
| 5 | 2441.4172 | 329.0525 | A | 2441.4095 | 13.120 | 42956 | 69 | 3.15 |
| 4 | 2112.3647 | 305.0413 | C | 2112.3571 | 13.052 | 1527354 | 100 | 3.60 |
| 3 | 1807.3234 | 305.0413 | C | 1807.3165 | 13.069 | 1451369 | 100 | 3.82 |
| 2 | 1502.2821 | 345.0474 | G | 1502.2772 | 13.207 | 113774 | 68 | 3.26 |
| 1 | 1157.2347 | 305.0413 | C | 1157.2301 | 13.961 | 766397 | 100 | 3.97 |

**Table S33.** LC/MS analysis of 5´-sulfo-Cy3-labeled RNA #8, showing its mass ladder components.

| Theoretical | | | | Extracted data file after LC/MS analysis | | | | Error |
|---|---|---|---|---|---|---|---|---|
| Fragments | Theoretical mass | Base mass | Base | MFE mass | $t_R$ | Volume | Quality Score | ppm |
| 20 | 7229.1033 | 249.0862 | A | 7229.0695 | 14.003 | 20040654 | 100 | 4.68 |
| 19 | 6980.0171 | 306.0253 | U | 6979.9807 | 13.902 | 342469 | 93 | 5.21 |
| 18 | 6673.9918 | 329.0525 | A | 6673.9395 | 13.923 | 96589 | 80 | 7.84 |
| 17 | 6344.9393 | 329.0525 | A | 6344.8887 | 13.883 | 446012 | 100 | 7.97 |
| 16 | 6015.8868 | 345.0475 | G | 6015.8539 | 13.811 | 789692 | 100 | 5.47 |
| 15 | 5670.8393 | 306.0253 | U | 5670.8112 | 13.810 | 791636 | 100 | 4.96 |
| 14 | 5364.8140 | 305.0413 | C | 5364.7851 | 13.819 | 362044 | 80 | 5.39 |
| 13 | 5059.7727 | 329.0525 | A | 5059.7461 | 13.868 | 339561 | 94 | 5.26 |
| 12 | 4730.7202 | 345.0474 | G | 4730.6953 | 13.791 | 747218 | 100 | 5.26 |
| 11 | 4385.6728 | 345.0475 | G | 4385.6481 | 13.785 | 214489 | 94 | 5.63 |
| 10 | 4040.6253 | 306.0253 | U | 4040.6034 | 13.783 | 610851 | 96 | 5.42 |
| 9 | 3734.6000 | 329.0525 | A | 3734.5797 | 13.829 | 119982 | 80 | 5.44 |
| 8 | 3405.5475 | 305.0413 | C | 3405.5304 | 13.722 | 821756 | 100 | 5.02 |
| 7 | 3100.5062 | 329.0525 | A | 3100.4915 | 13.818 | 232602 | 97 | 4.74 |
| 6 | 2771.4537 | 345.0474 | G | 2771.4408 | 13.716 | 597795 | 98 | 4.65 |
| 5 | 2426.4063 | 306.0253 | U | 2426.3956 | 13.699 | 984832 | 100 | 4.41 |
| 4 | 2120.3810 | 305.0413 | C | 2120.3722 | 13.781 | 756259 | 100 | 4.15 |
| 3 | 1815.3397 | 329.0525 | A | 1815.3334 | 13.920 | 202800 | 70 | 3.47 |
| 2 | 1486.2872 | 305.0413 | C | 1486.2813 | 13.960 | 1082970 | 100 | 3.97 |
| 1 | 1181.2459 | 329.0525 | A | 1181.2406 | 14.173 | 183863 | 100 | 4.49 |

**Table S34.** LC/MS analysis of 5´-sulfo-Cy3-labeled RNA #9, showing its mass ladder components.

| Theoretical | | | | Extracted data file after LC/MS analysis | | | | Error |
|---|---|---|---|---|---|---|---|---|
| Fragments | Theoretical mass | Base mass | Base | MFE mass | $t_R$ | Volume | Quality Score | ppm |
| 20 | 7271.0397 | 265.0811 | G | 7271.0150 | 13.447 | 60392476 | 100 | 3.40 |
| 19 | 7005.9586 | 345.0474 | G | 7005.9069 | 13.414 | 1544260 | 80 | 7.38 |
| 18 | 6660.9112 | 345.0474 | G | 6660.8417 | 13.401 | 115234 | 80 | 10.43 |
| 17 | 6315.8638 | 306.0253 | U | 6315.8150 | 13.385 | 1000227 | 99 | 7.73 |
| 16 | 6009.8385 | 306.0253 | U | 6009.8074 | 13.404 | 2545935 | 100 | 5.17 |
| 15 | 5703.8132 | 345.0475 | G | 5703.7940 | 13.412 | 2410664 | 100 | 3.37 |
| 14 | 5358.7657 | 329.0525 | A | 5358.7424 | 13.432 | 2729923 | 100 | 4.35 |
| 13 | 5029.7132 | 305.0413 | C | 5029.6903 | 13.335 | 4588952 | 100 | 4.55 |
| 12 | 4724.6719 | 306.0253 | U | 4724.6524 | 13.354 | 3608892 | 100 | 4.13 |
| 11 | 4418.6466 | 305.0413 | C | 4418.6275 | 13.370 | 2676034 | 100 | 4.32 |
| 10 | 4113.6053 | 345.0474 | G | 4113.5871 | 13.360 | 2671523 | 100 | 4.42 |
| 9 | 3768.5579 | 329.0525 | A | 3768.5431 | 13.376 | 2388710 | 100 | 3.93 |
| 8 | 3439.5054 | 306.0253 | U | 3439.4913 | 13.239 | 5653201 | 100 | 4.10 |
| 7 | 3133.4801 | 306.0253 | U | 3133.4702 | 13.243 | 5267381 | 100 | 3.16 |
| 6 | 2827.4548 | 306.0253 | U | 2827.4447 | 13.246 | 5120720 | 100 | 3.57 |
| 5 | 2521.4295 | 329.0525 | A | 2521.4201 | 13.262 | 2676447 | 100 | 3.73 |
| 4 | 2192.3770 | 345.0475 | G | 2192.3695 | 13.169 | 4164446 | 100 | 3.42 |
| 3 | 1847.3295 | 345.0474 | G | 1847.3232 | 13.196 | 3096228 | 100 | 3.41 |
| 2 | 1502.2821 | 305.0413 | C | 1502.7018 | 13.404 | 105885 | 100 | -279.37 |
| 1 | 1197.2408 | 345.0474 | G | 1197.2365 | 13.559 | 2206497 | 100 | 3.59 |

**Table S35.** LC/MS analysis of 5´-sulfo-Cy3-labeled RNA #10, showing its mass ladder components.

| Theoretical | | | | Extracted data file after LC/MS analysis | | | | Error |
|---|---|---|---|---|---|---|---|---|
| Fragments | Theoretical mass | Base mass | Base | MFE mass | $t_R$ | Volume | Quality Score | ppm |
| 20 | 7166.0699 | 265.0811 | G | 7166.0414 | 13.739 | 69199560 | 100 | 3.98 |
| 19 | 6900.9888 | 329.0525 | A | 6900.9627 | 13.715 | 919150 | 84 | 3.78 |
| 18 | 6571.9363 | 329.0525 | A | 6571.8812 | 13.658 | 1047891 | 80 | 8.38 |
| 17 | 6242.8838 | 305.0413 | C | 6242.8539 | 13.597 | 1775042 | 87 | 4.79 |
| 16 | 5937.8425 | 329.0525 | A | 5937.8328 | 13.633 | 1623713 | 100 | 1.63 |
| 15 | 5608.7900 | 306.0253 | U | 5608.7668 | 13.540 | 3247803 | 84 | 4.14 |
| 14 | 5302.7647 | 305.0413 | C | 5302.7398 | 13.580 | 2133663 | 80 | 4.70 |
| 13 | 4997.7234 | 306.0253 | U | 4997.6996 | 13.526 | 95112 | 99 | 4.76 |
| 12 | 4691.6981 | 306.0253 | U | 4691.6768 | 13.611 | 2450965 | 100 | 4.54 |
| 11 | 4385.6728 | 345.0475 | G | 4385.6522 | 13.605 | 1676478 | 95 | 4.70 |
| 10 | 4040.6253 | 345.0474 | G | 4040.6081 | 13.541 | 840397 | 87 | 4.26 |
| 9 | 3695.5779 | 305.0413 | C | 3695.5619 | 13.592 | 2706026 | 100 | 4.33 |
| 8 | 3390.5366 | 306.0253 | U | 3390.5202 | 13.605 | 2811611 | 80 | 4.84 |
| 7 | 3084.5113 | 306.0253 | U | 3084.5013 | 13.627 | 2928850 | 80 | 3.24 |
| 6 | 2778.4860 | 329.0525 | A | 2778.4757 | 13.655 | 1989313 | 80 | 3.71 |
| 5 | 2449.4335 | 329.0525 | A | 2449.4242 | 13.610 | 2067865 | 95 | 3.80 |
| 4 | 2120.3810 | 329.0525 | A | 2120.3722 | 13.528 | 1449232 | 80 | 4.15 |
| 3 | 1791.3285 | 305.0413 | C | 1791.3228 | 13.583 | 1030070 | 87 | 3.18 |
| 2 | 1486.2872 | 329.0525 | A | 1486.2816 | 13.482 | 1294548 | 81 | 3.77 |
| 1 | 1157.2347 | 305.0413 | C | 1157.2299 | 14.136 | 1103912 | 87 | 4.15 |

**Table S36.** LC/MS analysis of 5´-sulfo-Cy3-labeled RNA #11, showing its mass ladder components.

| Theoretical | | | | Extracted data file after LC/MS analysis | | | | Error |
|---|---|---|---|---|---|---|---|---|
| **Fragments** | **Theoretical mass** | **Base mass** | **Base** | **MFE mass** | **$t_R$** | **Volume** | **Quality Score** | **ppm** |
| 21 | 7600.0923 | 249.0862 | A | 7600.0602 | 13.263 | 21803014 | 100 | 4.22 |
| 20 | 7351.0061 | 345.0475 | G | 7350.9572 | 13.133 | 208063 | 80 | 6.65 |
| 19 | 7005.9586 | 345.0474 | G | 7007.9075 | 13.126 | 20219 | 85 | -278.18 |
| 18 | 6660.9112 | 345.0474 | G | 6660.8639 | 13.117 | 272418 | 80 | 7.10 |
| 17 | 6315.8638 | 306.0253 | U | 6315.8230 | 13.105 | 213624 | 80 | 6.46 |
| 16 | 6009.8385 | 306.0253 | U | 6009.7925 | 13.119 | 469394 | 80 | 7.65 |
| 15 | 5703.8132 | 345.0475 | G | 5703.7807 | 13.125 | 307370 | 100 | 5.70 |
| 14 | 5358.7657 | 329.0525 | A | 5358.7543 | 13.143 | 797008 | 99 | 2.13 |
| 13 | 5029.7132 | 305.0413 | C | 5029.6880 | 13.054 | 1304776 | 100 | 5.01 |
| 12 | 4724.6719 | 306.0253 | U | 4724.6479 | 13.077 | 822977 | 100 | 5.08 |
| 11 | 4418.6466 | 305.0413 | C | 4418.6277 | 13.098 | 935202 | 100 | 4.28 |
| 10 | 4113.6053 | 345.0474 | G | 4113.5865 | 13.091 | 823731 | 100 | 4.57 |
| 9 | 3768.5579 | 329.0525 | A | 3768.5416 | 13.108 | 903026 | 100 | 4.33 |
| 8 | 3439.5054 | 306.0253 | U | 3439.4924 | 12.970 | 1748702 | 100 | 3.78 |
| 7 | 3133.4801 | 306.0253 | U | 3133.4698 | 12.975 | 1760722 | 100 | 3.29 |
| 6 | 2827.4548 | 306.0253 | U | 2827.4439 | 12.980 | 1762939 | 100 | 3.86 |
| 5 | 2521.4295 | 329.0525 | A | 2521.4208 | 12.994 | 454731 | 100 | 3.45 |
| 4 | 2192.3770 | 345.0475 | G | 2192.3692 | 12.904 | 1509385 | 100 | 3.56 |
| 3 | 1847.3295 | 345.0474 | G | 1847.3231 | 12.929 | 1224721 | 100 | 3.46 |
| 2 | 1502.2821 | 305.0413 | C | 1502.2770 | 13.128 | 1429495 | 100 | 3.39 |
| 1 | 1197.2408 | 345.0474 | G | 1197.2365 | 13.271 | 832362 | 100 | 3.59 |

**Table S37.** LC/MS analysis of 3´-biotin-labeled RNA #12, showing its CMC-ψ-converted mass ladder components.

| Theoretical | | | | Extracted data file after LC/MS analysis | | | | Error |
|---|---|---|---|---|---|---|---|---|
| **Fragments** | **Theoretical mass** | **Base mass** | **Base** | **MFE mass** | $t_R$ | **Volume** | **Quality Score** | **ppm** |
| 20 | 7485.3767 | 329.0525 | A | 7485.3910 | 15.919 | 4301601 | 100 | -1.91 |
| 19 | 7156.3242 | 329.0525 | A | 7157.3265 | 15.917 | 39821 | 100 | -140.06 |
| 18 | 6827.2717 | 329.0525 | A | 6827.3170 | 15.917 | 56899 | 80 | -6.64 |
| 17 | 6498.2192 | 305.0413 | C | 6498.2258 | 15.896 | 30478 | 80 | -1.02 |
| 16 | 6193.1779 | 305.0413 | C | 6193.1808 | 15.928 | 47149 | 79 | -0.47 |
| 15 | 5888.1366 | 345.0475 | G | 5888.0997 | 15.924 | 84778 | 80 | 6.27 |
| 14 | 5543.0891 | 306.0253 | U | 5543.1180 | 15.924 | 132659 | 80 | -5.21 |
| 13 | 5237.0638 | 557.2251 | Converted ψ | 5237.0573 | 15.949 | 40639 | 80 | 1.24 |
| 12 | 4679.8387 | 329.0525 | A | 4679.8399 | 14.581 | 2275437 | 88 | -0.26 |
| 11 | 4350.7862 | 305.0413 | C | 4350.7877 | 14.104 | 356588 | 84 | -0.34 |
| 10 | 4045.7449 | 305.0413 | C | 4045.7446 | 14.070 | 158059 | 91 | 0.07 |
| 9 | 3740.7036 | 329.0525 | A | 3740.7018 | 14.437 | 797927 | 100 | 0.48 |
| 8 | 3411.6511 | 306.0253 | U | 3411.6501 | 13.988 | 281415 | 96 | 0.29 |
| 7 | 3105.6258 | 306.0253 | U | 3105.6155 | 13.593 | 11367 | 93 | 3.32 |
| 6 | 2799.6005 | 329.0525 | A | 2799.5971 | 14.370 | 30839 | 100 | 1.21 |
| 5 | 2470.5480 | 319.0570 | $m^5C$ | 2470.5463 | 14.271 | 3260900 | 100 | 0.69 |
| 4 | 2151.4910 | 306.0253 | U | 2151.4861 | 13.612 | 379983 | 79 | 2.28 |
| 3 | 1845.4657 | 345.0474 | G | 1845.4639 | 14.006 | 2888007 | 91 | 0.98 |
| 2 | 1500.4183 | 329.0525 | A | 1500.4155 | 14.916 | 8695 | 66 | 1.87 |
| 1 | 1171.3658 | 345.0475 | G | 1171.3639 | 15.168 | 1223164 | 80 | 1.62 |

**Table S38.** LC/MS analysis of 3´-biotin-labeled RNA #12, showing its ψ-unconverted mass ladder components.

| Theoretical | | | | Extracted data file after LC/MS analysis | | | | Error |
|---|---|---|---|---|---|---|---|---|
| **Fragments** | **Theoretical mass** | **Base mass** | **Base** | **MFE mass** | **$t_R$** | **Volume** | **Quality Score** | **ppm** |
| 20 | 7234.1769 | 329.0525 | A | 7234.1945 | 14.789 | 341038 | 100 | -2.43 |
| 19 | 6905.1244 | 329.0525 | A | 6905.1351 | 14.831 | 41837 | 80 | -1.55 |
| 18 | 6576.0719 | 329.0525 | A | 6576.0763 | 14.646 | 147954 | 81 | -0.67 |
| 17 | 6247.0194 | 305.0413 | C | 6247.0289 | 14.269 | 194023 | 99 | -1.52 |
| 16 | 5941.9781 | 305.0413 | C | 5941.9839 | 14.269 | 208740 | 100 | -0.98 |
| 15 | 5636.9368 | 345.0475 | G | 5636.9382 | 14.273 | 30200 | 80 | -0.25 |
| 14 | 5291.8893 | 306.0253 | U | 5291.8772 | 14.187 | 12930 | 91 | 2.29 |
| 13 | 4985.8640 | 306.0253 | Unconverted ψ | 4985.8436 | 14.236 | 19666 | 90 | 4.09 |
| 12 | 4679.8387 | 329.0525 | A | 4679.8399 | 14.581 | 2275437 | 88 | -0.26 |
| 11 | 4350.7862 | 305.0413 | C | 4350.7877 | 14.104 | 356588 | 84 | -0.34 |
| 10 | 4045.7449 | 305.0413 | C | 4045.7446 | 14.070 | 158059 | 91 | 0.07 |
| 9 | 3740.7036 | 329.0525 | A | 3740.7018 | 14.437 | 797927 | 100 | 0.48 |
| 8 | 3411.6511 | 306.0253 | U | 3411.6501 | 13.988 | 281415 | 96 | 0.29 |
| 7 | 3105.6258 | 306.0253 | U | 3105.6155 | 13.593 | 11367 | 93 | 3.32 |
| 6 | 2799.6005 | 329.0525 | A | 2799.5971 | 14.370 | 30839 | 100 | 1.21 |
| 5 | 2470.5480 | 319.0570 | $m^5C$ | 2470.5463 | 14.271 | 3260900 | 100 | 0.69 |
| 4 | 2151.4910 | 306.0253 | U | 2151.4861 | 13.612 | 379983 | 79 | 2.28 |
| 3 | 1845.4657 | 345.0474 | G | 1845.4639 | 14.006 | 2888007 | 91 | 0.98 |
| 2 | 1500.4183 | 329.0525 | A | 1500.4155 | 14.916 | 8695 | 66 | 1.87 |
| 1 | 1171.3658 | 345.0475 | G | 1171.3639 | 15.168 | 1223164 | 80 | 1.62 |

**Table S39.** LC/MS analysis of 3´-biotin-labeled RNA #3, showing its mass ladder components.

| Theoretical | | | | Extracted data file after LC/MS analysis | | | | Error |
|---|---|---|---|---|---|---|---|---|
| Fragments | Theoretical mass | Base mass | Base | MFE mass | $t_R$ | Volume | Quality Score | ppm |
| 20 | 7220.1613 | 329.0525 | A | 7220.1787 | 13.028 | 19619146 | 100 | -2.41 |
| 19 | 6891.1088 | 329.0525 | A | 6891.1344 | 13.094 | 1651958 | 100 | -3.71 |
| 18 | 6562.0563 | 329.0525 | A | 6562.0739 | 12.890 | 44447 | 80 | -2.68 |
| 17 | 6233.0038 | 305.0413 | C | 6233.0093 | 12.525 | 326235 | 80 | -0.88 |
| 16 | 5927.9625 | 305.0413 | C | 5927.9635 | 12.521 | 448797 | 99 | -0.17 |
| 15 | 5622.9212 | 345.0475 | G | 5622.9580 | 12.553 | 728856 | 82 | -6.54 |
| 14 | 5277.8737 | 306.0253 | U | 5277.8785 | 12.443 | 3047682 | 100 | -0.91 |
| 13 | 4971.8484 | 306.0253 | U | 4971.8500 | 12.366 | 3595803 | 100 | -0.32 |
| 12 | 4665.8231 | 329.0525 | A | 4665.8252 | 12.783 | 2318821 | 100 | -0.45 |
| 11 | 4336.7706 | 305.0413 | C | 4336.7689 | 12.336 | 3140948 | 100 | 0.39 |
| 10 | 4031.7293 | 305.0413 | C | 4031.7303 | 12.326 | 3300270 | 93 | -0.25 |
| 9 | 3726.6880 | 329.0525 | A | 3726.6888 | 12.634 | 1909624 | 100 | -0.21 |
| 8 | 3397.6355 | 306.0253 | U | 3397.6349 | 12.168 | 1067451 | 100 | 0.18 |
| 7 | 3091.6102 | 306.0253 | U | 3091.6099 | 12.274 | 1049322 | 100 | 0.10 |
| 6 | 2785.5849 | 329.0525 | A | 2785.5831 | 12.609 | 5644447 | 100 | 0.65 |
| 5 | 2456.5324 | 305.0413 | C | 2456.5286 | 11.834 | 910462 | 100 | 1.55 |
| 4 | 2151.4911 | 306.0253 | U | 2151.4895 | 11.886 | 7177460 | 100 | 0.74 |
| 3 | 1845.4658 | 345.0474 | G | 1845.4658 | 12.247 | 9218649 | 100 | 0.00 |
| 2 | 1500.4184 | 329.0525 | A | 1500.4166 | 13.308 | 7300732 | 100 | 1.20 |
| 1 | 1171.3659 | 345.0475 | G | 1171.3645 | 13.386 | 5051234 | 100 | 1.20 |

**Table S40.** LC/MS analysis of 3´-biotin-labeled RNA #14, showing its mass ladder components.

| Theoretical | | | | Extracted data file after LC/MS analysis | | | | Error |
|---|---|---|---|---|---|---|---|---|
| **Fragments** | **Theoretical mass** | **Base mass** | **Base** | **MFE mass** | **$t_R$** | **Volume** | **Quality Score** | **ppm** |
| 20 | 7234.1769 | 329.0525 | A | 7234.1992 | 13.124 | 107248480 | 100 | -3.08 |
| 19 | 6905.1244 | 329.0525 | A | 6905.1176 | 13.107 | 1877841 | 88 | 0.98 |
| 18 | 6576.0719 | 329.0525 | A | 6576.0803 | 12.921 | 278131 | 80 | -1.28 |
| 17 | 6247.0194 | 305.0413 | C | 6247.0279 | 12.596 | 421971 | 80 | -1.36 |
| 16 | 5941.9781 | 305.0413 | C | 5941.9981 | 12.592 | 4177858 | 100 | -3.37 |
| 15 | 5636.9368 | 345.0475 | G | 5636.9403 | 12.614 | 6652264 | 100 | -0.62 |
| 14 | 5291.8893 | 306.0253 | U | 5291.8944 | 12.514 | 1953738 | 100 | -0.96 |
| 13 | 4985.8640 | 306.0253 | U | 4985.8643 | 12.425 | 662393 | 98 | -0.06 |
| 12 | 4679.8387 | 329.0525 | A | 4679.8388 | 12.837 | 979270 | 100 | -0.02 |
| 11 | 4350.7862 | 305.0413 | C | 4350.7875 | 12.375 | 1975515 | 100 | -0.30 |
| 10 | 4045.7449 | 305.0413 | C | 4045.7450 | 12.384 | 766164 | 100 | -0.02 |
| 9 | 3740.7036 | 329.0525 | A | 3740.7042 | 12.706 | 6380425 | 100 | -0.16 |
| 8 | 3411.6511 | 306.0253 | U | 3411.6469 | 11.987 | 226571 | 100 | 1.23 |
| 7 | 3105.6258 | 306.0253 | U | 3105.6244 | 12.382 | 631119 | 100 | 0.45 |
| 6 | 2799.6005 | 329.0525 | A | 2799.5990 | 12.875 | 7042434 | 100 | 0.54 |
| 5 | 2470.5480 | 319.0570 | $m^5C$ | 2470.5459 | 12.498 | 4315993 | 100 | 0.85 |
| 4 | 2151.4910 | 306.0253 | U | 2151.4895 | 11.886 | 7177460 | 100 | 0.70 |
| 3 | 1845.4657 | 345.0474 | G | 1845.4658 | 12.247 | 9218649 | 100 | -0.05 |
| 2 | 1500.4183 | 329.0525 | A | 1500.4166 | 13.308 | 7300732 | 100 | 1.13 |
| 1 | 1171.3658 | 345.0475 | G | 1171.3645 | 13.386 | 5051234 | 100 | 1.11 |

**Table S41.** Integration of extracted ion current (EIC) of 3´-biotinylated non-modified and $m^5C$ modified RNA samples, respectively, prior to their formic acid degradation.

| RNA samples | EIC Area* | Percent | EIC Area | Percent | EIC Area | Percent | EIC Area | Percent | EIC Area | Percent | EIC Area | Percent |
|---|---|---|---|---|---|---|---|---|---|---|---|---|
| **Biotinylated non-modified** | 26024633 | 88% | 22646082 | 80% | 18965962 | 72% | 18599161 | 63% | 13587754 | 54% | 273207 | 1% |
| **Biotinylated $m^5C$ modified** | 3646078 | 12% | 5571600 | 20% | 7306014 | 28% | 10712696 | 37% | 11749320 | 46% | 24610700 | 99% |

***** Based on the relative quantification method published in 2013 (1).

### We processed the data by algorithm as per the following:

**Fig. 1b**: Maximum plotting window RT set to 15: ax.set_ylim(min_time, 15)
Maximum Mass < 7000
Take the top 500 by Volume (above and including 3486)

**Fig. 1c**: plotting window RT set to between 5.5 and 12: ax.set_ylim(5.5,12)
Maximum Mass < 7500
Take the top 500 by volume (above and including 1219)

**Fig. 2b**: Maximum Mass < 7000
Take the top 1000 by volume (above and including 33693)

**Fig. 4a**: Maximum Mass < 8000
Take the top 500 by volume (above and including 241698)

**Fig. 4b**: Maximum Mass <8000
Take the top 1000 by volume (above and including 63110)

**Fig. S2**: Maximum Mass <8000
Take the top 300 by volume (above and including 121230)

The second step was to analyze the LC/MS data and automatically recognize the RNA sequences. We utilized a modified version of the published algorithm (2).

We made the following modifications to the default.cfg file:

**BEFORE**

```
-WALK_STEPS_MIN_DRAFT, 10          # draft minimum number of steps per walk (before orientation determination)

-WALK_STEPS_MIN_FINAL, 14          # final minimum number of steps per walk

-WALK_PPM, 5.0                     # maximum allowable mass error (ppm) during walk traversal
```

**AFTER**

```
+WALK_STEPS_MIN_DRAFT, 5           # draft minimum number of steps per walk (before orientation determination)

+WALK_STEPS_MIN_FINAL, 8           # final minimum number of steps per walk

+WALK_PPM, 10.0                    # maximum allowable mass error (ppm) during walk traversal
```

We then made the following changes to the core.py file:

**1)** we deleted the requirement of a strictly monotonically increasing or decreasing sequence plot

Commented out:

```
#if len(nextpos) and nextposcall:
        #if not tdir and not bidirectional:
          # get the RT direction of the first step (up or down) and use this for the remainder of the walk
         #tdir = int(np.sign(nextpos['RT'] - pos[-1]['RT']))
```

```
                    # FOR TESTING: calculate tdir as slope
                    # tdir = (nextpos['RT'] - pos[-1]['RT'])/(nextpos['Mass'] - pos[-1]['Mass'])
```

**2)** we disabled a mass filtering step:

```
    # apply the selected filters
    #if PARAMS_['FILTER_MIN_MASS'] is not None:
        #cdb.filter(mass=(PARAMS_['FILTER_MIN_MASS'], startingpos['Mass']))
```

**3)** For **Figs. 1c** and later, the following regions of the code were commented out to display the data without sequencing.

```
##        for c in cpd:
##            for ft, i in zip(trials, range(len(trials))):
##                if (c in [f['Cpd'] for f in ft]) and (c not in top_c):
##                    p = [x for x in compounds if x['Cpd'] == c][0]
##                    if orientations[i]:
##                        top_m.append(p['Mass'])
##                        top_v.append(p['Vol'])
##                        top_t.append(p['RT'])
##                        top_c.append(p['Cpd'])
##                    else:
##                        bottom_m.append(p['Mass'])
##                        bottom_v.append(p['Vol'])
##                        bottom_t.append(p['RT'])
##                        bottom_c.append(p['Cpd'])
        if len(bottom_m):
            p4 = plt.scatter(bottom_m, bottom_t, c=bottom_v, s=msize, edgecolor='k', linewidth=1, marker='o',
```

```
                    alpha=alphahigh, cmap=cmap, norm=norm, zorder=3)
        cbar = fig.colorbar(p4)

    if len(top_m):
        p3 = plt.scatter(top_m, top_t, c=top_v, s=msize, edgecolor='k', linewidth=1, marker='s',
                    alpha=alphahigh, cmap=cmap, norm=norm, zorder=3)
        if 'cbar' in locals():
            cbar.vmin = np.min(np.hstack((top_v, bottom_v)))
            cbar.vmax = np.max(np.hstack((top_v, bottom_v)))
        else:
            cbar = fig.colorbar(p3)

##      #plot trial walks
##      for i in range(len(trials)):
##          if orientations[i]:
##              plt.plot([f['Mass'] for f in trials[i]], [f['RT'] for f in trials[i]], 'k-',
##                      alpha=alphahigh, linewidth=1, zorder=2)
##          else:
##              plt.plot([f['Mass'] for f in trials[i]], [f['RT'] for f in trials[i]], 'k-',
##                      alpha=alphahigh, linewidth=1, zorder=2)
##
##      if plot_labels:
##          ann1 = []
##          ann0 = []
##          for trial, orientation in zip(trials, orientations):
##              for i in range(len(trial) - 1):
##                  if orientation:
##                      a = {'text': baselist.findnamebyid(trial[i + 1]['Call']),
##                           'xy': (trial[i]['Mass'], trial[i]['RT']),
##                           'xytext': (trial[i]['Mass'] / 2 + trial[i + 1]['Mass'] / 2 + annotation_offset[0],
##                                      trial[i]['RT'] + annotation_offset[1]), 'color': trial[i + 1]['WalkScore']}
##                      if a not in ann1:
##                          a = dodgetext(a,ann1,-1)
```

```
##                          if a is not None:
##                              ann1.append(a)
##                  else:
##                      a = {'text': baselist.findnamebyid(trial[i + 1]['Call']),
##                           'xy': (trial[i]['Mass'], trial[i]['RT']),
##                           'xytext': (trial[i]['Mass'] / 2 + trial[i + 1]['Mass'] / 2 - annotation_offset[0],
##                                      trial[i]['RT'] - annotation_offset[1]), 'color': trial[i + 1]['WalkScore']}
##                      if a not in ann0:
##                          a = dodgetext(a,ann0,1)
##                          if a is not None:
##                              ann0.append(a)
##          ann = []
##          for a in chain(ann0, ann1):
##              ann.append(ax.annotate(a['text'], a['xy'], horizontalalignment='center', verticalalignment='center',
##                                     textcoords='data', xytext=a['xytext'],
##                                     arrowprops=dict(arrowstyle="-", color='#999999',
##                                                     alpha=alphalow,
##                                                     connectionstyle="angle,angleA=0,angleB=90,rad=0"),
##                                     color='k'))
##
##    elif len(trials):
##        p1 = plt.scatter(m, t, c=v, s=msize, linewidth=0, alpha=alphahigh, cmap=cmap, norm=norm,
##                         zorder=1)
##        # plot trial walks
##        for i in range(len(trials)):
##            plt.plot([f['Mass'] for f in trials[i]], [f['RT'] for f in trials[i]], 'k-',
##                     alpha=alphahigh, linewidth=1, zorder=2)
##    else:
##        p1 = plt.scatter(m, t, c=v, s=msize, linewidth=0, alpha=alphahigh, cmap=cmap, norm=norm, zorder=1)
##
    if 'cbar' not in locals():
        cbar = fig.colorbar(p1)
##
```

```
##    if plot_midline and len(midline):
##        p2 = plt.plot(midline[:, 0], midline[:, 1], 'k-.', zorder=1)
```

We also flipped the plotting direction for **Fig. 1b** from the original algorithm (changes in bold and highlighted):

```
    if plot_labels:
        ann1 = []
        ann0 = []
        for trial, orientation in zip(trials, orientations):
            for i in range(len(trial) - 1):
                if orientation:
                    a = {'text': baselist.findnamebyid(trial[i + 1]['Call']),
                         'xy': (trial[i]['Mass'], trial[i]['RT']),
                         'xytext': (trial[i]['Mass'] / 2 + trial[i + 1]['Mass'] / 2 + annotation_offset[0],
                                    trial[i]['RT'] + annotation_offset[1]), 'color': trial[i + 1]['WalkScore']}
                    if a not in ann1:
                        a = dodgetext(a,ann1,-1)
                        if a is not None:
                            ann1.append(a)
                else:
                    a = {'text': baselist.findnamebyid(trial[i + 1]['Call']),
```

```
            'xy': (trial[i]['Mass'], trial[i]['RT']),
            'xytext': (trial[i]['Mass'] / 2 + trial[i + 1]['Mass'] / 2 - annotation_offset[0],
                       trial[i]['RT'] - annotation_offset[1]), 'color': trial[i + 1]['WalkScore']}
        if a not in ann0:
            a = dodgetext(a,ann0,1)
            if a is not None:
                ann0.append(a)
```
